## Supplementary material for "RNA covariation at helix-level resolution for the identification of evolutionarily conserved RNA structure": Figure_3.pdf

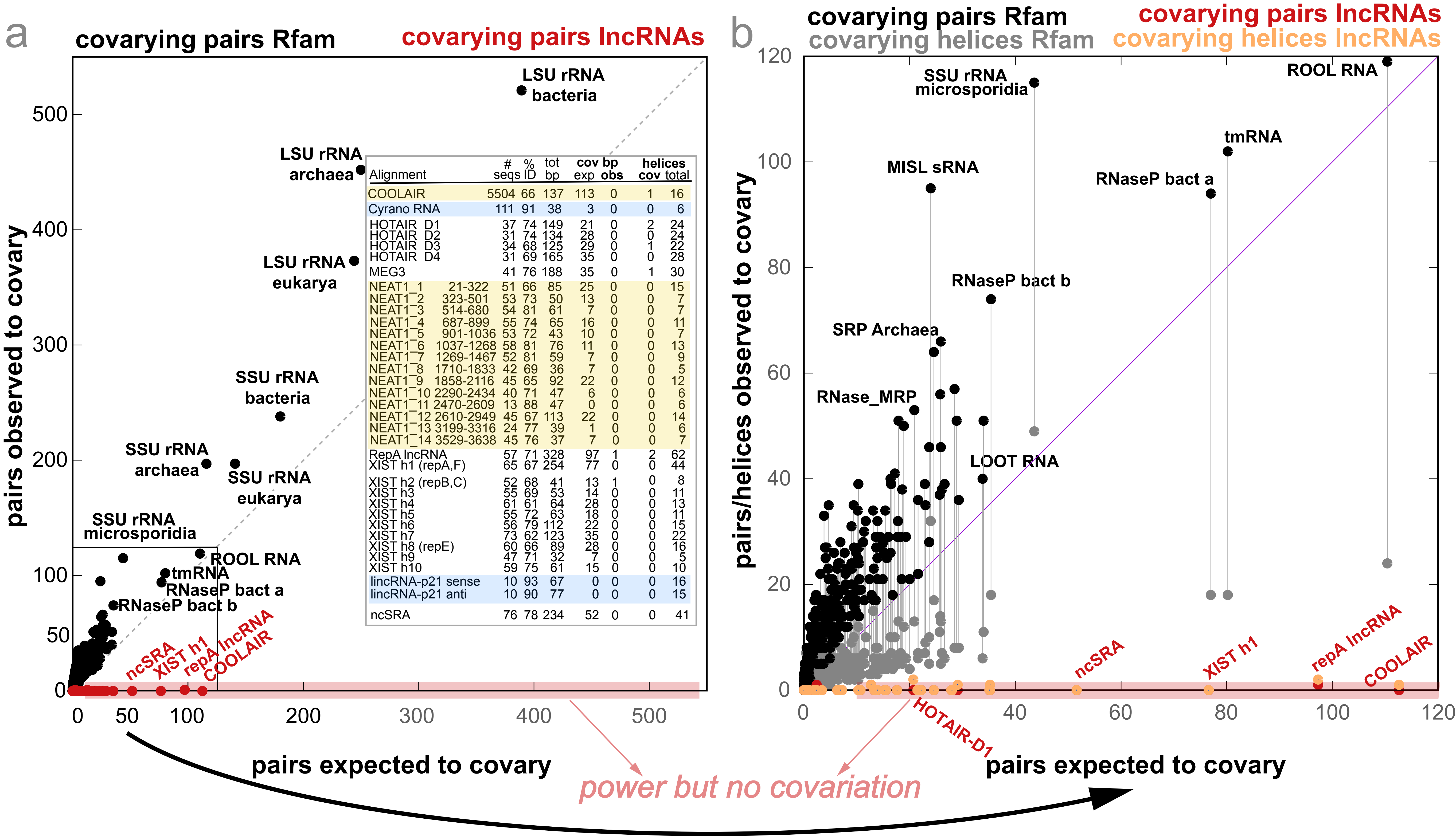


| Alignment | # seqs | % ID | tot bp | cov bp exp | cov bp obs | helices cov | helices total |
|---|---|---|---|---|---|---|---|
| COOLAIR | 5504 | 66 | 137 | 113 | 0 | 1 | 16 |
| Cyrano RNA | 111 | 91 | 38 | 3 | 0 | 0 | 6 |
| HOTAIR D1 | 37 | 74 | 149 | 21 | 0 | 2 | 24 |
| HOTAIR D2 | 31 | 74 | 134 | 28 | 0 | 0 | 24 |
| HOTAIR D3 | 34 | 68 | 125 | 29 | 0 | 1 | 22 |
| HOTAIR D4 | 31 | 69 | 165 | 35 | 0 | 0 | 28 |
| MEG3 | 41 | 76 | 188 | 35 | 0 | 1 | 30 |
| NEAT1_1 21-322 | 51 | 66 | 85 | 25 | 0 | 0 | 15 |
| NEAT1_2 323-501 | 53 | 73 | 50 | 13 | 0 | 0 | 7 |
| NEAT1_3 514-680 | 54 | 81 | 61 | 7 | 0 | 0 | 7 |
| NEAT1_4 687-899 | 55 | 74 | 65 | 16 | 0 | 0 | 11 |
| NEAT1_5 901-1036 | 53 | 72 | 43 | 10 | 0 | 0 | 7 |
| NEAT1_6 1037-1268 | 58 | 81 | 76 | 11 | 0 | 0 | 13 |
| NEAT1_7 1269-1467 | 52 | 81 | 59 | 7 | 0 | 0 | 9 |
| NEAT1_8 1710-1833 | 42 | 69 | 36 | 7 | 0 | 0 | 5 |
| NEAT1_9 1858-2116 | 45 | 65 | 92 | 22 | 0 | 0 | 12 |
| NEAT1_10 2290-2434 | 40 | 71 | 47 | 6 | 0 | 0 | 6 |
| NEAT1_11 2470-2609 | 13 | 88 | 47 | 0 | 0 | 0 | 6 |
| NEAT1_12 2610-2949 | 45 | 67 | 113 | 22 | 0 | 0 | 14 |
| NEAT1_13 3199-3316 | 24 | 77 | 39 | 1 | 0 | 0 | 6 |
| NEAT1_14 3529-3638 | 45 | 76 | 37 | 7 | 0 | 0 | 7 |
| RepA lncRNA | 57 | 71 | 328 | 97 | 1 | 2 | 62 |
| XIST h1 (repA,F) | 65 | 67 | 254 | 77 | 0 | 0 | 44 |
| XIST h2 (repB,C) | 52 | 68 | 41 | 13 | 1 | 0 | 8 |
| XIST h3 | 55 | 69 | 53 | 14 | 0 | 0 | 11 |
| XIST h4 | 61 | 61 | 64 | 28 | 0 | 0 | 13 |
| XIST h5 | 55 | 72 | 63 | 18 | 0 | 0 | 11 |
| XIST h6 | 56 | 79 | 112 | 22 | 0 | 0 | 15 |
| XIST h7 | 73 | 62 | 123 | 35 | 0 | 0 | 22 |
| XIST h8 (repE) | 60 | 66 | 89 | 28 | 0 | 0 | 16 |
| XIST h9 | 47 | 71 | 32 | 7 | 0 | 0 | 5 |
| XIST h10 | 59 | 75 | 61 | 15 | 0 | 0 | 10 |
| lincRNA-p21 sense | 10 | 93 | 67 | 0 | 0 | 0 | 16 |
| lincRNA-p21 anti | 10 | 90 | 77 | 0 | 0 | 0 | 15 |
| ncSRA | 76 | 78 | 234 | 52 | 0 | 0 | 41 |
