## Supplementary material for "RNA covariation at helix-level resolution for the identification of evolutionarily conserved RNA structure": Figure_4.pdf

### NEAT1 323-501

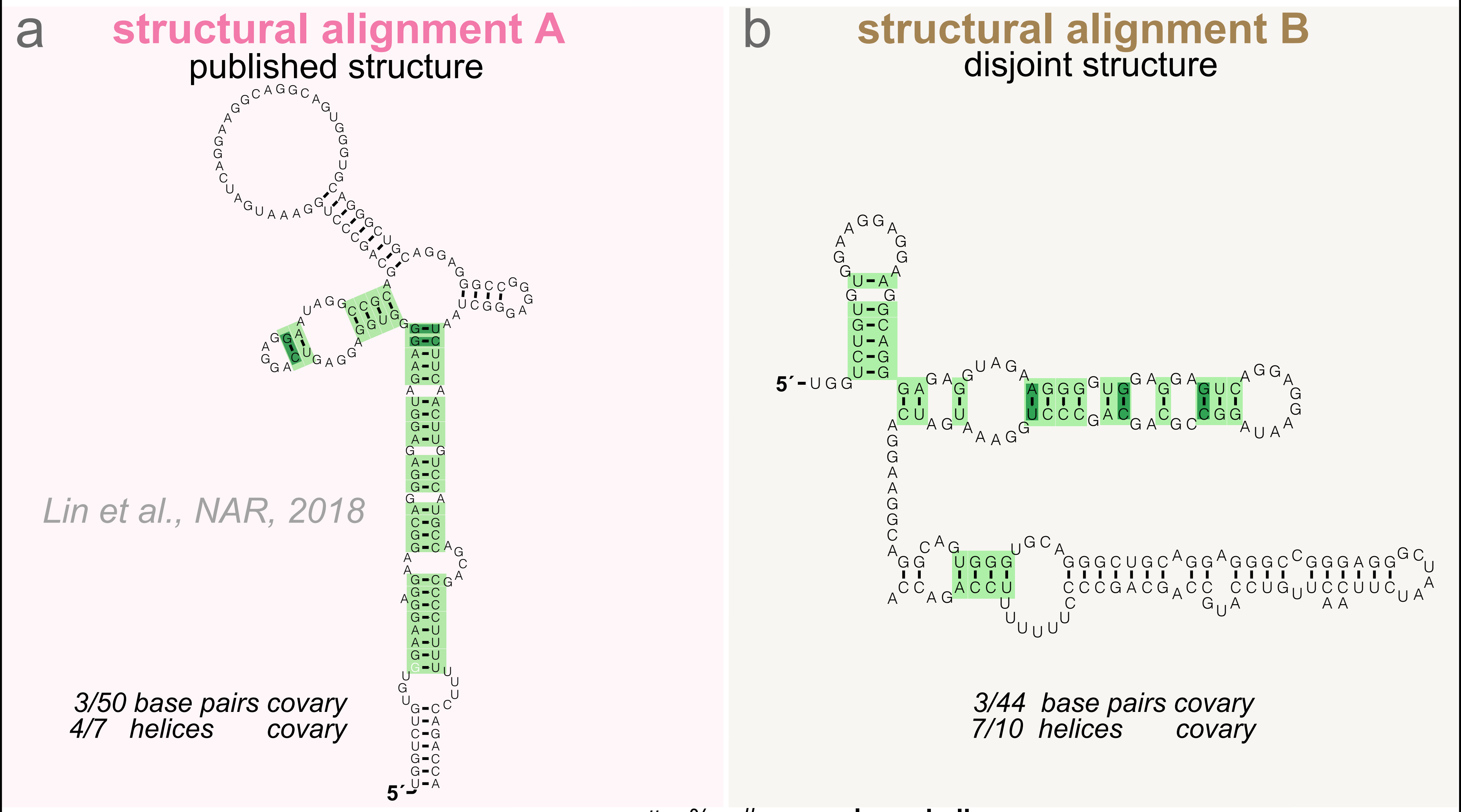


c

| Alignment | # seqs | % ID | # bp | cov bp exp | cov bp obs | helices cov | helices total |
|---|---|---|---|---|---|---|---|
| **nhmmer** | 53 | 73 | 50 | **13** | **0** | **0** | **7** |
| **muscle** | 54 | 66 | 50 | **19** | **0** | **0** | **7** |
| structural A | 53 | 66 | 50 | 18 | 3 | 4 | 7 |
| structural B | 53 | 66 | 44 | 17 | 3 | 7 | 10 |

### COOLAIR IIi

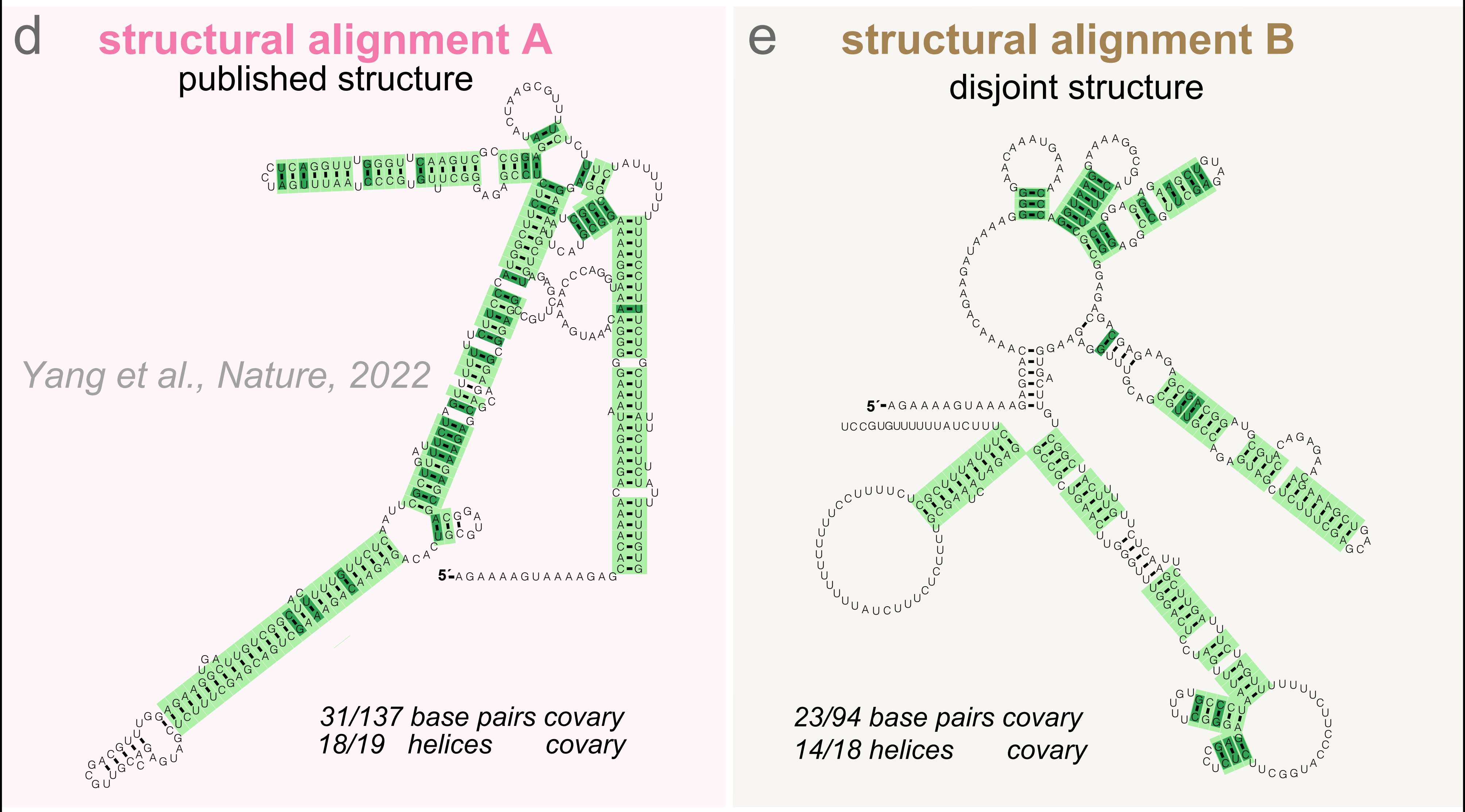


f

| Alignment | # seqs | % ID | # bp | cov bp exp | cov bp obs | helices cov | helices total |
|---|---|---|---|---|---|---|---|
| **nhmmer** | 5504 | 66 | 137 | **113** | **0** | **1** | **16** |
| **muscle** | 5497 | 64 | 137 | **118** | **0** | **1** | **17** |
| structural A | 5497 | 60 | 137 | 125 | 31 | 18 | 19 |
| structural B | 5497 | 61 | 94 | 92 | 23 | 14 | 18 |
