## Supplementary material for "RNA covariation at helix-level resolution for the identification of evolutionarily conserved RNA structure": compare.pl

```
#!/usr/bin/perl -w
# 
# Select Rfam families
#
use strict;
use Class::Struct;
use POSIX;
    
use constant GNUPLOT => '/usr/local/bin/gnuplot';
use lib '/Users/erivas/src/Mysrc/R-scape/scripts';
use FUNCS;

use vars qw ($opt_v);  # required if strict used
use Getopt::Std;
getopts ('v');

# Print a helpful message if the user provides no input file.
if (@ARGV) {
    print "usage:  Rfam_select.pl [options] <R-scape.seed> \n\n";
    print "options:\n";
    print "-v    :  be verbose\n";
    exit;
}

my $file1 = "data/Rfam14.8/select_R-scape_null/select_null.sto.list";
my $file2 = "data/Rfam14.8/select/select.sto.list";

my $n1 = 0;
my @list1;
open (F, "$file1") || die;
while(<F>) {
    if (/(RF\d+)/) { $list1[$n1++] = $1; }
}
close(F);

my $n2 = 0;
my @list2;
open (F, "$file2") || die;
while(<F>) {
    if (/(RF\d+)/) { $list2[$n2++] = $1; }
}
close(F);

my $nfound = 0;
my @found;
if ($n1 >= $n2) {
    for (my $n = 0; $n < $n1; $n ++) {
	my $this = $list1[$n];
	$found[$n] = 0;
	
	for (my $m = 0; $m < $n2; $m ++) {
	    if ($this =~ /^$list2[$m]$/) { $nfound ++; $found[$n] = 1; print "$list1[$n] found at $m\n"; last; }
	}
    }

    for (my $n = 0; $n < $n1; $n ++) {
	if ($found[$n] == 0) { print "$list1[$n] not found\n"; }
    }
    
    print "found $nfound/$n1\n";

}
```
