## Supplementary material for "RNA covariation at helix-level resolution for the identification of evolutionarily conserved RNA structure": profmark.pl

```
#!/usr/bin/perl -w
# 
# profmark
#
use strict;
use Class::Struct;
use POSIX;
    
#use constant GNUPLOT => '/usr/local/bin/gnuplot';
use constant GNUPLOT => '/n/helmod/apps/centos7/Core/gnuplot/5.2.6-fasrc01/bin/gnuplot';

#use lib '/Users/erivas/src/Mysrc/R-scape/scripts';
use lib '/n/eddy_lab/users/erivas/rscape_v2.0.0.p/scripts';
use FUNCS;

use vars qw ($opt_v $opt_R $opt_F);  # required if strict used
use Getopt::Std;
getopts ('FNRv');

# Print a helpful message if the user provides no input file.
if (!@ARGV) {
    print "usage:  profmark.pl [options] <Rfam> <maindir> <rscapedir> <family> \n\n";
    print "options:\n";
    print "-v    :  be verbose\n";
    exit;
}

my $Rfam      = shift;
my $maindir   = shift;
my $rscapedir = shift;
my $family    = shift;

my $do_rscape = 0;
if ($opt_R) { $do_rscape = 1; }

my $do_flanking = 0;
if ($opt_F) { $do_flanking = 1; }

my $nmsa = 20; # number of decoys per msa and decoy method
if ($family =~ /RF00177/ || $family =~ /RF01959/ || $family =~ /RF01960/ || $family =~ /RF02542/) { $nmsa = 2; } # SSU families
if ($family =~ /RF02540/ || $family =~ /RF02541/ || $family =~ /RF02543/)                         { $nmsa = 1; } # LSU families

my $natbl = 3;
my @atbl;
$atbl[0] = 0.2;
$atbl[1] = 0.4;
$atbl[2] = 0.5;

my $rscape     = "$rscapedir/bin/R-scape";
my $rscape_sim = "$rscapedir/bin/R-scape-sim";

my $profmark_dir         = "$maindir/data/$Rfam/profmark";
my $select_dir           = "$profmark_dir/select";
my $selectnull_dir       = "$profmark_dir/select_R-scape_null";
my $synthetic_dir        = "$profmark_dir/select_synthetic";
my $syntheticnull_dir    = "$profmark_dir/select_synthetic_null";

my $rfam_sto             = "$select_dir/$family.seed.sto";
my $rfamnull_sto         = "$selectnull_dir/$family.null.sto";
my $synthetic_sto        = "$synthetic_dir/$family.synthetic.sto";
my $syntheticnull_sto    = "$syntheticnull_dir/$family.synthetic.null.sto";
if ($do_rscape) {
    system("rm $selectnull_dir/$family.*\n");
    system("rm $synthetic_dir/$family.*\n");
    system("rm $syntheticnull_dir/$family.*\n");
}

# main outputs
my $results_dir                              = "$profmark_dir/results";
my $profmark_select_cov_out                  = "$results_dir/$family.select.cov.out";
my $profmark_select_hlx_fisher_out           = "$results_dir/$family.select.hlx.fisher.out";
my $profmark_select_hlx_sidak_out            = "$results_dir/$family.select.hlx.sidak.out";
my $profmark_select_hlx_lancaster_out        = "$results_dir/$family.select.hlx.lancaster.out";
my $profmark_select_hlx_wfisher_out          = "$results_dir/$family.select.hlx.wfisher.out";
my $profmark_select_hlx_lancaster_join_out   = "$results_dir/$family.select.hlx.lancaster_join.out";
my $profmark_select_hlx_wfisher_join_out     = "$results_dir/$family.select.hlx.wfisher_join.out";
my $profmark_select_hlx_lancaster_double_out = "$results_dir/$family.select.hlx.lancaster_double.out";
my $profmark_select_hlx_wfisher_double_out   = "$results_dir/$family.select.hlx.wfisher_double.out";
system("rm $profmark_select_cov_out\n");
system("rm $profmark_select_hlx_fisher_out\n");
system("rm $profmark_select_hlx_sidak_out\n");
system("rm $profmark_select_hlx_lancaster_out\n");
system("rm $profmark_select_hlx_wfisher_out\n");
system("rm $profmark_select_hlx_lancaster_join_out\n");
system("rm $profmark_select_hlx_wfisher_join_out\n");
system("rm $profmark_select_hlx_lancaster_double_out\n");
system("rm $profmark_select_hlx_wfisher_double_out\n");

my $profmark_select_noss_cov_out                  = "$results_dir/$family.select.noss.cov.out";
my $profmark_select_noss_hlx_fisher_out           = "$results_dir/$family.select.noss.hlx.fisher.out";
my $profmark_select_noss_hlx_sidak_out            = "$results_dir/$family.select.noss.hlx.sidak.out";
my $profmark_select_noss_hlx_lancaster_out        = "$results_dir/$family.select.noss.hlx.lancaster.out";
my $profmark_select_noss_hlx_wfisher_out          = "$results_dir/$family.select.noss.hlx.wfisher.out";
my $profmark_select_noss_hlx_lancaster_join_out   = "$results_dir/$family.select.noss.hlx.lancaster_join.out";
my $profmark_select_noss_hlx_wfisher_join_out     = "$results_dir/$family.select.noss.hlx.wfisher_join.out";
my $profmark_select_noss_hlx_lancaster_double_out = "$results_dir/$family.select.noss.hlx.lancaster_double.out";
my $profmark_select_noss_hlx_wfisher_double_out   = "$results_dir/$family.select.noss.hlx.wfisher_double.out";
system("rm $profmark_select_noss_cov_out\n");
system("rm $profmark_select_noss_hlx_fisher_out\n");
system("rm $profmark_select_noss_hlx_sidak_out\n");
system("rm $profmark_select_noss_hlx_lancaster_out\n");
system("rm $profmark_select_noss_hlx_wfisher_out\n");
system("rm $profmark_select_noss_hlx_lancaster_join_out\n");
system("rm $profmark_select_noss_hlx_wfisher_join_out\n");
system("rm $profmark_select_noss_hlx_lancaster_double_out\n");
system("rm $profmark_select_noss_hlx_wfisher_double_out\n");

my $profmark_synthetic_cov_out                  = "$results_dir/$family.synthetic.cov.out";
my $profmark_synthetic_hlx_fisher_out           = "$results_dir/$family.synthetic.hlx.fisher.out";
my $profmark_synthetic_hlx_sidak_out            = "$results_dir/$family.synthetic.hlx.sidak.out";
my $profmark_synthetic_hlx_lancaster_out        = "$results_dir/$family.synthetic.hlx.lancaster.out";
my $profmark_synthetic_hlx_wfisher_out          = "$results_dir/$family.synthetic.hlx.wfisher.out";
my $profmark_synthetic_hlx_lancaster_join_out   = "$results_dir/$family.synthetic.hlx.lancaster_join.out";
my $profmark_synthetic_hlx_wfisher_join_out     = "$results_dir/$family.synthetic.hlx.wfisher_join.out";
my $profmark_synthetic_hlx_lancaster_double_out = "$results_dir/$family.synthetic.hlx.lancaster_double.out";
my $profmark_synthetic_hlx_wfisher_double_out   = "$results_dir/$family.synthetic.hlx.wfisher_double.out";
system("rm $profmark_synthetic_cov_out\n");
system("rm $profmark_synthetic_hlx_fisher_out\n");
system("rm $profmark_synthetic_hlx_sidak_out\n");
system("rm $profmark_synthetic_hlx_lancaster_out\n");
system("rm $profmark_synthetic_hlx_wfisher_out\n");
system("rm $profmark_synthetic_hlx_lancaster_join_out\n");
system("rm $profmark_synthetic_hlx_wfisher_join_out\n");
system("rm $profmark_synthetic_hlx_lancaster_double_out\n");
system("rm $profmark_synthetic_hlx_wfisher_double_out\n");

my $profmark_synthetic_noss_cov_out                  = "$results_dir/$family.synthetic.noss.cov.out";
my $profmark_synthetic_noss_hlx_fisher_out           = "$results_dir/$family.synthetic.noss.hlx.fisher.out";
my $profmark_synthetic_noss_hlx_sidak_out            = "$results_dir/$family.synthetic.noss.hlx.sidak.out";
my $profmark_synthetic_noss_hlx_lancaster_out        = "$results_dir/$family.synthetic.noss.hlx.lancaster.out";
my $profmark_synthetic_noss_hlx_wfisher_out          = "$results_dir/$family.synthetic.noss.hlx.wfisher.out";
my $profmark_synthetic_noss_hlx_lancaster_join_out   = "$results_dir/$family.synthetic.noss.hlx.lancaster_join.out";
my $profmark_synthetic_noss_hlx_wfisher_join_out     = "$results_dir/$family.synthetic.noss.hlx.wfisher_join.out";
my $profmark_synthetic_noss_hlx_lancaster_double_out = "$results_dir/$family.synthetic.noss.hlx.lancaster_double.out";
my $profmark_synthetic_noss_hlx_wfisher_double_out   = "$results_dir/$family.synthetic.noss.hlx.wfisher_double.out";
system("rm $profmark_synthetic_noss_cov_out\n");
system("rm $profmark_synthetic_noss_hlx_fisher_out\n");
system("rm $profmark_synthetic_noss_hlx_sidak_out\n");
system("rm $profmark_synthetic_noss_hlx_lancaster_out\n");
system("rm $profmark_synthetic_noss_hlx_wfisher_out\n");
system("rm $profmark_synthetic_noss_hlx_lancaster_join_out\n");
system("rm $profmark_synthetic_noss_hlx_wfisher_join_out\n");
system("rm $profmark_synthetic_noss_hlx_lancaster_double_out\n");
system("rm $profmark_synthetic_noss_hlx_wfisher_double_out\n");

my $fisher           = "FISHER";    
my $sidak            = "SIDAK";    
my $lancaster        = "LANCASTER";    
my $wfisher          = "WFISHER";    
my $lancaster_join   = "LANCASTER_JOIN";
my $wfisher_join     = "WFISHER_JOIN";    
my $lancaster_double = "LANCASTER_DOUBLE";    
my $wfisher_double   = "WFISHER_DOUBLE";    

my $select_true_bp;
my $select_true_hlx;
my $synthetic_true_bp;
my $synthetic_true_hlx;


# R-scape with --outnull to save null alignments
#----------------------------------------------------------------------------------------------------------------------------------
my $synthetic = 0;
my $decoy     = 0;
my $twotest;
my $cmd;

if ($do_rscape) {
    $cmd = "$rscape -s --seed 0 -E 1001 --scmin -10000 --hpts 20000  --outnull --profmark --fisher --sidak --lancaster --wfisher  --outdir $select_dir/R-scape      --nofigures  $rfam_sto";
    print "$cmd\n";
    system("$cmd\n");

    $cmd = "$rscape    --seed 0 -E 1001 --scmin -10000 --hpts 20000  --outnull --profmark --fisher --sidak --lancaster --wfisher  --outdir $select_dir/R-scape_noss --nofigures  $rfam_sto";
    print "$cmd\n";
    system("$cmd\n");
}
$twotest = 1;
add_cov_file($profmark_select_cov_out,                $family, "$select_dir/R-scape",      $synthetic, $decoy, $twotest,   \$select_true_bp);
add_hlx_file($profmark_select_hlx_fisher_out,         $family, "$select_dir/R-scape",      $synthetic, $decoy, $fisher,    \$select_true_hlx);
add_hlx_file($profmark_select_hlx_sidak_out,          $family, "$select_dir/R-scape",      $synthetic, $decoy, $sidak,     \$select_true_hlx);
add_hlx_file($profmark_select_hlx_lancaster_out,      $family, "$select_dir/R-scape",      $synthetic, $decoy, $lancaster, \$select_true_hlx);
add_hlx_file($profmark_select_hlx_wfisher_out,        $family, "$select_dir/R-scape",      $synthetic, $decoy, $wfisher,   \$select_true_hlx);

$twotest = 0;
add_cov_file($profmark_select_noss_cov_out,           $family, "$select_dir/R-scape_noss", $synthetic, $decoy, $twotest,   \$select_true_bp);
add_hlx_file($profmark_select_noss_hlx_fisher_out,    $family, "$select_dir/R-scape_noss", $synthetic, $decoy, $fisher,    \$select_true_hlx);
add_hlx_file($profmark_select_noss_hlx_sidak_out,     $family, "$select_dir/R-scape_noss", $synthetic, $decoy, $sidak,     \$select_true_hlx);
add_hlx_file($profmark_select_noss_hlx_lancaster_out, $family, "$select_dir/R-scape_noss", $synthetic, $decoy, $lancaster, \$select_true_hlx);
add_hlx_file($profmark_select_noss_hlx_wfisher_out,   $family, "$select_dir/R-scape_noss", $synthetic, $decoy, $wfisher,   \$select_true_hlx);

if ($do_rscape) {
    $cmd = "$rscape -s            --scmin -10000 --hpts 20000 --lancaster_join   --wfisher_join   --joinsubs   --outdir $select_dir/R-scape_join      --nofigures   $rfam_sto";
    print "$cmd\n";
    system("$cmd\n");
    
    $cmd = "$rscape               --scmin -10000 --hpts 20000 --lancaster_join   --wfisher_join   --joinsubs   --outdir $select_dir/R-scape_noss_join --nofigures   $rfam_sto";
    print "$cmd\n";
    system("$cmd\n");
}
add_hlx_file($profmark_select_hlx_lancaster_join_out,      $family, "$select_dir/R-scape_join",      $synthetic, $decoy, $lancaster_join, \$select_true_hlx);
add_hlx_file($profmark_select_hlx_wfisher_join_out,        $family, "$select_dir/R-scape_join",      $synthetic, $decoy, $wfisher_join,   \$select_true_hlx);

add_hlx_file($profmark_select_noss_hlx_lancaster_join_out, $family, "$select_dir/R-scape_noss_join", $synthetic, $decoy, $lancaster_join, \$select_true_hlx);
add_hlx_file($profmark_select_noss_hlx_wfisher_join_out,   $family, "$select_dir/R-scape_noss_join", $synthetic, $decoy, $wfisher_join,   \$select_true_hlx);

if ($do_rscape){
    $cmd = "$rscape -s            --scmin -10000 --hpts 20000 --lancaster_double --wfisher_double --doublesubs --outdir $select_dir/R-scape_double      --nofigures $rfam_sto";
    print "$cmd\n";
    system("$cmd\n");
    
    $cmd = "$rscape               --scmin -10000 --hpts 20000 --lancaster_double --wfisher_double --doublesubs --outdir $select_dir/R-scape_noss_double --nofigures $rfam_sto";
    print "$cmd\n";
    system("$cmd\n");
}
add_hlx_file($profmark_select_hlx_lancaster_double_out,      $family, "$select_dir/R-scape_double",      $synthetic, $decoy, $lancaster_double,  \$select_true_hlx);
add_hlx_file($profmark_select_hlx_wfisher_double_out,        $family, "$select_dir/R-scape_double",      $synthetic, $decoy, $wfisher_double,    \$select_true_hlx);

add_hlx_file($profmark_select_noss_hlx_lancaster_double_out, $family, "$select_dir/R-scape_noss_double", $synthetic, $decoy, $lancaster_double,  \$select_true_hlx);
add_hlx_file($profmark_select_noss_hlx_wfisher_double_out,   $family, "$select_dir/R-scape_noss_double", $synthetic, $decoy, $wfisher_double,    \$select_true_hlx);


# copy the null alignments to their own directory select_R-scape_null
#-------------------------------------------------------------------------------------------------------------------------------------
$cmd = "mv $select_dir/R-scape/$family*.null.sto $rfamnull_sto";
print "$cmd\n";
system("$cmd\n");

$synthetic = 0;
$decoy     = 1;
if ($do_rscape) {
    $cmd = "$rscape -s -E 1001 --scmin -100000  --hpts 20000 --fisher --sidak --lancaster --wfisher --outdir $selectnull_dir/R-scape      --nofigures       $rfamnull_sto";
    print "$cmd\n";
    system("$cmd\n");
    $cmd = "$rscape    -E 1001 --scmin -100000  --hpts 20000 --fisher --sidak --lancaster --wfisher --outdir $selectnull_dir/R-scape_noss --nofigures       $rfamnull_sto";
    print "$cmd\n";
    system("$cmd\n");
}    
$twotest = 1;
add_cov_file($profmark_select_cov_out,                $family, "$selectnull_dir/R-scape",      $synthetic, $decoy, $twotest);
add_hlx_file($profmark_select_hlx_fisher_out,         $family, "$selectnull_dir/R-scape",      $synthetic, $decoy, $fisher);
add_hlx_file($profmark_select_hlx_sidak_out,          $family, "$selectnull_dir/R-scape",      $synthetic, $decoy, $sidak);
add_hlx_file($profmark_select_hlx_lancaster_out,      $family, "$selectnull_dir/R-scape",      $synthetic, $decoy, $lancaster);
add_hlx_file($profmark_select_hlx_wfisher_out,        $family, "$selectnull_dir/R-scape",      $synthetic, $decoy, $wfisher);

$twotest = 0;
add_cov_file($profmark_select_noss_cov_out,           $family, "$selectnull_dir/R-scape_noss", $synthetic, $decoy, $twotest);
add_hlx_file($profmark_select_noss_hlx_fisher_out,    $family, "$selectnull_dir/R-scape_noss", $synthetic, $decoy, $fisher);
add_hlx_file($profmark_select_noss_hlx_sidak_out,     $family, "$selectnull_dir/R-scape_noss", $synthetic, $decoy, $sidak);
add_hlx_file($profmark_select_noss_hlx_lancaster_out, $family, "$selectnull_dir/R-scape_noss", $synthetic, $decoy, $lancaster);
add_hlx_file($profmark_select_noss_hlx_wfisher_out,   $family, "$selectnull_dir/R-scape_noss", $synthetic, $decoy, $wfisher);

if ($do_rscape) {
    $cmd = "$rscape -s  --scmin -10000 --hpts 20000 --lancaster_join   --wfisher_join   --joinsubs   --outdir $selectnull_dir/R-scape_join      --nofigures   $rfamnull_sto";
    print "$cmd\n";
    system("$cmd\n");
    $cmd = "$rscape     --scmin -10000 --hpts 20000 --lancaster_join   --wfisher_join   --joinsubs   --outdir $selectnull_dir/R-scape_noss_join --nofigures   $rfamnull_sto";
    print "$cmd\n";
    system("$cmd\n");
}
add_hlx_file($profmark_select_hlx_lancaster_join_out,      $family, "$selectnull_dir/R-scape_join",      $synthetic, $decoy, $lancaster_join);
add_hlx_file($profmark_select_hlx_wfisher_join_out,        $family, "$selectnull_dir/R-scape_join",      $synthetic, $decoy, $wfisher_join);
add_hlx_file($profmark_select_noss_hlx_lancaster_join_out, $family, "$selectnull_dir/R-scape_noss_join", $synthetic, $decoy, $lancaster_join);
add_hlx_file($profmark_select_noss_hlx_wfisher_join_out,   $family, "$selectnull_dir/R-scape_noss_join", $synthetic, $decoy, $wfisher_join);

if ($do_rscape) {
    $cmd = "$rscape -s  --scmin -10000 --hpts 20000 --lancaster_double --wfisher_double --doublesubs --outdir $selectnull_dir/R-scape_double      --nofigures $rfamnull_sto";
    print "$cmd\n";
    system("$cmd\n");
    $cmd = "$rscape     --scmin -10000 --hpts 20000 --lancaster_double --wfisher_double --doublesubs --outdir $selectnull_dir/R-scape_noss_double --nofigures $rfamnull_sto";
    print "$cmd\n";
    system("$cmd\n");
}
add_hlx_file($profmark_select_hlx_lancaster_double_out,      $family, "$selectnull_dir/R-scape_double",      $synthetic, $decoy, $lancaster_double);
add_hlx_file($profmark_select_hlx_wfisher_double_out,        $family, "$selectnull_dir/R-scape_double",      $synthetic, $decoy, $wfisher_double);
add_hlx_file($profmark_select_noss_hlx_lancaster_double_out, $family, "$selectnull_dir/R-scape_noss_double", $synthetic, $decoy, $lancaster_double);
add_hlx_file($profmark_select_noss_hlx_wfisher_double_out,   $family, "$selectnull_dir/R-scape_noss_double", $synthetic, $decoy, $wfisher_double);


# create the synthetic alignments (with ss)
#------------------------------------------------------------------------------------------------------------------------------------------
$synthetic = 1;
$decoy     = 0;

if ($do_rscape) {
    # generate the synthetic alignments
    #
    for (my $a = 0; $a < $nmsa; $a ++) {
	for (my $b = 0; $b < $natbl; $b ++) {
	    my $newsto   = "$synthetic_sto.$a.$b";
	    my $newname  = "$newsto";
	    if ($newname =~ /\/([^\/]+)\s*$/) { $newname = $1; }
	    
	    $cmd = "$rscape_sim --seed 0 --given --rnass --atbl $atbl[$b] --sim_name $newname -o $newsto $rfam_sto";
	    print "$cmd\n";
	    system("$cmd\n");
	    
	    $cmd = "cat $newsto >> $synthetic_sto";
	    print "$cmd\n";
	    system("$cmd\n");
	    system("rm $newsto\n");
	}
    }

    $cmd = "$rscape -s -E 1001 --scmin -100000  --hpts 20000 --fisher --sidak --lancaster --wfisher  --outdir $synthetic_dir/R-scape      --nofigures       $synthetic_sto";
    print "$cmd\n";
    system("$cmd\n");
    
    $cmd = "$rscape    -E 1001 --scmin -100000  --hpts 20000 --fisher --sidak --lancaster --wfisher  --outdir $synthetic_dir/R-scape_noss --nofigures       $synthetic_sto";
    print "$cmd\n";
    system("$cmd\n");
}
$twotest = 1;
add_cov_file($profmark_synthetic_cov_out,                $family, "$synthetic_dir/R-scape", $synthetic, $decoy, $twotest,   \$synthetic_true_bp);
add_hlx_file($profmark_synthetic_hlx_fisher_out,         $family, "$synthetic_dir/R-scape", $synthetic, $decoy, $fisher,    \$synthetic_true_hlx);
add_hlx_file($profmark_synthetic_hlx_sidak_out,          $family, "$synthetic_dir/R-scape", $synthetic, $decoy, $sidak,     \$synthetic_true_hlx);
add_hlx_file($profmark_synthetic_hlx_lancaster_out,      $family, "$synthetic_dir/R-scape", $synthetic, $decoy, $lancaster, \$synthetic_true_hlx);
add_hlx_file($profmark_synthetic_hlx_wfisher_out,        $family, "$synthetic_dir/R-scape", $synthetic, $decoy, $wfisher,   \$synthetic_true_hlx);

$twotest = 0;
add_cov_file($profmark_synthetic_noss_cov_out,           $family, "$synthetic_dir/R-scape_noss", $synthetic, $decoy, $twotest,   \$synthetic_true_bp);
add_hlx_file($profmark_synthetic_noss_hlx_fisher_out,    $family, "$synthetic_dir/R-scape_noss", $synthetic, $decoy, $fisher,    \$synthetic_true_hlx);
add_hlx_file($profmark_synthetic_noss_hlx_sidak_out,     $family, "$synthetic_dir/R-scape_noss", $synthetic, $decoy, $sidak,     \$synthetic_true_hlx);
add_hlx_file($profmark_synthetic_noss_hlx_lancaster_out, $family, "$synthetic_dir/R-scape_noss", $synthetic, $decoy, $lancaster, \$synthetic_true_hlx);
add_hlx_file($profmark_synthetic_noss_hlx_wfisher_out,   $family, "$synthetic_dir/R-scape_noss", $synthetic, $decoy, $wfisher,   \$synthetic_true_hlx);

if ($do_rscape) {
    $cmd = "$rscape -s  --scmin -10000 --hpts 20000 --lancaster_join   --wfisher_join   --joinsubs   --outdir $synthetic_dir/R-scape_join     --nofigures   $synthetic_sto";
    print "$cmd\n";
    system("$cmd\n");
    
    $cmd = "$rscape     --scmin -10000 --hpts 20000 --lancaster_join   --wfisher_join   --joinsubs   --outdir $synthetic_dir/R-scape_noss_join --nofigures   $synthetic_sto";
    print "$cmd\n";
    system("$cmd\n");
}
add_hlx_file($profmark_synthetic_hlx_lancaster_join_out,      $family, "$synthetic_dir/R-scape_join",      $synthetic, $decoy, $lancaster_join, \$synthetic_true_hlx);
add_hlx_file($profmark_synthetic_hlx_wfisher_join_out,        $family, "$synthetic_dir/R-scape_join",      $synthetic, $decoy, $wfisher_join,   \$synthetic_true_hlx);

add_hlx_file($profmark_synthetic_noss_hlx_lancaster_join_out, $family, "$synthetic_dir/R-scape_noss_join", $synthetic, $decoy, $lancaster_join, \$synthetic_true_hlx);
add_hlx_file($profmark_synthetic_noss_hlx_wfisher_join_out,   $family, "$synthetic_dir/R-scape_noss_join", $synthetic, $decoy, $wfisher_join,   \$synthetic_true_hlx);

if ($do_rscape) {
    $cmd = "$rscape -s  --scmin -10000 --hpts 20000 --lancaster_double --wfisher_double --doublesubs --outdir $synthetic_dir/R-scape_double      --nofigures $synthetic_sto";
    print "$cmd\n";
    system("$cmd\n");

    $cmd = "$rscape     --scmin -10000 --hpts 20000 --lancaster_double --wfisher_double --doublesubs --outdir $synthetic_dir/R-scape_noss_double --nofigures $synthetic_sto";
    print "$cmd\n";
    system("$cmd\n");
}
add_hlx_file($profmark_synthetic_hlx_lancaster_double_out,      $family, "$synthetic_dir/R-scape_double",      $synthetic, $decoy, $lancaster_double,  \$synthetic_true_hlx);
add_hlx_file($profmark_synthetic_hlx_wfisher_double_out,        $family, "$synthetic_dir/R-scape_double",      $synthetic, $decoy, $wfisher_double,    \$synthetic_true_hlx);

add_hlx_file($profmark_synthetic_noss_hlx_lancaster_double_out, $family, "$synthetic_dir/R-scape_noss_double", $synthetic, $decoy, $lancaster_double,  \$synthetic_true_hlx);
add_hlx_file($profmark_synthetic_noss_hlx_wfisher_double_out,   $family, "$synthetic_dir/R-scape_noss_double", $synthetic, $decoy, $wfisher_double,    \$synthetic_true_hlx);


# create the synthetic alignments (no ss)
#------------------------------------------------------------------------------------------------------------------------------------------------------------------------------
$synthetic = 1;
$decoy     = 1;

if ($do_rscape) {
    # generate the synthetic alignments
    #
    for (my $a = 0; $a < $nmsa; $a ++) {
	for (my $b = 0; $b < $natbl; $b ++) {
	    my $newsto   = "$syntheticnull_sto.$a.$b";
	    my $newname  = "$newsto";
	    if ($newname =~ /\/([^\/]+)\s*$/) { $newname = $1; }
	    
	    $cmd = "$rscape_sim --seed 0 --given --profmark --atbl $atbl[$b] --sim_name $newname -o $newsto $rfam_sto";
	    print "$cmd\n";
	    system("$cmd\n");
	    
	    $cmd = "cat $newsto >> $syntheticnull_sto";
	    print "$cmd\n";
	    system("$cmd\n");
	    system("rm $newsto\n");
	}
    }
}


if ($do_rscape) {
    $cmd = "$rscape -s -E 1001 --scmin -100000  --hpts 20000 --fisher --sidak --lancaster --wfisher  --outdir $syntheticnull_dir/R-scape      --nofigures   $syntheticnull_sto";
    print "$cmd\n";
    system("$cmd\n");
    $cmd = "$rscape    -E 1001 --scmin -100000  --hpts 20000 --fisher --sidak --lancaster --wfisher  --outdir $syntheticnull_dir/R-scape_noss --nofigures   $syntheticnull_sto";
    print "$cmd\n";
    system("$cmd\n");
}
$twotest = 1;
add_cov_file($profmark_synthetic_cov_out,                $family, "$syntheticnull_dir/R-scape",      $synthetic, $decoy, $twotest);
add_hlx_file($profmark_synthetic_hlx_fisher_out,         $family, "$syntheticnull_dir/R-scape",      $synthetic, $decoy, $fisher);
add_hlx_file($profmark_synthetic_hlx_sidak_out,          $family, "$syntheticnull_dir/R-scape",      $synthetic, $decoy, $sidak);
add_hlx_file($profmark_synthetic_hlx_lancaster_out,      $family, "$syntheticnull_dir/R-scape",      $synthetic, $decoy, $lancaster);
add_hlx_file($profmark_synthetic_hlx_wfisher_out,        $family, "$syntheticnull_dir/R-scape",      $synthetic, $decoy, $wfisher);

$twotest = 0;
add_cov_file($profmark_synthetic_noss_cov_out,           $family, "$syntheticnull_dir/R-scape_noss", $synthetic, $decoy, $twotest);
add_hlx_file($profmark_synthetic_noss_hlx_fisher_out,    $family, "$syntheticnull_dir/R-scape_noss", $synthetic, $decoy, $fisher);
add_hlx_file($profmark_synthetic_noss_hlx_sidak_out,     $family, "$syntheticnull_dir/R-scape_noss", $synthetic, $decoy, $sidak);
add_hlx_file($profmark_synthetic_noss_hlx_lancaster_out, $family, "$syntheticnull_dir/R-scape_noss", $synthetic, $decoy, $lancaster);
add_hlx_file($profmark_synthetic_noss_hlx_wfisher_out,   $family, "$syntheticnull_dir/R-scape_noss", $synthetic, $decoy, $wfisher);

# added also as decoy to the "select" set
$twotest = 1;
add_cov_file($profmark_select_cov_out,                   $family, "$syntheticnull_dir/R-scape",      $synthetic, $decoy, $twotest);
add_hlx_file($profmark_select_hlx_fisher_out,            $family, "$syntheticnull_dir/R-scape",      $synthetic, $decoy, $fisher);
add_hlx_file($profmark_select_hlx_sidak_out,             $family, "$syntheticnull_dir/R-scape",      $synthetic, $decoy, $sidak);
add_hlx_file($profmark_select_hlx_lancaster_out,         $family, "$syntheticnull_dir/R-scape",      $synthetic, $decoy, $lancaster);
add_hlx_file($profmark_select_hlx_wfisher_out,           $family, "$syntheticnull_dir/R-scape",      $synthetic, $decoy, $wfisher);

$twotest = 0;
add_cov_file($profmark_select_noss_cov_out,              $family, "$syntheticnull_dir/R-scape_noss", $synthetic, $decoy, $twotest);
add_hlx_file($profmark_select_noss_hlx_fisher_out,       $family, "$syntheticnull_dir/R-scape_noss", $synthetic, $decoy, $fisher);
add_hlx_file($profmark_select_noss_hlx_sidak_out,        $family, "$syntheticnull_dir/R-scape_noss", $synthetic, $decoy, $sidak);
add_hlx_file($profmark_select_noss_hlx_lancaster_out,    $family, "$syntheticnull_dir/R-scape_noss", $synthetic, $decoy, $lancaster);
add_hlx_file($profmark_select_noss_hlx_wfisher_out,      $family, "$syntheticnull_dir/R-scape_noss", $synthetic, $decoy, $wfisher);

if ($do_rscape) {
    $cmd = "$rscape -s  --scmin -10000 --hpts 20000 --lancaster_join   --wfisher_join   --joinsubs   --outdir $syntheticnull_dir/R-scape_join      --nofigures   $syntheticnull_sto";
    print "$cmd\n";
    system("$cmd\n");
    $cmd = "$rscape     --scmin -10000 --hpts 20000 --lancaster_join   --wfisher_join   --joinsubs   --outdir $syntheticnull_dir/R-scape_noss_join --nofigures   $syntheticnull_sto";
    print "$cmd\n";
    system("$cmd\n");
}
add_hlx_file($profmark_synthetic_hlx_lancaster_join_out,      $family, "$syntheticnull_dir/R-scape_join",      $synthetic, $decoy, $lancaster_join);
add_hlx_file($profmark_synthetic_hlx_wfisher_join_out,        $family, "$syntheticnull_dir/R-scape_join",      $synthetic, $decoy, $wfisher_join);
add_hlx_file($profmark_synthetic_noss_hlx_lancaster_join_out, $family, "$syntheticnull_dir/R-scape_noss_join", $synthetic, $decoy, $lancaster_join);
add_hlx_file($profmark_synthetic_noss_hlx_wfisher_join_out,   $family, "$syntheticnull_dir/R-scape_noss_join", $synthetic, $decoy, $wfisher_join);

add_hlx_file($profmark_select_hlx_lancaster_join_out,         $family, "$syntheticnull_dir/R-scape_join",      $synthetic, $decoy, $lancaster_join);
add_hlx_file($profmark_select_hlx_wfisher_join_out,           $family, "$syntheticnull_dir/R-scape_join",      $synthetic, $decoy, $wfisher_join);
add_hlx_file($profmark_select_noss_hlx_lancaster_join_out,    $family, "$syntheticnull_dir/R-scape_noss_join", $synthetic, $decoy, $lancaster_join);
add_hlx_file($profmark_select_noss_hlx_wfisher_join_out,      $family, "$syntheticnull_dir/R-scape_noss_join", $synthetic, $decoy, $wfisher_join);

if ($do_rscape) {
    $cmd = "$rscape -s  --scmin -10000 --hpts 20000 --lancaster_double --wfisher_double --doublesubs --outdir $syntheticnull_dir/R-scape_double      --nofigures $syntheticnull_sto";
    print "$cmd\n";
    system("$cmd\n");
    $cmd = "$rscape     --scmin -10000 --hpts 20000 --lancaster_double --wfisher_double --doublesubs --outdir $syntheticnull_dir/R-scape_noss_double --nofigures $syntheticnull_sto";
    print "$cmd\n";
    system("$cmd\n");
}
add_hlx_file($profmark_synthetic_hlx_lancaster_double_out,      $family, "$syntheticnull_dir/R-scape_double",      $synthetic, $decoy, $lancaster_double);
add_hlx_file($profmark_synthetic_hlx_wfisher_double_out,        $family, "$syntheticnull_dir/R-scape_double",      $synthetic, $decoy, $wfisher_double);
add_hlx_file($profmark_synthetic_noss_hlx_lancaster_double_out, $family, "$syntheticnull_dir/R-scape_noss_double", $synthetic, $decoy, $lancaster_double);
add_hlx_file($profmark_synthetic_noss_hlx_wfisher_double_out,   $family, "$syntheticnull_dir/R-scape_noss_double", $synthetic, $decoy, $wfisher_double);

add_hlx_file($profmark_select_hlx_lancaster_double_out,         $family, "$syntheticnull_dir/R-scape_double",      $synthetic, $decoy, $lancaster_double);
add_hlx_file($profmark_select_hlx_wfisher_double_out,           $family, "$syntheticnull_dir/R-scape_double",      $synthetic, $decoy, $wfisher_double);
add_hlx_file($profmark_select_noss_hlx_lancaster_double_out,    $family, "$syntheticnull_dir/R-scape_noss_double", $synthetic, $decoy, $lancaster_double);
add_hlx_file($profmark_select_noss_hlx_wfisher_double_out,      $family, "$syntheticnull_dir/R-scape_noss_double", $synthetic, $decoy, $wfisher_double);

print "\ntrue_bp $select_true_bp  $synthetic_true_bp\n";
print   "true_hlx $select_true_hlx $synthetic_true_hlx\n";


#----------------------------------------------------------------------------------------------------------------------------------------------------

sub add_cov_file {
    my ($profmark_cov_out, $family, $rscapedir, $synthetic, $decoy, $twotest, $ret_true_bp) = @_;

    # append to outfile
    open(OUT, ">>$profmark_cov_out") || die;

    my $name = "$family";
    if ($synthetic) { $name .= "_synthetic"; }

    my $type = "$family";
    if ($decoy) { $type .= "_decoy"; }
    
    my @file;
    my $prefix = "$family";
    my $suffix = "sorted.cov";
 
    FUNCS::sorted_files($rscapedir, \@file, $suffix, $prefix);
    my $nf = $#file+1;
    print OUT "# nmsa $nf\n";

    my $eval;
    my $pval;
    my $score;
    my $i;
    my $j;
    for (my $n = 0; $n < $nf; $n++) {
	printf("cov $n/$nf> %s\n", $file[$n]);

	my $exp_bp;
	my $powerfile = "$file[$n]"; 
	if ($powerfile =~ /^(\S+).$suffix$/) { $powerfile = "$1.power"; }
	open(FILE, "$powerfile") || die;
	while(<FILE>) {
	    if (/^# BPAIRS expected to covary\s+(\S+)\s/) {
		$exp_bp = $1;
	    }
	}
	close(FILE);

	open(FILE, "$file[$n]") || die;
	while(<FILE>) {
	    if (/^#.+\[\d+\s\|\s\d+\s(\d+)\s\d+\s\|\s\S+\s\S+\s\S+\]/) {
		my $true_bp = $1;
		if ($ret_true_bp) { 
		    $$ret_true_bp = $true_bp;
		    print OUT "# true_bp $true_bp\n";
		    print OUT "# exp_bp  $exp_bp\n";
		}
	    }
	    elsif (/^#/) { }
	    elsif (/^\*\s+(\d+)\s+(\d+)\s+(\S+)\s+(\S+)\s+(\S+)\s+\d+/ && !$decoy) { # a tp only if in the proposed structure (others could be not annotated base pairs)
		$i     = $1;
		$j     = $2;
		$score = $3;
		$eval  = $4;
		$pval  = $5;
		print OUT "$eval $pval $score $i $j $name $type\n";
	    }
	    elsif (/^\*\s+(\d+)\s+(\d+)\s+(\S+)\s+(\S+)\s+(\S+)\s+\d+/ && $decoy) { # in twotest = 1, we only look at decoys tested as basepairs
		$i     = $1;
		$j     = $2;
		$score = $3;
		$eval  = $4;
		$pval  = $5;
		print OUT "$eval $pval $score $i $j $name $type\n";
	    }
	    elsif (/(\d+)\s+(\d+)\s+(\S+)\s+(\S+)\s+(\S+)\s+\d+/ && $decoy && !$twotest) { # in noss (twotest = 0) all decoys count as fp
		$i     = $1;
		$j     = $2;
		$score = $3;
		$eval  = $4;
		$pval  = $5;
		print OUT "$eval $pval $score $i $j $name $type\n";
	    }
	}
	close(FILE);
    }
    close(OUT);
}

sub add_hlx_file {
    my ($profmark_hlx_out, $family, $rscapedir, $synthetic, $decoy, $aggtype, $ret_true_hlx) = @_;

    # append to outfile
    open(OUT, ">>$profmark_hlx_out") || die;

    my $name = "$family";
    if ($synthetic) { $name .= "_synthetic"; }

    my $type = "$family";
    if ($decoy) { $type .= "_decoy"; }
    
    my @file;
    my $prefix = "$family";
    my $suffix = "helixcov";

    FUNCS::sorted_files($rscapedir, \@file, $suffix, $prefix);
    my $nf = $#file+1;
    print OUT "# nmsa $nf\n";

    for (my $n = 0; $n < $nf; $n++) {
	printf("hlx $n/$nf> %s\n", $file[$n]);

	my $eval;
	my $pval;
	my $left;
	my $right;
	my $nbp_per_hl;
	my $ncov_per_hl;
	my $pval1;
	my $pval2;
	my $minpval;

	my $nss = 0;
	my $nhl_per_ss;
	my $nbp_per_ss;
	my $fracbp_per_ss;
	my $len_hlx;

	my $len_hl;
	my $len_ss;

	# because the helixcov file only has p-values for base pairs, we need to reach to the .cov file for the relationship between E-val to p-val
	my $covfile = "$file[$n]";
	if ($covfile =~ /^(\S+).helixcov$/) { $covfile = "$1.sorted.cov"; }
	my $N = 0; # Eval = N * pval
	
	open(COV, "$covfile") || die;
	while(<COV>) {
	    if (/\*\s+\d+\s+\d+\s+\S+\s+(\S+)\s+(\S+)\s+\d+/) {
		my $eval = $1;
		my $pval = $2;
		$N = $eval/$pval;
		if ($N > 0) { last; }
	    }   
	}
	close(COV);
	
	open(FILE, "$file[$n]") || die;
	while(<FILE>) {
	    if (/^\# RMs = (\d+)\s+L = (\d+)/) {
		my $nhl_per_ss_new = $1;
		my $len_ss_new     = $2;

		if ($nss > 0) {
		    $fracbp_per_ss = 2*$nbp_per_ss/$len_ss;
		    print OUT "# SS $nhl_per_ss $nbp_per_ss $len_ss\n";
		}
		$nss ++;
		
		# init
		$nbp_per_ss = 0;
		$len_ss     = $len_ss_new;
		$nhl_per_ss = $nhl_per_ss_new;

		if ($ret_true_hlx) { 
		    $$ret_true_hlx = $nhl_per_ss;
		    print OUT "# true_hlx $nhl_per_ss\n";
		}
	    }
	    elsif (/^\# RM\s+(\S+)\s+(\S+),\s+nbp = (\d+) nbp_cov = (\d+)\s*$/) {
		$left         = $1;
		$right        = $2;
		$nbp_per_hl   = $3;
		#$ncov_per_hl = $4;
		$ncov_per_hl  = 0;  # calculate directly
		$len_hl       = 2*$nbp_per_hl;
		$nbp_per_ss  += $nbp_per_hl;
	    }
	    elsif (/^\# pvals:\s(\S+)\s*$/) {
		my $pvals = $1;
		# split
		my @pvals = split(/,/, $pvals);

		$minpval = 9999999999999999.;
		for (my $n = 0; $n < $nbp_per_hl; $n ++) {
		    
		    if ($pvals[$n] < $minpval) { $minpval = $pvals[$n]; }
		    
		    if ($N > 0 && $pvals[$n]*$N < 0.05) { 
			$ncov_per_hl ++; 
		    }
		}

		$pval1 = -1;
		$pval2 = -1;
		if    ($nbp_per_hl == 2) {
		    $pval1 = $pvals[0];
		    $pval2 = $pvals[1];
		}
		elsif ($nbp_per_hl > 2) {
		    my $r1 = int(rand($nbp_per_hl));
		    my $r2 = int(rand($nbp_per_hl));

		    while ($r2 == $r1) {
			$r2 = int(rand($nbp_per_hl));
		    }
		    $pval1 = $pvals[$r1];
		    $pval2 = $pvals[$r2];
		}
	    }
	    elsif (/# aggregated\s+$aggtype\s+E-value:\s+(\S+)\s+P-value:\s+(\S+)\s+/) {
		$eval = $1;
		$pval = $2;
		print OUT "$eval $pval $left $right $nbp_per_hl $ncov_per_hl $name $type $pval1 $pval2 $minpval\n";
	    }	    
	}
	close(FILE);

	# last case
	print OUT "# SS $nhl_per_ss $nbp_per_ss $len_ss\n";

    }    
    close(OUT);    
}
```
