## Supplementary material for "RNA covariation at helix-level resolution for the identification of evolutionarily conserved RNA structure": profmark_analyze.pl

```
#!/usr/bin/perl -w
# 
# profmark_analyze
#
use strict;
use Class::Struct;
use POSIX;
    
use constant GNUPLOT => '/usr/local/bin/gnuplot';
#use constant GNUPLOT => '/n/helmod/apps/centos7/Core/gnuplot/5.2.6-fasrc01/bin/gnuplot';

use lib '/Users/erivas/src/Mysrc/R-scape/scripts';
#use lib '/n/eddy_lab/users/erivas/rscape_v2.0.0.p/scripts';
use FUNCS;

use vars qw ($opt_v $opt_G);  # required if strict used
use Getopt::Std;
getopts ('Gv');

# Print a helpful message if the user provides no input file.
if (!@ARGV) {
    print "usage:  profmark_analyze.pl [options] <resultsdir>  \n\n";
    print "options:\n";
    print "-v    :  be verbose\n";
    exit;
}

my $profmark_dir = shift;
my $results_dir  = "$profmark_dir/results";
system("rm $results_dir/profmark*\n");

my $viewplot = 0;
my $plotonly = 0;
if ($opt_G) { $plotonly = 1; }

my $eval_cutoff = 1000;

my $nali_select         = 0;
my $nali_select_noss    = 0;
my $nali_synthetic      = 0;
my $nali_synthetic_noss = 0;
all_parse_covfile($results_dir, \$nali_select, \$nali_select_noss, \$nali_synthetic, \$nali_synthetic_noss);
printf("nali_select= $nali_select/$nali_select_noss nali_synthetic = $nali_synthetic/$nali_synthetic_noss\n");

my $nagg = 8;
my @aggmethod;
$aggmethod[0] = "fisher";
$aggmethod[1] = "sidak";
$aggmethod[2] = "lancaster";
$aggmethod[3] = "wfisher";
$aggmethod[4] = "lancaster_join";
$aggmethod[5] = "wfisher_join";
$aggmethod[6] = "lancaster_double";
$aggmethod[7] = "wfisher_double";
all_parse_hlxfile($nagg, \@aggmethod, $results_dir);

all_pvalue_pairs($results_dir);

sub all_parse_covfile {
    my ($results_dir, $ret_nali_select, $ret_nali_select_noss, $ret_nali_synthetic, $ret_nali_synthetic_noss) = @_;

    my $string = "select.cov";
    my $rocfile1 = "$results_dir/$string.roc";
    my $unifile1 = "$results_dir/$string.uni";
    if ($plotonly == 0) {
	system("rm $results_dir/$string.*\n");
	$$ret_nali_select = parse_covfile($rocfile1, $unifile1, $results_dir, $string);
    }
    
    $string = "select.noss.cov";
    my $rocfile2 = "$results_dir/$string.roc";
    my $unifile2 = "$results_dir/$string.uni";
    if ($plotonly == 0) {
	system("rm $results_dir/$string.*\n");
	$$ret_nali_select_noss = parse_covfile($rocfile2, $unifile2, $results_dir, $string);
    }
    
    $string = "synthetic.cov";
    my $rocfile3 = "$results_dir/$string.roc";
    my $unifile3 = "$results_dir/$string.uni";
    if ($plotonly == 0) {
	system("rm $results_dir/$string.*\n");
	$$ret_nali_synthetic = parse_covfile($rocfile3, $unifile3, $results_dir, $string);
    }
    
    $string = "synthetic.noss.cov";
    my $rocfile4 = "$results_dir/$string.roc";
    my $unifile4 = "$results_dir/$string.uni";
    if ($plotonly == 0) {
	system("rm $results_dir/$string.*\n");
	$$ret_nali_synthetic_noss = parse_covfile($rocfile4, $unifile4, $results_dir, $string);
    }
    
    gnuplot_rocplot_cov($results_dir, $rocfile1, $rocfile2, $rocfile3, $rocfile4);
    gnuplot_uniform_cov($results_dir, $unifile1, $unifile2, $unifile3, $unifile4);
}


sub all_parse_hlxfile {
    my ($nagg, $aggmethod_ref, $results_dir) = @_;

    my @rocfile1;
    my @rocfile2;
    my @rocfile3;
    my @rocfile4;
    
    my @unifile1;
    my @unifile2;
    my @unifile3;
    my @unifile4;
    
    my @nocfile1;
    my @nocfile2;
    my @nocfile3;
    my @nocfile4;

    for (my $agg = 0; $agg < $nagg; $agg ++) {
	one_parse_hlxfile($aggmethod_ref->[$agg], $results_dir, 
			  \$rocfile1[$agg], \$rocfile2[$agg], \$rocfile3[$agg], \$rocfile4[$agg], 
			  \$unifile1[$agg], \$unifile2[$agg], \$unifile3[$agg], \$unifile4[$agg], 
			  \$nocfile1[$agg], \$nocfile2[$agg], \$nocfile3[$agg], \$nocfile4[$agg]);
    }

    gnuplot_rocplot_hlx($results_dir, $nagg, $aggmethod_ref, \@rocfile1, \@rocfile2, \@rocfile3, \@rocfile4, $nocfile1[4], $nocfile2[4], $nocfile3[4], $nocfile4[4]);
    gnuplot_uniform_hlx($results_dir, $nagg, $aggmethod_ref, \@unifile1, \@unifile2, \@unifile3, \@unifile4);
}

sub all_pvalue_pairs {
    my ($results_dir) = @_;

    my $string;
    my $cmd;

    $string           = "select";
    my $pairsfile1r   = "$results_dir/$string.ppairs";
    my $pairsfile1n   = "$results_dir/$string.ppairs.null";
    my @hlxstatfile1r;
    $hlxstatfile1r[0] = "$results_dir/$string.hlx.len.stats";
    $hlxstatfile1r[1] = "$results_dir/$string.hlx.nhl.stats";
    $hlxstatfile1r[2] = "$results_dir/$string.hlx.fbp.stats";
    my @hlxstatfile1n;
    $hlxstatfile1n[0] = "$results_dir/$string.hlx.len.stats.null";
    $hlxstatfile1n[1] = "$results_dir/$string.hlx.nhl.stats.null";
    $hlxstatfile1n[2] = "$results_dir/$string.hlx.fbp.stats.null";
    if ($plotonly == 0) {
	my $outfile1      = "$results_dir/$string.hlx.lancaster.out";
	$cmd = "$results_dir/*.$string.hlx.lancaster.out";
	system("cat $cmd > $outfile1\n");
 
	system("rm $results_dir/$string.ppairs\n");
	pvalue_pairs($pairsfile1r, $pairsfile1n, \@hlxstatfile1r, \@hlxstatfile1n, $outfile1, $string);
	
	#system("rm $outfile1\n");
     }
    
    $string           = "synthetic";
    my $pairsfile2r   = "$results_dir/$string.ppairs";
    my $pairsfile2n   = "$results_dir/$string.ppairs.null";
    my @hlxstatfile2r;
    $hlxstatfile2r[0] = "$results_dir/$string.hlx.len.stats";
    $hlxstatfile2r[1] = "$results_dir/$string.hlx.nhl.stats";
    $hlxstatfile2r[2] = "$results_dir/$string.hlx.fbp.stats";
    my @hlxstatfile2n;
    $hlxstatfile2n[0] = "$results_dir/$string.hlx.len.stats.null";
    $hlxstatfile2n[1] = "$results_dir/$string.hlx.nhl.stats.null";
    $hlxstatfile2n[2] = "$results_dir/$string.hlx.fbp.stats.null";
    if ($plotonly == 0) {
	my $outfile2      = "$results_dir/$string.hlx.lancaster.out";
	$cmd = "$results_dir/*.$string.hlx.lancaster.out";
	system("cat $cmd > $outfile2\n");
	
	system("rm $results_dir/$string.ppairs\n");
	pvalue_pairs($pairsfile2r, $pairsfile2n, \@hlxstatfile2r, \@hlxstatfile2n, $outfile2, $string);
	
	#system("rm $outfile2\n");    
   }
 
    gnuplot_ppairs ($results_dir, $pairsfile1r,    $pairsfile1n,    $pairsfile2r,    $pairsfile2n);
    gnuplot_hlxstat($results_dir, \@hlxstatfile1r, \@hlxstatfile1n, \@hlxstatfile2r, \@hlxstatfile2n);
}

sub pvalue_pairs {
    my ($pairsfiler, $pairsfilen, $hlxstatfiler_ref, $hlxstatfilen_ref, $outfile, $string) = @_;

   open(PAIRR, ">$pairsfiler")   || die;
   open(PAIRN, ">$pairsfilen")   || die;

    my $N = 500;
    my $k = 1;
    my $shift = 0;
    my @his_len_hlr;
    my @his_len_hln;
    my @his_nhl_per_ssr;
    my @his_nhl_per_ssn;
    my @his_fracbp_per_ssr;
    my @his_fracbp_per_ssn;
    FUNCS::init_histo_array($N, $k, \@his_len_hlr);
    FUNCS::init_histo_array($N, $k, \@his_len_hln);
    FUNCS::init_histo_array($N, $k, \@his_nhl_per_ssr);
    FUNCS::init_histo_array($N, $k, \@his_nhl_per_ssn);

    my $N3 = 1;
    my $k3 = 100;
    FUNCS::init_histo_array($N3, $k3, \@his_fracbp_per_ssr);
    FUNCS::init_histo_array($N3, $k3, \@his_fracbp_per_ssn);
    
    my $isreal;
    open(FILE, "$outfile") || die;
    while(<FILE>) {
	if (/^\# nmsa/) {
	    $isreal = 0;
	}
	elsif (/^\# true_hlx/) {
	    $isreal = 1;
	}
	elsif (/^\# SS\s+(\d+)\s+(\d+)\s+(\d+)\s*$/) {
	    my $nhl_per_ss = $1;
	    my $nbp_per_ss = $2;
	    my $len_per_ss = $3;
	    my $fracbp_per_ss = 2*$nbp_per_ss/$len_per_ss;
	    
	    # histogram of nhl_per_ss
	    if ($isreal) { FUNCS::fill_histo_array(1, $nhl_per_ss, $N, $k, $shift, \@his_nhl_per_ssr); }
	    else         { FUNCS::fill_histo_array(1, $nhl_per_ss, $N, $k, $shift, \@his_nhl_per_ssn); }

	    # histogram of fracbp_per_ss
	    if ($isreal) { FUNCS::fill_histo_array(1, $fracbp_per_ss, $N3, $k3, $shift, \@his_fracbp_per_ssr); }
	    else         { FUNCS::fill_histo_array(1, $fracbp_per_ss, $N3, $k3, $shift, \@his_fracbp_per_ssn); }

	}
	#0.260057 0.260057 5-59 64-118 55 55 RF00001 RF00001_decoy 0.665069 0.723326
	elsif (/^\S+\s+\S+\s+\S+\s+\S+\s+(\d+)\s+\d+\s+\S+\s+(\S+)\s+(\S+)\s+(\S+)\s*$/) {
	    my $nbp   = $1;
	    my $name  = $2;
	    my $pval1 = $3;
	    my $pval2 = $4;

	    my $len_hl = 2*$nbp;

	    if ($isreal && $name =~ /decoy/) { print "bad assignment for $outfile\n"; exit; }
	    
	    # save the two random pvals
	    if ($isreal) { printf PAIRR "%g %g\n", $pval1, $pval2; }
	    else         { printf PAIRN "%g %g\n", $pval1, $pval2; }

	    # histogram of len_hl
	    if ($isreal) { FUNCS::fill_histo_array(1, $len_hl, $N, $k, $shift, \@his_len_hlr); }
	    else         { FUNCS::fill_histo_array(1, $len_hl, $N, $k, $shift, \@his_len_hln); }

	}
    }
    close(FILE);
    
    close(PAIRR);
    close(PAIRN);

    FUNCS::write_histogram($N,  $k,  $shift, \@his_len_hlr,        1, $hlxstatfiler_ref->[0], 0);
    FUNCS::write_histogram($N,  $k,  $shift, \@his_nhl_per_ssr,    1, $hlxstatfiler_ref->[1], 0);
    FUNCS::write_histogram($N3, $k3, $shift, \@his_fracbp_per_ssr, 1, $hlxstatfiler_ref->[2], 0);
    
    FUNCS::write_histogram($N,  $k,  $shift, \@his_len_hln,        1, $hlxstatfilen_ref->[0], 0);
    FUNCS::write_histogram($N,  $k,  $shift, \@his_nhl_per_ssn,    1, $hlxstatfilen_ref->[1], 0);
    FUNCS::write_histogram($N3, $k3, $shift, \@his_fracbp_per_ssn, 1, $hlxstatfilen_ref->[2], 0);

}

sub eval_from_covroc {
    my ($eval_target, $rocfile) = @_;
    
    my $evalfile = "$rocfile.eval";
    
    open(OUT, ">$evalfile") || die;
    
    open(FILE, "$rocfile") || die;
    while (<FILE>)
    {
	if (/^(\S+)\s+\S+\s+\S+\s+\S+\s+\d+\s+\d+\s+(\S+)\s+\S+\s*$/) {
	    my $oeval = $1;
	    my $eval  = $2;
	    if ($eval > $eval_target) {
		print OUT "$oeval 0\n";
		print OUT "$oeval 1\n";
		last;
	    }
	}
    }
    close(FILE);
    close(OUT);
    
    return $evalfile;
}

sub eval2eval_covroc {
    my ($rocfile) = @_;
    
    my $evalfile = "$rocfile.eval2eval";
    
    open(OUT, ">$evalfile") || die;
    
    open(FILE, "$rocfile") || die;
    while (<FILE>)
    {
	if (/^(\S+)\s+\S+\s+\S+\s+\S+\s+\d+\s+\d+\s+(\S+)\s+\S+\s*$/) {
	    my $oeval = $1;
	    my $eval  = $2;
	    if ($oeval > 0) { print OUT "$oeval $eval\n"; }
	}
    }
    close(FILE);
    close(OUT);
    
    return $evalfile;
}

sub eval_from_hlxroc {
    my ($eval_target, $rocfile) = @_;
    
    my $evalfile = "$rocfile.eval";
    
    open(OUT, ">$evalfile") || die;
    
    open(FILE, "$rocfile") || die;
    while (<FILE>)
    {
	if (/^(\S+)\s+\S+\s+\S+\s+\d+\s+\d+\s+(\S+)\s+\S+\s*$/) {
	    my $oeval = $1;
	    my $eval  = $2;
	    if ($eval > $eval_target) {
		print OUT "$oeval 0\n";
		print OUT "$oeval 1\n";
		last;
	    }
	}
    }
    close(FILE);
    close(OUT);
    
    return $evalfile;
}

sub eval2eval_hlxroc {
    my ($rocfile) = @_;
    
    my $evalfile = "$rocfile.eval2eval";
    
    open(OUT, ">$evalfile") || die;
    
    open(FILE, "$rocfile") || die;
    while (<FILE>)
    {
	if (/^(\S+)\s+\S+\s+\S+\s+\d+\s+\d+\s+(\S+)\s+\S+\s*$/) {
	    my $oeval = $1;
	    my $eval  = $2;
	    if ($oeval > 0) { print OUT "$oeval $eval\n"; }
	}
    }
    close(FILE);
    close(OUT);
    
    return $evalfile;
}

sub one_parse_hlxfile {
    my ($aggmethod, $results_dir, 
	$ret_rocfile1, $ret_rocfile2, $ret_rocfile3, $ret_rocfile4, 
	$ret_unifile1, $ret_unifile2, $ret_unifile3, $ret_unifile4, 
	$ret_nocfile1, $ret_nocfile2, $ret_nocfile3, $ret_nocfile4) = @_;

    my $string;

    $string      = "select.hlx".".$aggmethod";
    $$ret_rocfile1 = "$results_dir/$string.roc";
    $$ret_unifile1 = "$results_dir/$string.uni";
    $$ret_nocfile1 = "$results_dir/$string.noc";
    if ($plotonly == 0) {
	system("rm $results_dir/$string.*\n");
	parse_hlxfile($$ret_rocfile1, $$ret_unifile1, $$ret_nocfile1, $results_dir, $string);
    }

    $string      = "select.noss.hlx".".$aggmethod";
    $$ret_rocfile2 = "$results_dir/$string.roc";
    $$ret_unifile2 = "$results_dir/$string.uni";
    $$ret_nocfile2 = "$results_dir/$string.noc";
    if ($plotonly == 0) {
	system("rm $results_dir/$string.*\n");
	parse_hlxfile($$ret_rocfile2, $$ret_unifile2, $$ret_nocfile2, $results_dir, $string);
    }
    
    $string      = "synthetic.hlx".".$aggmethod";
    $$ret_rocfile3 = "$results_dir/$string.roc";
    $$ret_unifile3 = "$results_dir/$string.uni";
    $$ret_nocfile3 = "$results_dir/$string.noc";
    if ($plotonly == 0) {
	system("rm $results_dir/$string.*\n");
	parse_hlxfile($$ret_rocfile3, $$ret_unifile3, $$ret_nocfile3, $results_dir, $string);
    }
    
    $string      = "synthetic.noss.hlx".".$aggmethod";
    $$ret_rocfile4 = "$results_dir/$string.roc";
    $$ret_unifile4 = "$results_dir/$string.uni";
    $$ret_nocfile4 = "$results_dir/$string.noc";
    if ($plotonly == 0) {
	system("rm $results_dir/$string.*\n");
	parse_hlxfile($$ret_rocfile4, $$ret_unifile4, $$ret_nocfile4, $results_dir, $string);
    }
}

sub gnuplot_rocplot_cov {
    my ($results_dir, $rocfile1, $rocfile2, $rocfile3, $rocfile4) = @_;
    
    my $psfile = "$results_dir/profmark.cov.roc.ps";    
    my $pdfile = "$results_dir/profmark.cov.roc.pdf";
    open(GP,'|'.GNUPLOT) || die "Gnuplot: $!";
    print GP "set terminal postscript color 14\n";
    FUNCS::gnuplot_define_styles (*GP);
    print GP "set output '$psfile'\n";

    my $xlabel = "mean false positives per alignment (observed E-value)";
    my $which  = 1; # Observed E-value
    my $xmin  = 0.0005;
    my $xmax  = 1;
    gnuplot_rocplot_cov_one(*GP, $which, $xlabel, $xmin, $xmax, $rocfile1, $rocfile2, $rocfile3, $rocfile4, 1, 1);
    
    $xlabel = "false discovery rate";
    $which  = 2; # False Discovery Rate;
    $xmin   = 0.0005;
    $xmax   = 1;
    gnuplot_rocplot_cov_one(*GP, $which, $xlabel, $xmin, $xmax, $rocfile1, $rocfile2, $rocfile3, $rocfile4, 1, 0);

    $xlabel = "false positives";
    $which  = 5; # False positives
    $xmin = 0;
    $xmax = 50;
    gnuplot_rocplot_cov_one(*GP, $which, $xlabel, $xmin, $xmax, $rocfile1, $rocfile2, $rocfile3, $rocfile4, 0, 0);

    close(GP);
    
    system("ps2pdf $psfile $pdfile\n");
    system("rm $psfile\n");
    
    if ($viewplot) {
	system("open $pdfile&\n"); 
    }
    
}

sub gnuplot_rocplot_cov_one {
    my ($gp, $which, $xlabel, $xmin, $xmax, $rocfile1, $rocfile2, $rocfile3, $rocfile4, $xlogscale, $doeval) = @_;

    my $evalfile1 = eval_from_covroc(0.05, $rocfile1);
    my $evalfile2 = eval_from_covroc(0.05, $rocfile2);
    my $evalfile3 = eval_from_covroc(0.05, $rocfile3);
    my $evalfile4 = eval_from_covroc(0.05, $rocfile4);
    
    my $eval2evalfile1 = eval2eval_covroc($rocfile1);
    my $eval2evalfile2 = eval2eval_covroc($rocfile2);
    my $eval2evalfile3 = eval2eval_covroc($rocfile3);
    my $eval2evalfile4 = eval2eval_covroc($rocfile4);
    
    print $gp "set xlabel '$xlabel'\n";

    my $ylabel = "Fraction of base pairs that covary"; 
    print $gp "set ylabel '$ylabel'\n";

    print $gp "set key right bottom\n";
    if ($xlogscale) { print $gp "set logscale x\n"; } else { print $gp "unset logscale x\n"; }
    print $gp "set size nosquare\n";
    print $gp "unset logscale y\n";

    print $gp "set yrange [0:1.01]\n";
    if ($xmin >= 0 && $xmax >= 0) {
	print $gp "set xrange [$xmin:$xmax]\n";
    }
    elsif ($xmax >= 0) {
	print $gp "set xrange [*:$xmax]\n";
    }
    elsif ($xmax >= 0) {
	print $gp "set xrange [$xmin:*]\n";
    }
    else {
	print $gp "set xrange [*:*]\n";
    }
    
    my $cmd = "plot ";
    my $m = 1;
    $cmd .= "'$rocfile1'  using $which:3 title 'Rfam'           with lines ls $m, "; 
    if ($doeval) { $cmd .= "'$evalfile1' using 1:2     title ''                with lines ls $m, "; } 
    $m ++;
    
    $cmd .= "'$rocfile2'  using $which:3 title 'Rfam noss'      with lines ls $m, ";     
    if ($doeval) { $cmd .= "'$evalfile2' using 1:2     title ''                with lines ls $m, "; }      
    $m ++;
    
    $cmd .= "'$rocfile3'  using $which:3 title 'Synthetic'      with lines ls $m, ";       
    if ($doeval) { $cmd .= "'$evalfile3' using 1:2     title ''                with lines ls $m, "; }    
    $m ++;

    if ($doeval) {
	$cmd .= "'$rocfile4'  using $which:3 title 'Synthetic noss' with lines ls $m, ";
	$cmd .= "'$evalfile4' using 1:2     title ''                with lines ls $m";
    }
    else {
	$cmd .= "'$rocfile4'  using $which:3 title 'Synthetic noss' with lines ls $m";
    }
    print $gp "$cmd\n";
    #print "$cmd";

    if ($doeval) {
	$xlabel = "observed E-value";
	$ylabel = "expected E-value";
	print $gp "set logscale x\n";
	print $gp "set logscale y\n";
	print $gp "set size square\n";
	print $gp "set yrange [0.00001:10]\n";
	print $gp "set xrange [0.00001:10]\n";
	print $gp "set xlabel '$xlabel'\n";
	print $gp "set ylabel '$ylabel'\n";
	
	$cmd = "plot ";
	$m = 1;
	$cmd .= "x, ";
	$cmd .= "'$eval2evalfile1' using 1:2 title 'Rfam'  with lines ls $m"; 
	print $gp "$cmd\n";
	#print "$cmd";
	
	$cmd = "plot ";
	$m = 1;
	$cmd .= "x, ";
	$cmd .= "'$eval2evalfile2' using 1:2 title 'Rfam-noss'  with lines ls $m"; 
	print $gp "$cmd\n";
	#print "$cmd";
	
	$cmd = "plot ";
	$m = 1;
	$cmd .= "x, ";
	$cmd .= "'$eval2evalfile3' using 1:2 title 'Rfam-Synthetic'  with lines ls $m"; 
	print $gp "$cmd\n";
	#print "$cmd";
	
	$cmd = "plot ";
	$m = 1;
	$cmd .= "x, ";
	$cmd .= "'$eval2evalfile4' using 1:2 title 'Rfam-Synthetic-noss'  with lines ls $m"; 
	print $gp "$cmd\n";
	#print "$cmd";
    }
}


sub gnuplot_rocplot_hlx {
    my ($results_dir, $nagg, $aggmethod_ref, $rocfile1_ref, $rocfile2_ref, $rocfile3_ref, $rocfile4_ref, $nocfile1, $nocfile2, $nocfile3, $nocfile4) = @_;

    my $psfile = "$results_dir/profmark.hlx.roc.ps";    
    my $pdfile = "$results_dir/profmark.hlx.roc.pdf";

    open(GP,'|'.GNUPLOT) || die "Gnuplot: $!";
    print GP "set terminal postscript color 14\n";
    print GP "set output '$psfile'\n";
    FUNCS::gnuplot_define_styles (*GP);

    # Observed E-value
    my $ylabel = "Fraction of helices that covary";
    my $xlabel = "False positives per alignment (observd E-value)";
    my $which  = 1; # False Positive Rate
    my $xmin   = -1;
    my $xmax   = 1;
    gnuplot_rocplot_hlx_one(*GP, $which, $xlabel, $xmin, $xmax, $ylabel, $nagg, $aggmethod_ref, 
			    $rocfile1_ref, $rocfile2_ref, $rocfile3_ref, $rocfile4_ref, $nocfile1, $nocfile2, $nocfile3, $nocfile4, 1, 1);
    
    #FDR
    $xlabel = "False Discovery Rate";
    $which  = 2; # False Discovery Rate
    $xmin = -1;
    $xmax = 1;
    gnuplot_rocplot_hlx_one(*GP, $which, $xlabel, $xmin, $xmax, $ylabel, $nagg, $aggmethod_ref, 
			    $rocfile1_ref, $rocfile2_ref, $rocfile3_ref, $rocfile4_ref, $nocfile1, $nocfile2, $nocfile3, $nocfile4, 1, 0);
 
    #FP
    $xlabel = "False Positives";
    $which  = 4;
    $xmin   = 0;
    $xmax   = 50;
    gnuplot_rocplot_hlx_one(*GP, $which, $xlabel, $xmin, $xmax, $ylabel, $nagg, $aggmethod_ref, 
			    $rocfile1_ref, $rocfile2_ref, $rocfile3_ref, $rocfile4_ref, $nocfile1, $nocfile2, $nocfile3, $nocfile4, 0, 0);
 
    close(GP);
    
    system("ps2pdf $psfile $pdfile\n");
    system("rm $psfile\n");
    
    if ($viewplot) {
	system("open $pdfile&\n"); 
    }
}

sub gnuplot_rocplot_hlx_one {
    my ($gp, $which, $xlabel, $xmin, $xmax, $ylabel, $nagg, $aggmethod_ref, 
	$rocfile1_ref, $rocfile2_ref, $rocfile3_ref, $rocfile4_ref, $nocfile1, $nocfile2, $nocfile3, $nocfile4,
	$xlogscale, $doeval) = @_;
    
    print $gp "set key right bottom\n";
    if ($xlogscale) { print $gp "set logscale x\n"; } else { print $gp "unset logscale x\n"; }
    
    my $title = "Rfam";
    gnuplot_rocplot_hlx_onecase($gp, $title, $which, $xlabel, $xmin, $xmax, $ylabel, $nagg, $aggmethod_ref, $rocfile1_ref, $nocfile1, $doeval);
      
    $title = "Rfam noss";
    gnuplot_rocplot_hlx_onecase($gp, $title, $which, $xlabel, $xmin, $xmax, $ylabel, $nagg, $aggmethod_ref, $rocfile2_ref, $nocfile2, $doeval);

    $title = "synthetic";
    gnuplot_rocplot_hlx_onecase($gp, $title, $which, $xlabel, $xmin, $xmax, $ylabel, $nagg, $aggmethod_ref, $rocfile3_ref, $nocfile3, $doeval);
  
    $title = "Synthetic noss";
    gnuplot_rocplot_hlx_onecase($gp, $title, $which, $xlabel, $xmin, $xmax, $ylabel, $nagg, $aggmethod_ref, $rocfile4_ref, $nocfile4, $doeval);    
}

    
sub gnuplot_rocplot_hlx_onecase {
    my ($gp, $title, $which, $xlabel, $xmin, $xmax, $ylabel, $nagg, $aggmethod_ref, $rocfile_ref, $nocfile, $doeval) = @_;
    
    my $evalfile;
    my $eval2evalfile;

    print $gp "set size nosquare\n";
    print $gp "unset logscale y\n";
    print $gp "set title '$title'\n";
    print $gp "set yrange [0:1.01]\n";
    if    ($xmin >= 0 && $xmax >= 0) { print $gp "set xrange [$xmin:$xmax]\n"; } 
    elsif ($xmin >= 0)               { print $gp "set xrange [$xmin:*]\n"; } 
    elsif ($xmax >= 0)               { print $gp "set xrange [*:$xmax]\n"; } 
    else                             { print $gp "set xrange [*:*]\n"; }
    print $gp "set xlabel '$xlabel'\n";
    print $gp "set ylabel '$ylabel'\n";

    my $cmd = "plot ";
    my $m = 1;
    $cmd .= "'$nocfile' using 1:2 title 'no agg'  ls 9, "; 
    for (my $agg = 0; $agg < $nagg-1; $agg ++) {
	$evalfile = eval_from_hlxroc(0.05, $rocfile_ref->[$agg]);

	$cmd .= "'$rocfile_ref->[$agg]' using $which:3 title '$aggmethod_ref->[$agg]' with lines ls $m, "; 
	if ($doeval) { 
	    $cmd .= "'$evalfile' using 1:2 title '' with lines ls $m, "; 
	}
	$m ++;
    }
    $evalfile = eval_from_hlxroc(0.05, $rocfile_ref->[$nagg-1]);
    $cmd .= "'$rocfile_ref->[$nagg-1]' using $which:3 title '$aggmethod_ref->[$nagg-1]' with lines ls $m, "; 
    if ($doeval) { 
	$cmd .= "'$evalfile' using 1:2 title '' with lines ls $m"; } else { $cmd .= "\n"; 
    }
    print $gp "$cmd\n";

    if ($doeval) {
	my $xlabel = "observed Aggregated E-value";
	my $ylabel = "expected Aggregated E-value";
	print $gp "set title '$title'\n";
	print $gp "set logscale x\n";
	print $gp "set logscale y\n";
	print $gp "set size square\n";
	print $gp "set yrange [0.00001:10]\n";
	print $gp "set xrange [0.00001:10]\n";
	print $gp "set xlabel '$xlabel'\n";
	print $gp "set ylabel '$ylabel'\n";
	
	for (my $agg = 0; $agg < $nagg; $agg ++) {
	    $eval2evalfile = eval2eval_hlxroc($rocfile_ref->[$agg]);

	    $cmd = "plot ";
	    $m = 1;
	    $cmd .= "x, ";
	    $cmd .= "'$eval2evalfile' using 1:2 title '$aggmethod_ref->[$agg]'  with lines ls $m"; 
	    print $gp "$cmd\n";
	    #print "$cmd";
	}
    }
}

sub gnuplot_uniform_cov {
    my ($results_dir, $unifile1, $unifile2, $unifile3, $unifile4) = @_;
    
    my $psfile = "$results_dir/profmark.cov.uniform.ps";    
    my $pdfile = "$results_dir/profmark.cov.uniform.pdf";
    open(GP,'|'.GNUPLOT) || die "Gnuplot: $!";
    print GP "set terminal postscript color 14\n";
    FUNCS::gnuplot_define_styles (*GP);
    print GP "set output '$psfile'\n";

    my $xlabel = "Null base pair P-values";
    gnuplot_uniform_cov_one(*GP, $xlabel, $unifile1, $unifile2, $unifile3, $unifile4);
    close(GP);
    
    system("ps2pdf $psfile $pdfile\n");
    system("rm $psfile\n");
    
    if ($viewplot) {
	system("open $pdfile&\n"); 
    }
}

sub gnuplot_uniform_cov_one {
    my ($gp, $xlabel, $unifile1, $unifile2, $unifile3, $unifile4) = @_;

    print $gp "set xlabel '$xlabel'\n";

    my $ylabel = "Occurrence"; 
    print $gp "set ylabel '$ylabel'\n";

    print $gp "set key right top\n";
    print $gp "set xrange [0:1.01]\n";

    my $cmd = "plot ";
    my $m = 1;
    $cmd .= "'$unifile1'  using 1:2 title 'Rfam'           with boxes ls $m"; 
    print $gp "$cmd\n";
    $m ++;

    $cmd  = "plot ";
    $cmd .= "'$unifile2'  using 1:2 title 'Rfam noss'      with boxes ls $m";     
    print $gp "$cmd\n";
    $m ++;
    
    $cmd  = "plot ";
    $cmd .= "'$unifile3'  using 1:2 title 'Synthetic'      with boxes ls $m";       
    print $gp "$cmd\n";
    $m ++;

    $cmd  = "plot ";
    $cmd .= "'$unifile4'  using 1:2 title 'Synthetic noss' with boxes ls $m";
    print $gp "$cmd\n";
}

sub gnuplot_uniform_hlx {
    my ($results_dir, $nagg, $aggmethod_ref, $unifile1, $unifile2, $unifile3, $unifile4) = @_;

    my $psfile = "$results_dir/profmark.hlx.uniform.ps";    
    my $pdfile = "$results_dir/profmark.hlx.uniform.pdf";

    open(GP,'|'.GNUPLOT) || die "Gnuplot: $!";
    print GP "set terminal postscript color 14\n";
    print GP "set output '$psfile'\n";
    FUNCS::gnuplot_define_styles (*GP);

    # Observed E-value
    my $ylabel = "Occupancy";
    my $xlabel = "Null helix P-values";
    gnuplot_uniform_hlx_one(*GP, $xlabel, $ylabel, $nagg, $aggmethod_ref, $unifile1, $unifile2, $unifile3, $unifile4);
    
    close(GP);
    
    system("ps2pdf $psfile $pdfile\n");
    system("rm $psfile\n");
    
    if ($viewplot) {
	system("open $pdfile&\n"); 
    }
}

sub gnuplot_uniform_hlx_one {
    my ($gp, $xlabel, $ylabel, $nagg, $aggmethod_ref, $unifile1, $unifile2, $unifile3, $unifile4) = @_;
    
    print $gp "set key right top\n";
    print $gp "set xlabel '$xlabel'\n";
    print $gp "set ylabel '$ylabel'\n";
    print $gp "set xrange [0:1.01]\n";

    my $title = "Rfam";
    gnuplot_uniform_hlx_onecase($gp, $title, $nagg, $aggmethod_ref, $unifile1);
      
    $title = "Rfam noss";
    gnuplot_uniform_hlx_onecase($gp, $title, $nagg, $aggmethod_ref, $unifile2);

    $title = "synthetic";
    gnuplot_uniform_hlx_onecase($gp, $title, $nagg, $aggmethod_ref, $unifile3);
  
    $title = "Synthetic noss";
    gnuplot_uniform_hlx_onecase($gp, $title, $nagg, $aggmethod_ref, $unifile4);    
}

sub gnuplot_uniform_hlx_onecase {
    my ($gp, $title, $nagg, $aggmethod_ref, $unifile_ref) = @_;
    
    print $gp "set title '$title'\n";
    my $cmd;
    my $m = 1;
    for (my $agg = 0; $agg < $nagg-1; $agg ++) {
	$cmd = "plot ";
	$cmd .= "'$unifile_ref->[$agg]' using 1:2 title '$aggmethod_ref->[$agg]' with boxes ls $m, "; 
	print $gp "$cmd\n";
	$m ++;
    }
    $cmd .= "'$unifile_ref->[$nagg-1]' using 1:2 title '$aggmethod_ref->[$nagg-1]' with boxes ls $m";     
    print $gp "$cmd\n";
}

sub gnuplot_ppairs {
    my ($results_dir, $pairsfile1, $pairsfile2, $pairsfile3, $pairsfile4) = @_;

    my $psfile = "$results_dir/profmark.ppairs.ps";    
    my $pdfile = "$results_dir/profmark.ppairs.pdf";
    open(GP,'|'.GNUPLOT) || die "Gnuplot: $!";
    print GP "set terminal postscript color 14\n";
    FUNCS::gnuplot_define_styles (*GP);
    print GP "set output '$psfile'\n";

    my $xlabel = "P-value of two base pairs in same helix";
    print GP "set size square\n";
    print GP "set yrange [0.:1]\n";
    print GP "set xrange [0:1]\n";
    print GP "set xlabel '$xlabel'\n";
 
    my $title  = "Rfam";
    gnuplot_ppairs_one(*GP, $title, $pairsfile1);
    $title  = "R-scape Null";
    gnuplot_ppairs_one(*GP, $title, $pairsfile2);
    $title  = "Synthetic";
    gnuplot_ppairs_one(*GP, $title, $pairsfile3);
    $title  = "Synthetic Null";
    gnuplot_ppairs_one(*GP, $title, $pairsfile4);
    
    close(GP);
    
    system("ps2pdf $psfile $pdfile\n");
    system("rm $psfile\n");
    
    if ($viewplot) {
	system("open $pdfile&\n"); 
    }
}

sub gnuplot_ppairs_one {
    my ($gp, $title, $pairsfile) = @_;
    
    print $gp "set title '$title'\n";
    
    my $cmd = "plot ";
    my $m = 1;
    $cmd .= "'$pairsfile' using 1:2 title ''  with dots ls $m"; 
    print $gp "$cmd\n";
    #print "$cmd";
}

sub gnuplot_hlxstat {
    my ($results_dir, $hlxstatfile1r_ref, $hlxstatfile1n_ref, $hlxstatfile2r_ref, $hlxstatfile2n_ref) = @_;
    
    my $psfile = "$results_dir/profmark.hlx.stats.ps";    
    my $pdfile = "$results_dir/profmark.hlx.stats.pdf";
    open(GP,'|'.GNUPLOT) || die "Gnuplot: $!";
    print GP "set terminal postscript color 14\n";
    FUNCS::gnuplot_define_styles (*GP);
    print GP "set output '$psfile'\n";

    my $ylabel = "occurence";
    print GP "set ylabel '$ylabel'\n";
    print GP "set yrange [0:*]\n";
  
    my $title  = "Rfam";
    gnuplot_hlxstat_one(*GP, $title, $hlxstatfile1r_ref, $hlxstatfile1n_ref);
    $title  = "Synthetic";
    gnuplot_hlxstat_one(*GP, $title, $hlxstatfile2r_ref, $hlxstatfile2n_ref);
    
    close(GP);
    
    system("ps2pdf $psfile $pdfile\n");
    system("rm $psfile\n");
    
    if ($viewplot) {
	system("open $pdfile&\n"); 
    }
}

sub gnuplot_hlxstat_one {
    my ($gp, $title, $hlxstatfiler_ref, $hlxstatfilen_ref) = @_;
    
    print $gp "set title '$title'\n";
    
    my $cmd = "plot ";
    my $m = 1;
    my $xlabel = "helix length";
    print $gp "set xlabel '$xlabel'\n";
    $cmd .= "'$hlxstatfilen_ref->[0]' using 1:2 title 'Null'  with boxes ls $m, "; $m ++;
    $cmd .= "'$hlxstatfiler_ref->[0]' using 1:2 title 'Real'  with boxes ls $m "; 
    print $gp "$cmd\n";
    #print "$cmd";
    
    $cmd = "plot ";
    $m = 1;
    $xlabel = "number of helices";
    print $gp "set xlabel '$xlabel'\n";
    $cmd .= "'$hlxstatfilen_ref->[1]' using 1:2 title 'Null'  with boxes ls $m, "; $m ++;
    $cmd .= "'$hlxstatfiler_ref->[1]' using 1:2 title 'Real'  with boxes ls $m "; 
    print $gp "$cmd\n";
    #print "$cmd";

    $cmd = "plot ";
    $m = 1;
    $xlabel = "fraction of base paired nucleotides";
    print $gp "set xlabel '$xlabel'\n";
    $cmd .= "'$hlxstatfilen_ref->[2]' using 1:2 title 'Null'  with boxes ls $m, "; $m ++;
    $cmd .= "'$hlxstatfiler_ref->[2]' using 1:2 title 'Real'  with boxes ls $m "; 
    print $gp "$cmd\n";
    #print "$cmd";
}

sub parse_covfile {
    my ($rocfile, $unifile, $dir, $string) = @_;

    my $outfile = "$dir/$string.out";
    my $cmd = "$dir/*.$string.out | sort -g";
    system("cat $cmd > $outfile\n");    

    my $fp    = 0;
    my $tp    = 0;
    my $found = 0;
    my $total_true_bp = 0;
    my $total_exp_bp  = 0;
    my $total_msa     = 0;

    my $N = 1;
    my $k = 1000;
    my $shift = 0;
    my @his_pval;
    my @his_pval_null;
    FUNCS::init_histo_array($N, $k, \@his_pval);
    FUNCS::init_histo_array($N, $k, \@his_pval_null);

    my @eachfile;
    my $suffix = "$string.out";
    my $prefix = "RF";
    FUNCS::sorted_files($dir, \@eachfile, $suffix, $prefix);
    my $nf = $#eachfile+1;
    for (my $f = 0; $f < $nf; $f ++) {
	open(FILE, "$eachfile[$f]") || die;
	while (<FILE>)
	{
	    if (/# true_bp\s+(\d+)\s*$/) {
		$total_true_bp += $1;
	    }
	    if (/# exp_bp\s+(\S+)\s*$/) {
		$total_exp_bp += $1;
	    }
	    if (/# nmsa\s+(\S+)\s*$/) {
		$total_msa += $1;
	    }
	}
	close(FILE);
    }
    printf("$rocfile: Total_bp $total_true_bp Exp_bp $total_exp_bp\n");

    open(ROC,  ">$rocfile") || die;
    open(FILE, "$outfile")  || die;
    while (<FILE>)
    {
	if (/^(\S+)\s+(\S+)\s+\S+\s+\d+\s+\d+\s+\S+\s+(\S+)\s*$/) {

	    my $eval   = $1;
	    my $pval   = $2;
	    my $target = $3;

	    if ($target =~ /decoy/) # a false positive
	    {
		$fp++;
		FUNCS::fill_histo_array(1, $pval, $N, $k, $shift, \@his_pval_null);
	    }
	    else { 
		$tp ++;
		FUNCS::fill_histo_array(1, $pval, $N, $k, $shift, \@his_pval);
	    }

	    $found = $fp + $tp;
	    my $fdr = ($found >= 0)? $fp/$found : 0;
	    if ($fp >= 0 && $eval < $eval_cutoff) {
		printf(ROC "%f %f %f %f %d %d %g %g\n", $fp/$total_msa, $fdr, $tp/$total_true_bp, $tp/$total_exp_bp, $fp, $tp, $eval, $pval);
	    }
	}
    }
    printf(ROC "&\n");
	
    close(FILE);
    close(ROC);

    FUNCS::write_histogram($N, $k, $shift, \@his_pval_null, 1, $unifile, 0);

    return $total_msa;
}

sub parse_hlxfile {
    my ($rocfile, $unifile, $nocfile, $dir, $string) = @_;

    my $outfile = "$dir/$string.out";
    
    my $fp    = 0;
    my $tp    = 0;
    my $found = 0;
    my $total_true_hlx = 0;
    my $total_msa      = 0;
    
    my $N = 1;
    my $k = 1000;
    my $shift = 0;
    my @his_pval;
    my @his_pval_null;
    FUNCS::init_histo_array($N, $k, \@his_pval);
    FUNCS::init_histo_array($N, $k, \@his_pval_null);

    my @eachfile;
    my $suffix = "$string.out";
    my $prefix = "RF";
    FUNCS::sorted_files($dir, \@eachfile, $suffix, $prefix);
    my $nf = $#eachfile+1;
    for (my $f = 0; $f < $nf; $f ++) {
	open(FILE, "$eachfile[$f]")  || die;
	while (<FILE>)
	{
	    if (/# true_hlx\s+(\d+)\s*$/) {
		$total_true_hlx += $1;
	    }
	    if (/# nmsa\s+(\S+)\s*$/) {
		$total_msa += $1;
	    }
	}
	close(FILE);
    }

    my $ncvt = 4;
    my @ncv_cutoff;
    $ncv_cutoff[0] = 1;
    $ncv_cutoff[1] = 2;
    $ncv_cutoff[2] = 3;
    $ncv_cutoff[3] = 4;
    my @hlx_ncv_tp;
    my @hlx_ncv_fp;
    $hlx_ncv_tp[0] = 0;
    $hlx_ncv_tp[1] = 0;
    $hlx_ncv_tp[2] = 0;
    $hlx_ncv_tp[3] = 0;
    $hlx_ncv_fp[0] = 0;
    $hlx_ncv_fp[1] = 0;
    $hlx_ncv_fp[2] = 0;
    $hlx_ncv_fp[3] = 0;
     
    my $cmd = "$dir/*.$string.out | sort -g";
    system("cat $cmd > $outfile\n");
    open(ROC,  ">$rocfile") || die;
    open(NOC,  ">$nocfile") || die;
    open(FILE, "$outfile")  || die;
    while (<FILE>)
    {
	if (/^(\S+)\s+(\S+)\s+\S+\s+\S+\s+(\d+)\s+(\d+)\s+\S+\s+(\S+)\s+/) {
	    my $eval   = $1;
	    my $pval   = $2;
	    my $nbp    = $3;
	    my $ncv    = $4;
	    my $target = $5;

	    if ($target =~ /decoy/) # a false positive
	    {
		$fp++;
		FUNCS::fill_histo_array(1, $pval, $N, $k, $shift, \@his_pval_null);
		
		for (my $x = 0; $x < $ncvt; $x ++) {
		    if ($ncv > $ncv_cutoff[$x]) { $hlx_ncv_fp[$x] ++; }
		}
	    }
	    else { 
		$tp ++;
		FUNCS::fill_histo_array(1, $pval, $N, $k, $shift, \@his_pval);
		
		for (my $x = 0; $x < $ncvt; $x ++) {
		    if ($ncv > $ncv_cutoff[$x]) { $hlx_ncv_tp[$x] ++; }
		}
	    }
	    $found = $tp + $fp;

	    my $fdr = ($found >= 0)? $fp/$found : 0;
	    if ($fp >= 0 && $eval < $eval_cutoff) {
		printf(ROC "%f %f %f %d %d %g %g\n", $fp/$total_msa, $fdr, $tp/$total_true_hlx, $fp, $tp, $eval, $pval);
	    }
	}
    }
    printf(ROC "&\n");

    for (my $x = 0; $x < $ncvt; $x ++) {
	printf(NOC "%f %f %d %d\n", $hlx_ncv_fp[$x]/$total_msa, $hlx_ncv_tp[$x]/$total_true_hlx, $hlx_ncv_fp[$x], $hlx_ncv_tp[$x]);
    }
    
    close(FILE);
    close(ROC);
    close(NOC);

    FUNCS::write_histogram($N, $k, $shift, \@his_pval_null, 1, $unifile, 0);

    system("rm $outfile\n");
    return $nf;
}
```
