## Supplementary material for "RNA covariation at helix-level resolution for the identification of evolutionarily conserved RNA structure": profmark_analyze_cov.pl

```
#!/usr/bin/perl -w
# 
# profmark_analyze_cov
#
use strict;
use Class::Struct;
use POSIX;
    
#use constant GNUPLOT => '/usr/local/bin/gnuplot';
use constant GNUPLOT => '/n/helmod/apps/centos7/Core/gnuplot/5.2.6-fasrc01/bin/gnuplot';

#use lib '/Users/erivas/src/Mysrc/R-scape/scripts';
use lib '/n/eddy_lab/users/erivas/rscape_v2.0.0.p/scripts';
use FUNCS;

use vars qw ($opt_v $opt_O $opt_G);  # required if strict used
use Getopt::Std;
getopts ('O:Gv');

# Print a helpful message if the user provides no input file.
if (!@ARGV) {
    print "usage:  profmark_analyze_cov.pl [options] <resultsdir> <which> \n\n";
    print "options:\n";
    print "-v    :  be verbose\n";
    exit;
}

my $results_dir = shift;
my $which       = shift;

my $family = "all";
if ($opt_O) { $family = "$opt_O"; }

system("rm $results_dir/profmark.$family.cov.*\n");

my $string;
if    ($which == 1) { $string = "select.$family.ss.cov";      }
elsif ($which == 2) { $string = "select.$family.noss.cov";    }
elsif ($which == 3) { $string = "synthetic.$family.ss.cov";   }
elsif ($which == 4) { $string = "synthetic.$family.noss.cov"; }
else { print "which has to be 1,2,3,4\n"; exit; }


my $viewplot = 0;
my $plotonly = 0;
if ($opt_G) { $plotonly = 1; }

my $eval_cutoff = 10000;

my $nali_select         = 0;
my $nali_select_noss    = 0;
my $nali_synthetic      = 0;
my $nali_synthetic_noss = 0;
all_parse_covfile($results_dir, $which, $family, \$nali_select, \$nali_select_noss, \$nali_synthetic, \$nali_synthetic_noss);
printf("nali_select= $nali_select/$nali_select_noss nali_synthetic = $nali_synthetic/$nali_synthetic_noss\n");


sub all_parse_covfile {
    my ($results_dir, $which, $family, $ret_nali_select, $ret_nali_select_noss, $ret_nali_synthetic, $ret_nali_synthetic_noss) = @_;

    my $string1   = "select.$family.cov";
    my $string    = "select.cov";
    my $rocfile1  = "$results_dir/$string1.roc";
    my $unifile1r = "$results_dir/$string1.uni";
    my $unifile1n = "$results_dir/$string1.null.uni";
    if ($plotonly == 0 && $which == 1) {
	system("rm $results_dir/$string1.*\n");
	$$ret_nali_select = parse_covfile($rocfile1, $unifile1r, $unifile1n, $results_dir, $string, $string1, $family);
    }
    
    $string1      = "select.noss.$family.cov";
    $string       = "select.noss.cov";
    my $rocfile2  = "$results_dir/$string1.roc";
    my $unifile2r = "$results_dir/$string1.uni";
    my $unifile2n = "$results_dir/$string1.null.uni";
    if ($plotonly == 0 && $which == 2) {
	system("rm $results_dir/$string1.*\n");
	$$ret_nali_select_noss = parse_covfile($rocfile2, $unifile2r, $unifile2n, $results_dir, $string, $string1, $family);
    }
    
    $string1      = "synthetic.$family.cov";
    $string       = "synthetic.cov";
    my $rocfile3  = "$results_dir/$string1.roc";
    my $unifile3r = "$results_dir/$string1.uni";
    my $unifile3n = "$results_dir/$string1.null.uni";
    if ($plotonly == 0  && $which == 3) {
	system("rm $results_dir/$string1.*\n");
	$$ret_nali_synthetic = parse_covfile($rocfile3, $unifile3r, $unifile3n, $results_dir, $string, $string1, $family);
    }
    
    $string1      = "synthetic.noss.$family.cov";
    $string       = "synthetic.noss.cov";
    my $rocfile4  = "$results_dir/$string1.roc";
    my $unifile4r = "$results_dir/$string1.uni";
    my $unifile4n = "$results_dir/$string1.null.uni";
    if ($plotonly == 0 && $which == 4) {
	system("rm $results_dir/$string1.*\n");
	$$ret_nali_synthetic_noss = parse_covfile($rocfile4, $unifile4r, $unifile4n, $results_dir, $string, $string1, $family);
    }

    if ($plotonly) {
	gnuplot_rocplot_cov($results_dir, $family, $rocfile1, $rocfile2, $rocfile3, $rocfile4);
	gnuplot_uniform_cov($results_dir, $family,
			    $unifile1r, $unifile1n, 
			    $unifile2r, $unifile2n, 
			    $unifile3r, $unifile3n, 
			    $unifile4r, $unifile4n);
    }
}


sub eval_from_covroc {
    my ($eval_target, $rocfile) = @_;
    
    my $evalfile = "$rocfile.eval";
    
    open(OUT, ">$evalfile") || die;
    
    open(FILE, "$rocfile") || die;
    while (<FILE>)
    {
	if (/^(\S+)\s+\S+\s+\S+\s+\S+\s+\d+\s+\d+\s+(\S+)\s+\S+\s*$/) {
	    my $oeval = $1;
	    my $eval  = $2;
	    if ($eval > $eval_target) {
		print OUT "$oeval 0\n";
		print OUT "$oeval 1\n";
		last;
	    }
	}
    }
    close(FILE);
    close(OUT);
    
    return $evalfile;
}

sub eval2eval_covroc {
    my ($rocfile) = @_;
    
    my $evalfile = "$rocfile.eval2eval";
    
    open(OUT, ">$evalfile") || die;
    
    open(FILE, "$rocfile") || die;
    while (<FILE>)
    {
	if (/^(\S+)\s+\S+\s+\S+\s+\S+\s+\d+\s+\d+\s+(\S+)\s+\S+\s*$/) {
	    my $oeval = $1;
	    my $eval  = $2;
	    if ($oeval > 0) { print OUT "$oeval $eval\n"; }
	}
    }
    close(FILE);
    close(OUT);
    
    return $evalfile;
}


sub gnuplot_rocplot_cov {
    my ($results_dir, $family, $rocfile1, $rocfile2, $rocfile3, $rocfile4) = @_;
    
    my $psfile = "$results_dir/profmark.$family.cov.roc.ps";    
    my $pdfile = "$results_dir/profmark.$family.cov.roc.pdf";
    open(GP,'|'.GNUPLOT) || die "Gnuplot: $!";
    print GP "set terminal postscript color 14\n";
    FUNCS::gnuplot_define_styles (*GP);
    print GP "set output '$psfile'\n";

    my $xlabel = "mean false positives per alignment (observed E-value)";
    my $which  = 1; # Observed E-value
    my $xmin   = 0.000026;
    my $xmax   = 10;
    my $ymax   = 1;
    gnuplot_rocplot_cov_one(*GP, $which, $xlabel, $xmin, $xmax, $ymax, $rocfile1, $rocfile2, $rocfile3, $rocfile4, 1, 1);
    
    $xlabel = "false discovery rate";
    $which  = 2; # False Discovery Rate;
    $xmin   = 0.000026;
    $xmax   = 1;
    $ymax   = 1;
    gnuplot_rocplot_cov_one(*GP, $which, $xlabel, $xmin, $xmax, $ymax, $rocfile1, $rocfile2, $rocfile3, $rocfile4, 1, 0);

    $xlabel = "false positives";
    $which  = 5; # False positives
    $xmin = 0;
    $xmax = 50;
    $ymax = 0.3;
    gnuplot_rocplot_cov_one(*GP, $which, $xlabel, $xmin, $xmax, $ymax, $rocfile1, $rocfile2, $rocfile3, $rocfile4, 0, 0);

    $xlabel = "false positives";
    $which  = 5; # False positives
    $xmin = 0;
    $xmax = 500;
    $ymax = 1;
    gnuplot_rocplot_cov_one(*GP, $which, $xlabel, $xmin, $xmax, $ymax, $rocfile1, $rocfile2, $rocfile3, $rocfile4, 0, 0);

    close(GP);
    
    system("ps2pdf $psfile $pdfile\n");
    system("rm $psfile\n");
    
    if ($viewplot) {
	system("open $pdfile&\n"); 
    }
    
}

sub gnuplot_rocplot_cov_one {
    my ($gp, $which, $xlabel, $xmin, $xmax, $ymax, $rocfile1, $rocfile2, $rocfile3, $rocfile4, $xlogscale, $doeval) = @_;

    my $evalfile1 = eval_from_covroc(0.05, $rocfile1);
    my $evalfile2 = eval_from_covroc(0.05, $rocfile2);
    my $evalfile3 = eval_from_covroc(0.05, $rocfile3);
    my $evalfile4 = eval_from_covroc(0.05, $rocfile4);
    
    my $eval2evalfile1 = eval2eval_covroc($rocfile1);
    my $eval2evalfile2 = eval2eval_covroc($rocfile2);
    my $eval2evalfile3 = eval2eval_covroc($rocfile3);
    my $eval2evalfile4 = eval2eval_covroc($rocfile4);
    
    print $gp "set xlabel '$xlabel'\n";

    my $ylabel = "Fraction of base pairs that covary"; 
    print $gp "set ylabel '$ylabel'\n";

    print $gp "set key right bottom\n";
    if ($xlogscale) { print $gp "set logscale x\n"; } else { print $gp "unset logscale x\n"; }
    print $gp "set size nosquare\n";
    print $gp "unset logscale y\n";

    print $gp "set yrange [0:$ymax]\n";
    if ($xmin >= 0 && $xmax >= 0) {
	print $gp "set xrange [$xmin:$xmax]\n";
    }
    elsif ($xmax >= 0) {
	print $gp "set xrange [*:$xmax]\n";
    }
    elsif ($xmax >= 0) {
	print $gp "set xrange [$xmin:*]\n";
    }
    else {
	print $gp "set xrange [*:*]\n";
    }
    
    my $cmd = "plot ";
    my $m = 1;
    $cmd .= "'$rocfile1'  using $which:3 title 'Rfam'           with lines ls $m, "; 
    if ($doeval) { $cmd .= "'$evalfile1' using 1:2     title ''                with lines ls $m, "; } 
    $m ++;
    
    $cmd .= "'$rocfile2'  using $which:3 title 'Rfam noss'      with lines ls $m, ";     
    if ($doeval) { $cmd .= "'$evalfile2' using 1:2     title ''                with lines ls $m, "; }      
    $m ++;
    
    $cmd .= "'$rocfile3'  using $which:3 title 'Synthetic'      with lines ls $m, ";       
    if ($doeval) { $cmd .= "'$evalfile3' using 1:2     title ''                with lines ls $m, "; }    
    $m ++;

    if ($doeval) {
	$cmd .= "'$rocfile4'  using $which:3 title 'Synthetic noss' with lines ls $m, ";
	$cmd .= "'$evalfile4' using 1:2     title ''                with lines ls $m";
    }
    else {
	$cmd .= "'$rocfile4'  using $which:3 title 'Synthetic noss' with lines ls $m";
    }
    print $gp "$cmd\n";
    #print "$cmd";

    my $eval_display = 50;
    if ($doeval) {
	$xlabel = "observed E-value";
	$ylabel = "expected E-value";
	print $gp "set logscale x\n";
	print $gp "set logscale y\n";
	print $gp "set size square\n";
	print $gp "set yrange [0.00001:$eval_display]\n";
	print $gp "set xrange [0.00001:$eval_display]\n";
	print $gp "set xlabel '$xlabel'\n";
	print $gp "set ylabel '$ylabel'\n";
	
	$cmd = "plot ";
	$m = 1;
	$cmd .= "x, ";
	$cmd .= "'$eval2evalfile1' using 1:2 title 'Rfam'  with lines ls $m"; 
	print $gp "$cmd\n";
	#print "$cmd";
	
	$cmd = "plot ";
	$m = 1;
	$cmd .= "x, ";
	$cmd .= "'$eval2evalfile2' using 1:2 title 'Rfam-noss'  with lines ls $m"; 
	print $gp "$cmd\n";
	#print "$cmd";
	
	$cmd = "plot ";
	$m = 1;
	$cmd .= "x, ";
	$cmd .= "'$eval2evalfile3' using 1:2 title 'Rfam-Synthetic'  with lines ls $m"; 
	print $gp "$cmd\n";
	#print "$cmd";
	
	$cmd = "plot ";
	$m = 1;
	$cmd .= "x, ";
	$cmd .= "'$eval2evalfile4' using 1:2 title 'Rfam-Synthetic-noss'  with lines ls $m"; 
	print $gp "$cmd\n";
	#print "$cmd";
    }
}


sub gnuplot_uniform_cov {
    my ($results_dir, $family, $unifile1r, $unifile1n, $unifile2r, $unifile2n, $unifile3r, $unifile3n, $unifile4r, $unifile4n) = @_;
    
    my $psfile = "$results_dir/profmark.$family.cov.uniform.ps";    
    my $pdfile = "$results_dir/profmark.$family.cov.uniform.pdf";
    open(GP,'|'.GNUPLOT) || die "Gnuplot: $!";
    print GP "set terminal postscript color 14\n";
    FUNCS::gnuplot_define_styles (*GP);
    print GP "set output '$psfile'\n";

    my $xlabel = "Null base pair P-values";
    gnuplot_uniform_cov_one(*GP, $xlabel, $unifile1n, $unifile2n, $unifile3n, $unifile4n);
    
    $xlabel = "True base pair P-values";
    gnuplot_uniform_cov_one(*GP, $xlabel, $unifile1r, $unifile2r, $unifile3r, $unifile4r);
    close(GP);
    
    system("ps2pdf $psfile $pdfile\n");
    system("rm $psfile\n");
    
    if ($viewplot) {
	system("open $pdfile&\n"); 
    }
}

sub gnuplot_uniform_cov_one {
    my ($gp, $xlabel, $unifile1, $unifile2, $unifile3, $unifile4) = @_;

    print $gp "set xlabel '$xlabel'\n";

    my $ylabel = "Occurrence"; 
    print $gp "set ylabel '$ylabel'\n";

    print $gp "set key right top\n";
    print $gp "set xrange [0:1.01]\n";

    my $cmd = "plot ";
    my $m = 1;
    $cmd .= "'$unifile1'  using 1:2 title 'Rfam'           with boxes ls $m"; 
    print $gp "$cmd\n";
    $m ++;

    $cmd  = "plot ";
    $cmd .= "'$unifile2'  using 1:2 title 'Rfam noss'      with boxes ls $m";     
    print $gp "$cmd\n";
    $m ++;
    
    $cmd  = "plot ";
    $cmd .= "'$unifile3'  using 1:2 title 'Synthetic'      with boxes ls $m";       
    print $gp "$cmd\n";
    $m ++;

    $cmd  = "plot ";
    $cmd .= "'$unifile4'  using 1:2 title 'Synthetic noss' with boxes ls $m";
    print $gp "$cmd\n";
}


sub parse_covfile {
    my ($rocfile, $unifiler, $unifilen, $dir, $string, $string1, $family) = @_;

    my $outfile = "$dir/$string1.out";
    my $cmd = ($family =~ /^all$/)? ("$dir/*.$string.out | sort -g") : ("$dir/$family.$string.out | sort -g"); 
    
    system("cat $cmd > $outfile\n");    

    my $fp    = 0;
    my $tp    = 0;
    my $found = 0;
    my $total_true_bp = 0;
    my $total_exp_bp  = 0;
    my $total_msa     = 0;

    my $N = 1;
    my $k = 100;
    my $shift = 0;
    my @his_pval;
    my @his_pval_null;
    FUNCS::init_histo_array($N, $k, \@his_pval);
    FUNCS::init_histo_array($N, $k, \@his_pval_null);

    my @eachfile;
    my $suffix = "$string.out";
    my $prefix = "RF";
    FUNCS::sorted_files($dir, \@eachfile, $suffix, $prefix);
    my $nf = $#eachfile+1;
    for (my $f = 0; $f < $nf; $f ++) {
	open(FILE, "$eachfile[$f]") || die;
	while (<FILE>)
	{
	    if (/# true_bp\s+(\d+)\s*$/) {
		$total_true_bp += $1;
	    }
	    if (/# exp_bp\s+(\S+)\s*$/) {
		$total_exp_bp += $1;
	    }
	    if (/# nmsa\s+(\S+)\s*$/) {
		$total_msa += $1;
	    }
	}
	close(FILE);
    }
    printf("$rocfile: Total_msa $total_msa Total_bp $total_true_bp Exp_bp $total_exp_bp\n");

    open(ROC,  ">$rocfile") || die;
    open(FILE, "$outfile")  || die;
    while (<FILE>)
    {
	if (/^(\S+)\s+(\S+)\s+\S+\s+\d+\s+\d+\s+\S+\s+(\S+)\s*$/) {

	    my $eval   = $1;
	    my $pval   = $2;
	    my $target = $3;

	    if ($target =~ /decoy/) # a false positive
	    {
		$fp++;
		FUNCS::fill_histo_array(1, $pval, $N, $k, $shift, \@his_pval_null);
	    }
	    else { 
		$tp ++;
		FUNCS::fill_histo_array(1, $pval, $N, $k, $shift, \@his_pval);
	    }

	    $found = $fp + $tp;
	    my $fdr = ($found >= 0)? $fp/$found : 0;
	    if ($fp >= 0 && $eval < $eval_cutoff) {
		printf(ROC "%f %f %f %f %d %d %g %g\n", $fp/$total_msa, $fdr, $tp/$total_true_bp, $tp/$total_exp_bp, $fp, $tp, $eval, $pval);
	    }
	}
    }
    printf(ROC "&\n");
	
    close(FILE);
    close(ROC);

    FUNCS::write_histogram($N, $k, $shift, \@his_pval,      1, $unifiler, 0);
    FUNCS::write_histogram($N, $k, $shift, \@his_pval_null, 1, $unifilen, 0);

    return $total_msa;
}
```
