## Supplementary material for "RNA covariation at helix-level resolution for the identification of evolutionarily conserved RNA structure": profmark_analyze_hlx.pl

```
#!/usr/bin/perl -w
# 
# profmark_analyze_hlx
#
use strict;
use Class::Struct;
use POSIX;
    
#use constant GNUPLOT => '/usr/local/bin/gnuplot';
use constant GNUPLOT => '/n/helmod/apps/centos7/Core/gnuplot/5.2.6-fasrc01/bin/gnuplot';

#use lib '/Users/erivas/src/Mysrc/R-scape/scripts';
use lib '/n/eddy_lab/users/erivas/rscape_v2.0.0.p/scripts';
 use FUNCS;

use vars qw ($opt_v $opt_G $opt_O $opt_Y);  # required if strict used
use Getopt::Std;
getopts ('GO:vY:');

# Print a helpful message if the user provides no input file.
if (!@ARGV) {
    print "usage:  profmark_analyze_hlx.pl [options] <resultsdir>  \n\n";
    print "options:\n";
    print "-v    :  be verbose\n";
    exit;
}

my $results_dir = shift;

my $family = "all";
if ($opt_O) { $family = "$opt_O"; }

my $ymax = 2600;
if ($opt_Y) { $ymax = $opt_Y; }


system("rm $results_dir/profmark.$family.hlx.*\n");
system("rm $results_dir/profmark.$family.ppairs.*\n");


my $viewplot = 0;
my $plotonly = 0;
if ($opt_G) { $plotonly = 1; }

my $eval_cutoff = 1000;

my $nagg = 8;
my @aggmethod;
$aggmethod[2] = "fisher";
$aggmethod[1] = "sidak";
$aggmethod[0] = "lancaster";
 $aggmethod[3] = "wfisher";
$aggmethod[4] = "lancaster_join";
$aggmethod[5] = "wfisher_join";
$aggmethod[6] = "lancaster_double";
$aggmethod[7] = "wfisher_double";
all_parse_hlxfile($ymax, $nagg, \@aggmethod, $results_dir, $family);

all_pvalue_pairs($results_dir, $family);


sub all_parse_hlxfile {
    my ($ymax, $nagg, $aggmethod_ref, $results_dir, $family) = @_;

    my @rocfile1;
    my @rocfile2;
    my @rocfile3;
    my @rocfile4;
    
    my @unifile1;
    my @unifile2;
    my @unifile3;
    my @unifile4;
    
    my @nocfile1;
    my @nocfile2;
    my @nocfile3;
    my @nocfile4;

    my @edgefile1;
    my @edgefile2;
    my @edgefile3;
    my @edgefile4;

    for (my $agg = 0; $agg < $nagg; $agg ++) {
	one_parse_hlxfile($aggmethod_ref->[$agg], $results_dir, $family,
			  \$rocfile1[$agg],  \$rocfile2[$agg],  \$rocfile3[$agg],  \$rocfile4[$agg], 
			  \$unifile1[$agg],  \$unifile2[$agg],  \$unifile3[$agg],  \$unifile4[$agg], 
			  \$nocfile1[$agg],  \$nocfile2[$agg],  \$nocfile3[$agg],  \$nocfile4[$agg], 
			  \$edgefile1[$agg], \$edgefile2[$agg], \$edgefile3[$agg], \$edgefile4[$agg]);
    }

    gnuplot_rocplot_hlx($results_dir, $family,        $nagg, $aggmethod_ref, \@rocfile1, \@rocfile2, \@rocfile3, \@rocfile4, $nocfile1[4], $nocfile2[4], $nocfile3[4], $nocfile4[4]);
    gnuplot_uniform_hlx($results_dir, $family, $ymax, $nagg, $aggmethod_ref, \@unifile1, \@unifile2, \@unifile3, \@unifile4);
}

sub all_pvalue_pairs {
    my ($results_dir, $family) = @_;

    my $string;
    my $cmd;

    $string           = "select";
    my $pairsfile1r   = "$results_dir/$string.$family.ppairs";
    my $pairsfile1n   = "$results_dir/$string.$family.ppairs.null";
    my @hlxstatfile1r;
    $hlxstatfile1r[0] = "$results_dir/$string.$family.hlx.len.stats";
    $hlxstatfile1r[1] = "$results_dir/$string.$family.hlx.nhl.stats";
    $hlxstatfile1r[2] = "$results_dir/$string.$family.hlx.fbp.stats";
    my @hlxstatfile1n;
    $hlxstatfile1n[0] = "$results_dir/$string.$family.hlx.len.stats.null";
    $hlxstatfile1n[1] = "$results_dir/$string.$family.hlx.nhl.stats.null";
    $hlxstatfile1n[2] = "$results_dir/$string.$family.hlx.fbp.stats.null";
    if ($plotonly == 0) {
	my $outfile1    = "$results_dir/$string.$family.hlx.lancaster.unsorted.out";
	$cmd = ($family =~ /^all$/)? "$results_dir/*.$string.hlx.lancaster.out" : "$results_dir/$family.$string.hlx.lancaster.out";
	system("cat $cmd > $outfile1\n");
 
	system("rm $results_dir/$string.$family.ppairs\n");
	pvalue_pairs($pairsfile1r, $pairsfile1n, \@hlxstatfile1r, \@hlxstatfile1n, $outfile1, $string);
	
	#system("rm $outfile1\n");
     }
    
    $string           = "synthetic";
    my $pairsfile2r   = "$results_dir/$string.$family.ppairs";
    my $pairsfile2n   = "$results_dir/$string.$family.ppairs.null";
    my @hlxstatfile2r;
    $hlxstatfile2r[0] = "$results_dir/$string.$family.hlx.len.stats";
    $hlxstatfile2r[1] = "$results_dir/$string.$family.hlx.nhl.stats";
    $hlxstatfile2r[2] = "$results_dir/$string.$family.hlx.fbp.stats";
    my @hlxstatfile2n;
    $hlxstatfile2n[0] = "$results_dir/$string.$family.hlx.len.stats.null";
    $hlxstatfile2n[1] = "$results_dir/$string.$family.hlx.nhl.stats.null";
    $hlxstatfile2n[2] = "$results_dir/$string.$family.hlx.fbp.stats.null";
    if ($plotonly == 0) {
	my $outfile2      = "$results_dir/$string.$family.hlx.lancaster.unsorted.out";
	$cmd = ($family =~ /^all$/)? "$results_dir/*.$string.hlx.lancaster.out" : "$results_dir/$family.$string.hlx.lancaster.out";
	system("cat $cmd > $outfile2\n");
	
	system("rm $results_dir/$string.$family.ppairs\n");
	pvalue_pairs($pairsfile2r, $pairsfile2n, \@hlxstatfile2r, \@hlxstatfile2n, $outfile2, $string);
	
	#system("rm $outfile2\n");    
   }
 
    gnuplot_ppairs ($results_dir, $family, $pairsfile1r,    $pairsfile1n,    $pairsfile2r,    $pairsfile2n);
    gnuplot_hlxstat($results_dir, $family, \@hlxstatfile1r, \@hlxstatfile1n, \@hlxstatfile2r, \@hlxstatfile2n);
}

sub pvalue_pairs {
    my ($pairsfiler, $pairsfilen, $hlxstatfiler_ref, $hlxstatfilen_ref, $outfile, $string) = @_;

   open(PAIRR, ">$pairsfiler")   || die;
   open(PAIRN, ">$pairsfilen")   || die;

    my $N = 500;
    my $k = 1;
    my $shift = 0;
    my @his_len_hlr;
    my @his_len_hln;
    my @his_nhl_per_ssr;
    my @his_nhl_per_ssn;
    my @his_fracbp_per_ssr;
    my @his_fracbp_per_ssn;
    FUNCS::init_histo_array($N, $k, \@his_len_hlr);
    FUNCS::init_histo_array($N, $k, \@his_len_hln);
    FUNCS::init_histo_array($N, $k, \@his_nhl_per_ssr);
    FUNCS::init_histo_array($N, $k, \@his_nhl_per_ssn);

    my $N3 = 1;
    my $k3 = 100;
    FUNCS::init_histo_array($N3, $k3, \@his_fracbp_per_ssr);
    FUNCS::init_histo_array($N3, $k3, \@his_fracbp_per_ssn);

    my $nss_r = 0;
    my $nss_n = 0;
    my $nhl_r = 0;
    my $nhl_n = 0;
    
    my $nbp;
    my $label;
    my $name;
    my $pval1;
    my $pval2;
    my $len_hl;

    my $isreal;
    open(FILE, "$outfile") || die;
    while(<FILE>) {
	if (/^\# nmsa/) {
	    $isreal = 0;
	}
	elsif (/^\# true_hlx/) {
	    $isreal = 1;
	}
	elsif (/^\# SS\s+(\d+)\s+(\d+)\s+(\d+)\s*$/) {
	    my $nhl_per_ss = $1;
	    my $nbp_per_ss = $2;
	    my $len_per_ss = $3;
	    my $fracbp_per_ss = 2*$nbp_per_ss/$len_per_ss;
	    
	    # histogram of nhl_per_ss
	    if ($isreal) { FUNCS::fill_histo_array(1, $nhl_per_ss, $N, $k, $shift, \@his_nhl_per_ssr); $nss_r ++; }
	    else         { FUNCS::fill_histo_array(1, $nhl_per_ss, $N, $k, $shift, \@his_nhl_per_ssn); $nss_n ++; }

	    # histogram of fracbp_per_ss
	    if ($isreal) { FUNCS::fill_histo_array(1, $fracbp_per_ss, $N3, $k3, $shift, \@his_fracbp_per_ssr); }
	    else         { FUNCS::fill_histo_array(1, $fracbp_per_ss, $N3, $k3, $shift, \@his_fracbp_per_ssn); }

	}
	#0.260057 0.260057 5-59 64-118 55 55 RF00001 RF00001_decoy 0.665069 0.723326
	elsif (/^\S+\s+\S+\s+\S+\s+\S+\s+(\d+)\s+\d+\s+(\S+)\s+(\S+)\s+(\S+)\s+(\S+)\s+\S+\s*$/) {
	    $nbp    = $1;
	    $label  = $2;
	    $name   = $3;
	    $pval1  = $4;
	    $pval2  = $5;
	    $len_hl = 2*$nbp;

	    if ($isreal && $name =~ /decoy/) { print "bad assignment for $outfile\n"; exit; }
	    
	    # save the two random pvals
	    if ($isreal) { printf PAIRR "%g %g\n", $pval1, $pval2; }
	    else         { printf PAIRN "%g %g\n", $pval1, $pval2; }

	    # histogram of len_hl
	    if ($isreal) { FUNCS::fill_histo_array(1, $len_hl, $N, $k, $shift, \@his_len_hlr); $nhl_r ++; }
	    else         { FUNCS::fill_histo_array(1, $len_hl, $N, $k, $shift, \@his_len_hln); $nhl_n ++; }

	}
    }
    close(FILE);
    
    close(PAIRR);
    close(PAIRN);

    my $scale_ss_r = ($nss_r > 0)? 1/$nss_r : 1;
    my $scale_ss_n = ($nss_n > 0)? 1/$nss_n : 1;
    my $scale_hl_r = ($nhl_r > 0)? 1/$nhl_r : 1;
    my $scale_hl_n = ($nhl_n > 0)? 1/$nhl_n : 1;
    FUNCS::write_histogram($N,  $k,  $shift, \@his_len_hlr,        $scale_hl_r, $hlxstatfiler_ref->[0], 0, 1);
    FUNCS::write_histogram($N,  $k,  $shift, \@his_nhl_per_ssr,    $scale_ss_r, $hlxstatfiler_ref->[1], 0, 1);
    FUNCS::write_histogram($N3, $k3, $shift, \@his_fracbp_per_ssr, $scale_ss_r, $hlxstatfiler_ref->[2], 0, 1);
    
    FUNCS::write_histogram($N,  $k,  $shift, \@his_len_hln,        $scale_hl_n, $hlxstatfilen_ref->[0], 0, 1);
    FUNCS::write_histogram($N,  $k,  $shift, \@his_nhl_per_ssn,    $scale_ss_n, $hlxstatfilen_ref->[1], 0, 1);
    FUNCS::write_histogram($N3, $k3, $shift, \@his_fracbp_per_ssn, $scale_ss_n, $hlxstatfilen_ref->[2], 0, 1);

    printf("nss_r $nss_r nss_n $nss_n nhl_r $nhl_r nhl_b $nhl_n\n");
}


sub eval_from_hlxroc {
    my ($eval_target, $rocfile) = @_;
    
    my $evalfile = "$rocfile.eval";
    
    open(OUT, ">$evalfile") || die;
    
    open(FILE, "$rocfile") || die;
    while (<FILE>)
    {
	if (/^(\S+)\s+\S+\s+\S+\s+\d+\s+\d+\s+(\S+)\s+\S+\s*$/) {
	    my $oeval = $1;
	    my $eval  = $2;
	    if ($eval > $eval_target) {
		print OUT "$oeval 0\n";
		print OUT "$oeval 1\n";
		last;
	    }
	}
    }
    close(FILE);
    close(OUT);
    
    return $evalfile;
}

sub eval2eval_hlxroc {
    my ($rocfile) = @_;
    
    my $evalfile = "$rocfile.eval2eval";
    
    open(OUT, ">$evalfile") || die;
    
    open(FILE, "$rocfile") || die;
    while (<FILE>)
    {
	if (/^(\S+)\s+\S+\s+\S+\s+\d+\s+\d+\s+(\S+)\s+\S+\s*$/) {
	    my $oeval = $1;
	    my $eval  = $2;
	    if ($oeval > 0) { print OUT "$oeval $eval\n"; }
	}
    }
    close(FILE);
    close(OUT);
    
    return $evalfile;
}

sub one_parse_hlxfile {
    my ($aggmethod, $results_dir, $family,
	$ret_rocfile1,  $ret_rocfile2,  $ret_rocfile3,  $ret_rocfile4, 
	$ret_unifile1,  $ret_unifile2,  $ret_unifile3,  $ret_unifile4, 
	$ret_nocfile1,  $ret_nocfile2,  $ret_nocfile3,  $ret_nocfile4, 
	$ret_edgefile1, $ret_edgefile2, $ret_edgefile3, $ret_edgefile4) = @_;

    my $string1;
    my $string;

    $string1     = "select.$family.hlx".".$aggmethod";
    $string      = "select.hlx".".$aggmethod";
    $$ret_rocfile1  = "$results_dir/$string1.roc";
    $$ret_unifile1  = "$results_dir/$string1.uni";
    $$ret_nocfile1  = "$results_dir/$string1.noc";
    $$ret_edgefile1 = "$results_dir/$string1.edge";
    if ($plotonly == 0) {
	system("rm $results_dir/$string.*\n");
	system("rm $results_dir/$string1.*\n");
	parse_hlxfile($$ret_rocfile1, $$ret_unifile1, $$ret_nocfile1, $$ret_edgefile1, $results_dir, $string, $string1);
    }

    $string1     = "select.noss.$family.hlx".".$aggmethod";
    $string      = "select.noss.hlx".".$aggmethod";
    $$ret_rocfile2  = "$results_dir/$string1.roc";
    $$ret_unifile2  = "$results_dir/$string1.uni";
    $$ret_nocfile2  = "$results_dir/$string1.noc";
    $$ret_edgefile2 = "$results_dir/$string1.edge";
    if ($plotonly == 0) {
	system("rm $results_dir/$string.*\n");
	system("rm $results_dir/$string1.*\n");
	parse_hlxfile($$ret_rocfile2, $$ret_unifile2, $$ret_nocfile2, $$ret_edgefile2, $results_dir, $string, $string1);
    }
    
    $string1     = "synthetic.$family.hlx".".$aggmethod";
    $string      = "synthetic.hlx".".$aggmethod";
    $$ret_rocfile3  = "$results_dir/$string1.roc";
    $$ret_unifile3  = "$results_dir/$string1.uni";
    $$ret_nocfile3  = "$results_dir/$string1.noc";
    $$ret_edgefile3 = "$results_dir/$string1.edge";
    if ($plotonly == 0) {
	system("rm $results_dir/$string.*\n");
	system("rm $results_dir/$string1.*\n");
	parse_hlxfile($$ret_rocfile3, $$ret_unifile3, $$ret_nocfile3, $$ret_edgefile3, $results_dir, $string, $string1);
    }
    
    $string1     = "synthetic.noss.$family.hlx".".$aggmethod";
    $string      = "synthetic.noss.hlx".".$aggmethod";
    $$ret_rocfile4  = "$results_dir/$string1.roc";
    $$ret_unifile4  = "$results_dir/$string1.uni";
    $$ret_nocfile4  = "$results_dir/$string1.noc";
    $$ret_edgefile4 = "$results_dir/$string1.edge";
    if ($plotonly == 0) {
	system("rm $results_dir/$string.*\n");
	system("rm $results_dir/$string1.*\n");
	parse_hlxfile($$ret_rocfile4, $$ret_unifile4, $$ret_nocfile4, $$ret_edgefile4, $results_dir, $string, $string1);
    }
}


sub gnuplot_rocplot_hlx {
    my ($results_dir, $family, $nagg, $aggmethod_ref, $rocfile1_ref, $rocfile2_ref, $rocfile3_ref, $rocfile4_ref, $nocfile1, $nocfile2, $nocfile3, $nocfile4) = @_;

    my $psfile = "$results_dir/profmark.$family.hlx.roc.ps";    
    my $pdfile = "$results_dir/profmark.$family.hlx.roc.pdf";

    open(GP,'|'.GNUPLOT) || die "Gnuplot: $!";
    print GP "set terminal postscript color 14\n";
    print GP "set output '$psfile'\n";
    FUNCS::gnuplot_define_styles (*GP);

    # Observed E-value
    my $ylabel = "Fraction of helices that covary";
    my $xlabel = "False positives per alignment (observd E-value)";
    my $which  = 1; # False Positive Rate
    my $xmin   = -1;
    my $xmax   = 1;
    my $ymin   = 0.0;
    my $ymax   = 1.0;
    gnuplot_rocplot_hlx_one(*GP, $which, $xlabel, $xmin, $xmax, $ymin, $ymax, $ylabel, $nagg, $aggmethod_ref, 
			    $rocfile1_ref, $rocfile2_ref, $rocfile3_ref, $rocfile4_ref, $nocfile1, $nocfile2, $nocfile3, $nocfile4, 1, 1);
    $ymin = 0.55;
    $ymax = 1.00;
    gnuplot_rocplot_hlx_one(*GP, $which, $xlabel, $xmin, $xmax, $ymin, $ymax, $ylabel, $nagg, $aggmethod_ref, 
			    $rocfile1_ref, $rocfile2_ref, $rocfile3_ref, $rocfile4_ref, $nocfile1, $nocfile2, $nocfile3, $nocfile4, 1, 0);

    #FDR
    $xlabel = "False Discovery Rate";
    $which  = 2; # False Discovery Rate
    $xmin = -1;
    $xmax = 1;
    #$xmax = 0.001;
    $ymin = 0.0;
    $ymax = 1.0;
    gnuplot_rocplot_hlx_one(*GP, $which, $xlabel, $xmin, $xmax, $ymin, $ymax, $ylabel, $nagg, $aggmethod_ref, 
			    $rocfile1_ref, $rocfile2_ref, $rocfile3_ref, $rocfile4_ref, $nocfile1, $nocfile2, $nocfile3, $nocfile4, 1, 0);
    $xmax = 0.01;
    $ymin = 0.55;
    $ymax = 1.00;
    gnuplot_rocplot_hlx_one(*GP, $which, $xlabel, $xmin, $xmax, $ymin, $ymax, $ylabel, $nagg, $aggmethod_ref, 
			    $rocfile1_ref, $rocfile2_ref, $rocfile3_ref, $rocfile4_ref, $nocfile1, $nocfile2, $nocfile3, $nocfile4, 1, 0);
  
    #FP
    $xlabel = "False Positives";
    $which  = 4;
    $xmin   = 0;
    $xmax   = 50;
    #$xmax   = 5;
    $ymin = 0.0;
    $ymax = 1.0;
    gnuplot_rocplot_hlx_one(*GP, $which, $xlabel, $xmin, $xmax, $ymin, $ymax, $ylabel, $nagg, $aggmethod_ref, 
			    $rocfile1_ref, $rocfile2_ref, $rocfile3_ref, $rocfile4_ref, $nocfile1, $nocfile2, $nocfile3, $nocfile4, 0, 0);
    $xmax = 25;
    $ymin = 0.55;
    $ymax = 1.00;
    gnuplot_rocplot_hlx_one(*GP, $which, $xlabel, $xmin, $xmax, $ymin, $ymax, $ylabel, $nagg, $aggmethod_ref, 
			    $rocfile1_ref, $rocfile2_ref, $rocfile3_ref, $rocfile4_ref, $nocfile1, $nocfile2, $nocfile3, $nocfile4, 0, 0);
  
    close(GP);
    
    system("ps2pdf $psfile $pdfile\n");
    system("rm $psfile\n");
    
    if ($viewplot) {
	system("open $pdfile&\n"); 
    }
}

sub gnuplot_rocplot_hlx_one {
    my ($gp, $which, $xlabel, $xmin, $xmax, $ymin, $ymax, $ylabel, $nagg, $aggmethod_ref, 
	$rocfile1_ref, $rocfile2_ref, $rocfile3_ref, $rocfile4_ref, $nocfile1, $nocfile2, $nocfile3, $nocfile4,
	$xlogscale, $doeval) = @_;
    
    print $gp "set key right bottom\n";
    if ($xlogscale) { print $gp "set logscale x\n"; } else { print $gp "unset logscale x\n"; }
    
    my $title = "Rfam ss";
    gnuplot_rocplot_hlx_onecase($gp, $title, $which, $xlabel, $xmin, $xmax, $ymin, $ymax, $ylabel, $nagg, $aggmethod_ref, $rocfile1_ref, $nocfile1, $doeval);
      
    $title = "Rfam noss";
    gnuplot_rocplot_hlx_onecase($gp, $title, $which, $xlabel, $xmin, $xmax, $ymin, $ymax, $ylabel, $nagg, $aggmethod_ref, $rocfile2_ref, $nocfile2, $doeval);

    $title = "synthetic ss";
    gnuplot_rocplot_hlx_onecase($gp, $title, $which, $xlabel, $xmin, $xmax, $ymin, $ymax, $ylabel, $nagg, $aggmethod_ref, $rocfile3_ref, $nocfile3, $doeval);
  
    $title = "Synthetic noss";
    gnuplot_rocplot_hlx_onecase($gp, $title, $which, $xlabel, $xmin, $xmax, $ymin, $ymax, $ylabel, $nagg, $aggmethod_ref, $rocfile4_ref, $nocfile4, $doeval);    
}

    
sub gnuplot_rocplot_hlx_onecase {
    my ($gp, $title, $which, $xlabel, $xmin, $xmax, $ymin, $ymax, $ylabel, $nagg, $aggmethod_ref, $rocfile_ref, $nocfile, $doeval) = @_;
    
    my $evalfile;
    my $eval2evalfile;

    $nagg = 4;

    print $gp "set size nosquare\n";
    print $gp "unset logscale y\n";
    print $gp "set title '$title'\n";
    print $gp "set yrange [$ymin:$ymax]\n";
    if    ($xmin >= 0 && $xmax >= 0) { print $gp "set xrange [$xmin:$xmax]\n"; } 
    elsif ($xmin >= 0)               { print $gp "set xrange [$xmin:*]\n"; } 
    elsif ($xmax >= 0)               { print $gp "set xrange [*:$xmax]\n"; } 
    else                             { print $gp "set xrange [*:*]\n"; }
    print $gp "set xlabel '$xlabel'\n";
    print $gp "set ylabel '$ylabel'\n";

    my $cmd = "plot ";
    my $m = 1;
    for (my $agg = 0; $agg < $nagg; $agg ++) {
	$evalfile = eval_from_hlxroc(0.05, $rocfile_ref->[$agg]);

	$cmd .= "'$rocfile_ref->[$agg]' using $which:3 title '$aggmethod_ref->[$agg]' with lines ls $m, "; 
	if ($doeval) { 
	    $cmd .= "'$evalfile' using 1:2 title '' with lines ls $m, "; 
	}
	$m ++;
    }
    $cmd .= "'$nocfile' using $which:3 title 'no agg' ls 9,  "; 
    $cmd .= "'$nocfile' using $which:3 title 'no agg' with lines"; 
    print $gp "$cmd\n";

    if ($doeval) {
	my $xlabel = "observed Aggregated E-value";
	my $ylabel = "expected Aggregated E-value";
	print $gp "set title '$title'\n";
	print $gp "set logscale x\n";
	print $gp "set logscale y\n";
	print $gp "set size square\n";
	print $gp "set yrange [0.00001:10]\n";
	print $gp "set xrange [0.00001:10]\n";
	print $gp "set xlabel '$xlabel'\n";
	print $gp "set ylabel '$ylabel'\n";

	for (my $agg = 0; $agg < $nagg; $agg ++) {
	    $eval2evalfile = eval2eval_hlxroc($rocfile_ref->[$agg]);

	    $cmd = "plot ";
	    $m = 1;
	    $cmd .= "x, ";
	    $cmd .= "'$eval2evalfile' using 1:2 title '$aggmethod_ref->[$agg]'  with lines ls $m"; 
	    print $gp "$cmd\n";
	    #print "$cmd";
	}
    }
}


sub gnuplot_uniform_hlx {
    my ($results_dir, $family, $ymax, $nagg, $aggmethod_ref, $unifile1, $unifile2, $unifile3, $unifile4) = @_;

    my $psfile = "$results_dir/profmark.$family.hlx.uniform.ps";    
    my $pdfile = "$results_dir/profmark.$family.hlx.uniform.pdf";

    open(GP,'|'.GNUPLOT) || die "Gnuplot: $!";
    print GP "set terminal postscript color 14\n";
    print GP "set output '$psfile'\n";
    FUNCS::gnuplot_define_styles (*GP);

    # Observed E-value
    my $ylabel = "Occupancy";
    my $xlabel = "Null helix P-values";
    gnuplot_uniform_hlx_one(*GP, $xlabel, $ylabel, $ymax, $nagg, $aggmethod_ref, $unifile1, $unifile2, $unifile3, $unifile4);
    close(GP);
    
    system("ps2pdf $psfile $pdfile\n");
    system("rm $psfile\n");
    
    if ($viewplot) {
	system("open $pdfile&\n"); 
    }
}

sub gnuplot_uniform_hlx_one {
    my ($gp, $xlabel, $ylabel, $ymax, $nagg, $aggmethod_ref, $unifile1, $unifile2, $unifile3, $unifile4) = @_;
    
    print $gp "set key right top\n";
    print $gp "set xlabel '$xlabel'\n";
    print $gp "set ylabel '$ylabel'\n";
    print $gp "set xrange [0:1.01]\n";
    if ($ymax <= 0) { print $gp "set yrange [0:*]\n"; }
    else            { print $gp "set yrange [0:$ymax]\n"; }

    my $title = "Rfam ss";
    gnuplot_uniform_hlx_onecase($gp, $title, $nagg, $aggmethod_ref, $unifile1);
      
    $title = "Rfam noss";
    gnuplot_uniform_hlx_onecase($gp, $title, $nagg, $aggmethod_ref, $unifile2);

    $title = "synthetic ss";
    gnuplot_uniform_hlx_onecase($gp, $title, $nagg, $aggmethod_ref, $unifile3);
  
    $title = "Synthetic noss";
    gnuplot_uniform_hlx_onecase($gp, $title, $nagg, $aggmethod_ref, $unifile4);    
}

sub gnuplot_uniform_hlx_onecase {
    my ($gp, $title, $nagg, $aggmethod_ref, $unifile_ref) = @_;
    
    print $gp "set title '$title'\n";
    my $cmd;
    my $m = 1;
    for (my $agg = 0; $agg < $nagg-1; $agg ++) {
	$cmd  = "plot ";
	$cmd .= "'$unifile_ref->[$agg]' using 1:2 title '$aggmethod_ref->[$agg]' with boxes ls $m"; 
	$m ++;
	print $gp "$cmd\n";
    }
    $cmd  = "plot ";
    $cmd .= "'$unifile_ref->[$nagg-1]' using 1:2 title '$aggmethod_ref->[$nagg-1]' with boxes ls $m";     
    print $gp "$cmd\n";
}

sub gnuplot_ppairs {
    my ($results_dir, $family, $pairsfile1, $pairsfile2, $pairsfile3, $pairsfile4) = @_;

    my $psfile = "$results_dir/profmark.$family.ppairs.ps";    
    my $pdfile = "$results_dir/profmark.$family.ppairs.pdf";
    open(GP,'|'.GNUPLOT) || die "Gnuplot: $!";
    print GP "set terminal postscript color 14\n";
    FUNCS::gnuplot_define_styles (*GP);
    print GP "set output '$psfile'\n";

    my $min = 1e-30;
    my $xlabel = "P-value of two base pairs in same helix";
    print GP "set size square\n";
    print GP "set yrange [$min:1]\n";
    print GP "set xrange [$min:1]\n";
    print GP "set xlabel '$xlabel'\n";
 
    my $title  = "Rfam";
    gnuplot_ppairs_one(*GP, $title, $pairsfile1);    

    $title  = "Synthetic";
    gnuplot_ppairs_one(*GP, $title, $pairsfile3);

    $title  = "R-scape Null";
    gnuplot_ppairs_one(*GP, $title, $pairsfile2);
    
    $title  = "Synthetic Null";
    gnuplot_ppairs_one(*GP, $title, $pairsfile4); 
        
    close(GP);
    
    system("ps2pdf $psfile $pdfile\n");
    system("rm $psfile\n");
    
    if ($viewplot) {
	system("open $pdfile&\n"); 
    }
}

sub gnuplot_ppairs_one {
    my ($gp, $title, $pairsfile) = @_;
    
    print $gp "set title '$title'\n";
    
    my $cmd = "plot ";
    my $m = 1;
    $cmd .= "'$pairsfile' using 1:2 title ''  with dots ls $m"; 
    print $gp "$cmd\n";
    #print "$cmd";
}

sub gnuplot_hlxstat {
    my ($results_dir, $family, $hlxstatfile1r_ref, $hlxstatfile1n_ref, $hlxstatfile2r_ref, $hlxstatfile2n_ref) = @_;
    
    my $psfile = "$results_dir/profmark.$family.hlx.stats.ps";    
    my $pdfile = "$results_dir/profmark.$family.hlx.stats.pdf";
    open(GP,'|'.GNUPLOT) || die "Gnuplot: $!";
    print GP "set terminal postscript color 14\n";
    FUNCS::gnuplot_define_styles (*GP);
    print GP "set output '$psfile'\n";

    my $ylabel = "occurence";
    print GP "set ylabel '$ylabel'\n";
  
    my $title  = "Rfam";
    gnuplot_hlxstat_one(*GP, $title, "Rfam",      $hlxstatfile1r_ref, "Rfam-null",      $hlxstatfile1n_ref);
    $title  = "Synthetic";
    gnuplot_hlxstat_one(*GP, $title, "Synthetic", $hlxstatfile2r_ref, "Synthetic-null", $hlxstatfile2n_ref);
    $title  = "Rfam/Synthetic";
    gnuplot_hlxstat_one(*GP, $title, "Rfam",      $hlxstatfile1r_ref, "Synthetic",      $hlxstatfile2r_ref);

    $ylabel = "frequency";
    print GP "set ylabel '$ylabel'\n";
    gnuplot_hlxstat_cumulative(*GP, $hlxstatfile1r_ref, $hlxstatfile1n_ref, $hlxstatfile2r_ref, $hlxstatfile2n_ref);
    
    close(GP);
    
    system("ps2pdf $psfile $pdfile\n");
    system("rm $psfile\n");
    
    if ($viewplot) {
	system("open $pdfile&\n"); 
    }
}

sub gnuplot_hlxstat_cumulative {
    my ($gp, $hlxstatfile1r_ref, $hlxstatfile1n_ref, $hlxstatfile2r_ref, $hlxstatfile2n_ref) = @_;

    my $title = "structure comparison";
    print $gp "set title '$title'\n";
    print $gp "set yrange [0:1.01]\n";

    # HELIX LENGTH
    my $cmd = "plot ";
    my $m = 1;
    my $xlabel = "helix length";
    print $gp "set xlabel '$xlabel'\n";
    print $gp "set xrange [1:150]\n";
    $cmd .= "'$hlxstatfile1r_ref->[0]' using 1:3 title 'Rfam'            with lines ls $m, "; $m ++;
    $cmd .= "'$hlxstatfile2r_ref->[0]' using 1:3 title 'Synthetic'       with lines ls $m"; 
    print $gp "$cmd\n";
    #print "$cmd";

    $cmd = "plot ";
    $m = 1;
    $cmd .= "'$hlxstatfile1r_ref->[0]' using 1:3 title 'Rfam'            with lines ls $m, "; $m ++;
    $cmd .= "'$hlxstatfile2r_ref->[0]' using 1:3 title 'Synthetic'       with lines ls $m, "; $m ++;
    $cmd .= "'$hlxstatfile1n_ref->[0]' using 1:3 title 'null'            with lines ls $m, "; $m ++;
    $cmd .= "'$hlxstatfile2n_ref->[0]' using 1:3 title 'null-synthetic'  with lines ls $m "; 
    print $gp "$cmd\n";
    #print "$cmd";

    # NUMBER OF HELICES PER STRUCTURE
    $cmd = "plot ";
    $m = 1;
    $xlabel = "number of helices per structure";
    print $gp "set xlabel '$xlabel'\n";
    print $gp "set xrange [0:40]\n";
    $cmd .= "'$hlxstatfile1r_ref->[1]' using 1:3 title 'Rfam'            with lines ls $m, "; $m ++;
    $cmd .= "'$hlxstatfile2r_ref->[1]' using 1:3 title 'Synthetic'       with lines ls $m"; 
    print $gp "$cmd\n";
    #print "$cmd";

    $cmd = "plot ";
    $m = 1;
    $cmd .= "'$hlxstatfile1r_ref->[1]' using 1:3 title 'Rfam'            with lines ls $m, "; $m ++;
    $cmd .= "'$hlxstatfile2r_ref->[1]' using 1:3 title 'Synthetic'       with lines ls $m, "; $m ++;
    $cmd .= "'$hlxstatfile1n_ref->[1]' using 1:3 title 'null'            with lines ls $m, "; $m ++;
    $cmd .= "'$hlxstatfile2n_ref->[1]' using 1:3 title 'null-synthetic'  with lines ls $m "; 
    print $gp "$cmd\n";
    #print "$cmd";

    # AVG #of base pairs per nucleotide
    $cmd = "plot ";
    $m = 1;
    $xlabel = "average number of base pairs per nucleotide";
    print $gp "set xlabel '$xlabel'\n";
    print $gp "set xrange [0:1.01]\n";
    $cmd .= "'$hlxstatfile1r_ref->[2]' using 1:3 title 'Rfam'            with lines ls $m, "; $m ++;
    $cmd .= "'$hlxstatfile2r_ref->[2]' using 1:3 title 'Sythetic'        with lines ls $m"; 
    print $gp "$cmd\n";
    #print "$cmd";
    
    $cmd = "plot ";
    $m = 1;
    $cmd .= "'$hlxstatfile1r_ref->[2]' using 1:3 title 'Rfam'            with lines ls $m, "; $m ++;
    $cmd .= "'$hlxstatfile2r_ref->[2]' using 1:3 title 'Sythetic'        with lines ls $m, "; $m ++;
    $cmd .= "'$hlxstatfile1n_ref->[2]' using 1:3 title 'null'            with lines ls $m, "; $m ++;
    $cmd .= "'$hlxstatfile2n_ref->[2]' using 1:3 title 'null-synthetic'  with lines ls $m "; 
    print $gp "$cmd\n";
    #print "$cmd";
}

sub gnuplot_hlxstat_one {
    my ($gp, $title, $name1, $hlxstatfile1_ref, $name2, $hlxstatfile2_ref) = @_;
    
    print $gp "set title '$title'\n";
    
    my $cmd = "plot ";
    my $m = 1;
    my $xlabel = "helix length";
    print $gp "set xlabel '$xlabel'\n";
    print $gp "set xrange [0:*]\n";
    print $gp "set yrange [0:*]\n";
    $cmd .= "'$hlxstatfile2_ref->[0]' using 1:2 title '$name2'  with boxes ls $m, "; $m ++;
    $cmd .= "'$hlxstatfile1_ref->[0]' using 1:2 title '$name1'  with boxes ls $m "; 
    print $gp "$cmd\n";
    #print "$cmd";
    
    $cmd = "plot ";
    $m = 1;
    $xlabel = "number of helices";
    print $gp "set xlabel '$xlabel'\n";
    print $gp "set xrange [0:*]\n";
    $cmd .= "'$hlxstatfile2_ref->[1]' using 1:2 title '$name2'  with boxes ls $m, "; $m ++;
    $cmd .= "'$hlxstatfile1_ref->[1]' using 1:2 title '$name1'  with boxes ls $m "; 
    print $gp "$cmd\n";
    #print "$cmd";

    $cmd = "plot ";
    $m = 1;
    $xlabel = "average number of base pairs per nucleotide";
    print $gp "set xlabel '$xlabel'\n";
    print $gp "set yrange [0:*]\n";
    print $gp "set xrange [0:1.01]\n";
    $cmd .= "'$hlxstatfile2_ref->[2]' using 1:2 title '$name2'  with boxes ls $m, "; $m ++;
    $cmd .= "'$hlxstatfile1_ref->[2]' using 1:2 title '$name1'  with boxes ls $m "; 
    print $gp "$cmd\n";
    #print "$cmd";
}


sub parse_hlxfile {
    my ($rocfile, $unifile, $nocfile, $edgefile, $dir, $string, $string1) = @_;

    my $outfile = "$dir/$string1.out";
    
    my $fp    = 0;
    my $tp    = 0;
    my $found = 0;
    my $total_true_hlx = 0;
    my $total_msa      = 0;
    
    my $N = 1;
    my $k = 100;
    my $shift = 0;
    my @his_pval;
    my @his_pval_null;
    FUNCS::init_histo_array($N, $k, \@his_pval);
    FUNCS::init_histo_array($N, $k, \@his_pval_null);

    my @eachfile;
    my $suffix = "$string.out";
    my $prefix = "RF";
    FUNCS::sorted_files($dir, \@eachfile, $suffix, $prefix);
    my $nf = $#eachfile+1;
    for (my $f = 0; $f < $nf; $f ++) {
	open(FILE, "$eachfile[$f]")  || die;
	while (<FILE>)
	{
	    if (/# true_hlx\s+(\d+)\s*$/) {
		$total_true_hlx += $1;
	    }
	    if (/# nmsa\s+(\S+)\s*$/) {
		$total_msa += $1;
	    }
	}
	close(FILE);
    }

    my $ncvt = 4;
    my @ncv_cutoff;
    $ncv_cutoff[0] = 1;
    $ncv_cutoff[1] = 2;
    $ncv_cutoff[2] = 3;
    $ncv_cutoff[3] = 4;
    my @hlx_ncv_tp;
    my @hlx_ncv_fp;
    my @hlx_ncv_fdr;
    $hlx_ncv_tp[0]  = 0;
    $hlx_ncv_tp[1]  = 0;
    $hlx_ncv_tp[2]  = 0;
    $hlx_ncv_tp[3]  = 0;
    $hlx_ncv_fp[0]  = 0;
    $hlx_ncv_fp[1]  = 0;
    $hlx_ncv_fp[2]  = 0;
    $hlx_ncv_fp[3]  = 0;
    $hlx_ncv_fdr[0] = 0;
    $hlx_ncv_fdr[1] = 0;
    $hlx_ncv_fdr[2] = 0;
    $hlx_ncv_fdr[3] = 0;

    my $n_plus  = 0;
    my $n_minus = 0;
    my $n_one   = 0;
    my $n_less  = 0;
    
    my $cmd = ($family =~ /^all$/)? ("$dir/*.$string.out | sort -g") : ("$dir/$family.$string.out | sort -g");
    system("cat $cmd > $outfile\n");
    open(ROC,  ">$rocfile")  || die;
    open(NOC,  ">$nocfile")  || die;
    open(EDGE, ">$edgefile") || die;
    open(FILE, "$outfile")   || die;
    while (<FILE>)
    {
	if (/^(\S+)\s+(\S+)\s+\S+\s+\S+\s+(\d+)\s+(\d+)\s+\S+\s+(\S+)\s+\S+\s+\S+\s+(\S+)\s*$/) {
	    my $eval    = $1;
	    my $pval    = $2;
	    my $nbp     = $3;
	    my $ncv     = $4;
	    my $target  = $5;
	    my $minpval = $6;

	    if ($target =~ /decoy/) # a false positive
	    {
		$fp++;
		FUNCS::fill_histo_array(1, $pval, $N, $k, $shift, \@his_pval_null);
		
		for (my $x = 0; $x < $ncvt; $x ++) {
		    if ($ncv >= $ncv_cutoff[$x]) { $hlx_ncv_fp[$x] ++; }
		}
	    }
	    else { 
		$tp ++;
		FUNCS::fill_histo_array(1, $pval, $N, $k, $shift, \@his_pval);
		
		for (my $x = 0; $x < $ncvt; $x ++) {
		    if ($ncv >= $ncv_cutoff[$x]) { $hlx_ncv_tp[$x] ++; }
		}

		# for Figure 1
		if ($ncv == 0 && $eval <= 0.05) {
		    printf(EDGE "++$string1>$_");
		    printf("++$string1>$_");
		    $n_plus ++;
		}
		if ($ncv == 1 && $eval <= 0.05) {
		    printf(EDGE "11$string1>$_");
		    $n_one ++;
		}
		if ($ncv > 0 && $eval > 0.05) {
		    printf(EDGE "--$string1>$_");
		    $n_minus ++;
		}
		if (($ncv == 1 || $nbp-$ncv > 6) && $eval <= $nbp*$minpval && $eval <= 0.05 && $nbp*$minpval > 0.05) {
		    printf(EDGE "<<$string1>$_");
		    $n_less ++;
		}
		
	    }
	    $found = $tp + $fp;
	    
	    for (my $x = 0; $x < $ncvt; $x ++) {
		if ($ncv >= $ncv_cutoff[$x]) {
		    my $found_ncv = $hlx_ncv_fp[$x] + $hlx_ncv_tp[$x];
		    $hlx_ncv_fdr[$x] = ($found_ncv > 0)? $hlx_ncv_fp[$x] / $found : 0; 
		}
	    }

	    my $fdr = ($found >= 0)? $fp/$found : 0;
	    if ($fp >= 0 && $eval < $eval_cutoff) {
		printf(ROC "%g %g %g %d %d %g %g\n", $fp/$total_msa, $fdr, $tp/$total_true_hlx, $fp, $tp, $eval, $pval);
	    }
	}
    }
    printf(ROC "&\n");
    printf(EDGE "n++ $n_plus %f n-- $n_minus %f n11 $n_one %f n<< $n_less %f total $tp\n", $n_plus/$tp, $n_minus/$tp, $n_one/$tp, $n_less/$tp);

    for (my $x = 0; $x < $ncvt; $x ++) {
	printf(NOC "%g %g %g %d %d\n", $hlx_ncv_fp[$x]/$total_msa, $hlx_ncv_fdr[$x], $hlx_ncv_tp[$x]/$total_true_hlx, $hlx_ncv_fp[$x], $hlx_ncv_tp[$x]);
    }
    
    close(FILE);
    close(ROC);
    close(NOC);
    close(EDGE);

    FUNCS::write_histogram($N, $k, $shift, \@his_pval_null, 1, $unifile, 0, 0);

    #system("rm $outfile\n");
    return $nf;
}
```
