## Supplementary material for "RNA covariation at helix-level resolution for the identification of evolutionarily conserved RNA structure": profmark_analyze_msa.pl

```
#!/usr/bin/perl -w
# 
# profmark_analyze_msa
#
use strict;
use Class::Struct;
use POSIX;
    
#use constant GNUPLOT => '/usr/local/bin/gnuplot';
use constant GNUPLOT => '/n/helmod/apps/centos7/Core/gnuplot/5.2.6-fasrc01/bin/gnuplot';

#use lib '/Users/erivas/src/Mysrc/R-scape/scripts';
use lib '/n/eddy_lab/users/erivas/rscape_v2.0.0.p/scripts';
use FUNCS;

use vars qw ($opt_v);  # required if strict used
use Getopt::Std;
getopts ('v');

# Print a helpful message if the user provides no input file.
if (!@ARGV) {
    print "usage:  profmark_analyze_msa.pl [options] <profmarkdir> <results> <rscapedir> \n\n";
    print "options:\n";
    print "-v    :  be verbose\n";
    exit;
}

my $profmark_dir = shift;
my $results      = shift;
my $rscapedir    = shift;
my $results_dir  = "$profmark_dir/$results";

my $viewplot = 0;
my $verbose  = 0;
if ($opt_v) { $verbose = 1; }

system("rm $results_dir/profmark.msastats.*\n");

my $rfam_dir          = "$profmark_dir/select";
my $rfamnull_dir      = "$profmark_dir/select_R-scape_null";
my $synthetic_dir     = "$profmark_dir/select_synthetic";
my $syntheticnull_dir = "$profmark_dir/select_synthetic_null";

my $rfam_tag          = "seed.sto";
my $rfamnull_tag      = "null.sto";
my $synthetic_tag     = "synthetic.sto";
my $syntheticnull_tag = "synthetic.null.sto";

my $esl_alistat = "$rscapedir/lib/hmmer/easel/miniapps/esl-alistat";
my $esl_seqstat = "$rscapedir/lib/hmmer/easel/miniapps/esl-seqstat";

my $Na = 10000;
my $ka = 5;
my $Ni = 100;
my $ki = 10;
my $shift = 0;

my $file_alen_rfam = "$rfam_dir/$rfam_tag.msa.alen";
my $file_slen_rfam = "$rfam_dir/$rfam_tag.msa.slen";
my $file_id_rfam   = "$rfam_dir/$rfam_tag.msa.id";
my $file_gc_rfam   = "$rfam_dir/$rfam_tag.msa.gc";

my $file_alen_rfamnull = "$rfamnull_dir/$rfamnull_tag.msa.alen";
my $file_slen_rfamnull = "$rfamnull_dir/$rfamnull_tag.msa.slen";
my $file_id_rfamnull   = "$rfamnull_dir/$rfamnull_tag.msa.id";
my $file_gc_rfamnull   = "$rfamnull_dir/$rfamnull_tag.msa.gc";
	
my $file_alen_synthetic = "$synthetic_dir/$synthetic_tag.msa.alen";
my $file_slen_synthetic = "$synthetic_dir/$synthetic_tag.msa.slen";
my $file_id_synthetic   = "$synthetic_dir/$synthetic_tag.msa.id";
my $file_gc_synthetic   = "$synthetic_dir/$synthetic_tag.msa.gc";

my $file_alen_syntheticnull = "$syntheticnull_dir/$syntheticnull_tag.msa.alen";
my $file_slen_syntheticnull = "$syntheticnull_dir/$syntheticnull_tag.msa.slen";
my $file_id_syntheticnull   = "$syntheticnull_dir/$syntheticnull_tag.msa.id";
my $file_gc_syntheticnull   = "$syntheticnull_dir/$syntheticnull_tag.msa.gc";
	
my $msastatsfile = "$results_dir/profmark.msastats";
system("rm $msastatsfile\n");
print "output: $msastatsfile\n";
open(OUTF, ">$msastatsfile") || die;

parse_msafiles($rfam_dir,          $rfam_tag,          $file_alen_rfam,          $file_slen_rfam,          $file_id_rfam,          $file_gc_rfam);
parse_msafiles($rfamnull_dir,      $rfamnull_tag,      $file_alen_rfamnull,      $file_slen_rfamnull,      $file_id_rfamnull,      $file_gc_rfamnull);
parse_msafiles($synthetic_dir,     $synthetic_tag,     $file_alen_synthetic,     $file_slen_synthetic,     $file_id_synthetic,     $file_gc_synthetic);
parse_msafiles($syntheticnull_dir, $syntheticnull_tag, $file_alen_syntheticnull, $file_slen_syntheticnull, $file_id_syntheticnull, $file_gc_syntheticnull);

close(OUTF);

gnuplot_msa_histo($msastatsfile);

sub gnuplot_msa_histo {
    my ($statsfile) = @_;

    my $psfile = "$statsfile.ps";    
    my $pdfile = "$statsfile.pdf";

    open(GP,'|'.GNUPLOT) || die "Gnuplot: $!";
    print GP "set terminal postscript color 14\n";
    print GP "set output '$psfile'\n";
    FUNCS::gnuplot_define_styles (*GP);

    my $ylabel = "frequency";
    print GP "set ylabel '$ylabel'\n";
    
    my $m = 1;
    my $xlabel = "alignment length";
    my $xmax = 1000;
    print GP "set xlabel '$xlabel'\n";
    print GP "set xrange [0:$xmax]\n";
    my $cmd = "plot ";
    $cmd   .= "'$file_alen_rfam'          using 1:3 title 'Rfam'           with lines ls $m, "; $m ++;
    $cmd   .= "'$file_alen_rfamnull'      using 1:3 title 'Rfam-null'      with lines ls $m, "; $m ++;
    $cmd   .= "'$file_alen_synthetic'     using 1:3 title 'Synthetic'      with lines ls $m, "; $m ++;
    $cmd   .= "'$file_alen_syntheticnull' using 1:3 title 'Synthetic-null' with lines ls $m";
    print GP "$cmd\n";

    $m = 1;
    $cmd = "plot ";
    $cmd   .= "'$file_alen_rfam'          using 1:2 title 'Rfam'           with boxes ls $m, "; $m ++;
    $cmd   .= "'$file_alen_rfamnull'      using 1:2 title 'Rfam-null'      with boxes ls $m, "; $m ++;
    $cmd   .= "'$file_alen_synthetic'     using 1:2 title 'Synthetic'      with boxes ls $m, "; $m ++;
    $cmd   .= "'$file_alen_syntheticnull' using 1:2 title 'Synthetic-null' with boxes ls $m";
    print GP "$cmd\n";

    $m = 1;
    $xlabel = "average length";
    $xmax = 1000;
    print GP "set xlabel '$xlabel'\n";
    print GP "set xrange [0:$xmax]\n";
    $cmd = "plot ";
    $cmd   .= "'$file_slen_rfam'          using 1:3 title 'Rfam'           with lines ls $m, "; $m ++;
    $cmd   .= "'$file_slen_rfamnull'      using 1:3 title 'Rfam-null'      with lines ls $m, "; $m ++;
    $cmd   .= "'$file_slen_synthetic'     using 1:3 title 'Synthetic'      with lines ls $m, "; $m ++;
    $cmd   .= "'$file_slen_syntheticnull' using 1:3 title 'Synthetic-null' with lines ls $m";
    print GP "$cmd\n";
    
    $m = 1;
    $cmd = "plot ";
    $cmd   .= "'$file_slen_rfam'          using 1:2 title 'Rfam'           with boxes ls $m, "; $m ++;
    $cmd   .= "'$file_slen_rfamnull'      using 1:2 title 'Rfam-null'      with boxes ls $m, "; $m ++;
    $cmd   .= "'$file_slen_synthetic'     using 1:2 title 'Synthetic'      with boxes ls $m, "; $m ++;
    $cmd   .= "'$file_slen_syntheticnull' using 1:2 title 'Synthetic-null' with boxes ls $m";
    print GP "$cmd\n";

    $m = 1;
    $xlabel = "percentage ID";
    $xmax = 100;
    print GP "set xlabel '$xlabel'\n";
    print GP "set xrange [0:$xmax]\n";
    $cmd = "plot ";
    $cmd   .= "'$file_id_rfam'          using 1:3 title 'Rfam'           with lines ls $m, "; $m ++;
    $cmd   .= "'$file_id_rfamnull'      using 1:3 title 'Rfam-null'      with lines ls $m, "; $m ++;
    $cmd   .= "'$file_id_synthetic'     using 1:3 title 'Synthetic'      with lines ls $m, "; $m ++;
    $cmd   .= "'$file_id_syntheticnull' using 1:3 title 'Synthetic-null' with lines ls $m";
    print GP "$cmd\n";

    $m = 1;
    $cmd = "plot ";
    $cmd = "plot ";
    $cmd   .= "'$file_id_rfam'          using 1:2 title 'Rfam'           with boxes ls $m, "; $m ++;
    $cmd   .= "'$file_id_rfamnull'      using 1:2 title 'Rfam-null'      with boxes ls $m, "; $m ++;
    $cmd   .= "'$file_id_synthetic'     using 1:2 title 'Synthetic'      with boxes ls $m, "; $m ++;
    $cmd   .= "'$file_id_syntheticnull' using 1:2 title 'Synthetic-null' with boxes ls $m";
    print GP "$cmd\n";
    
    $m = 1;
    $xlabel = "percentage GC";
    $xmax = 100;
    print GP "set xlabel '$xlabel'\n";
    print GP "set xrange [0:$xmax]\n";
    $cmd = "plot ";
    $cmd   .= "'$file_gc_rfam'          using 1:3 title 'Rfam'           with lines ls $m, "; $m ++;
    $cmd   .= "'$file_gc_rfamnull'      using 1:3 title 'Rfam-null'      with lines ls $m, "; $m ++;
    $cmd   .= "'$file_gc_synthetic'     using 1:3 title 'Synthetic'      with lines ls $m, "; $m ++;
    $cmd   .= "'$file_gc_syntheticnull' using 1:3 title 'Synthetic-null' with lines ls $m";
    print GP "$cmd\n";

    $m = 1;
    $cmd = "plot ";
    $cmd = "plot ";
    $cmd   .= "'$file_gc_rfam'          using 1:2 title 'Rfam'           with boxes ls $m, "; $m ++;
    $cmd   .= "'$file_gc_rfamnull'      using 1:2 title 'Rfam-null'      with boxes ls $m, "; $m ++;
    $cmd   .= "'$file_gc_synthetic'     using 1:2 title 'Synthetic'      with boxes ls $m, "; $m ++;
    $cmd   .= "'$file_gc_syntheticnull' using 1:2 title 'Synthetic-null' with boxes ls $m";
    print GP "$cmd\n";
    
    close(GP);
    
    system("ps2pdf $psfile $pdfile\n");
    system("rm $psfile\n");
    
    if ($viewplot) {
	system("open $pdfile&\n"); 
    }
    
}

sub parse_msafiles {
    my ($msadir, $msatag, $alenfile, $slenfile, $idfile, $gcfile) = @_;

    my @his_alen;
    my @his_slen;
    my @his_id;
    my @his_gc;
    FUNCS::init_histo_array($Na, $ka, \@his_alen);
    FUNCS::init_histo_array($Na, $ka, \@his_slen);
    FUNCS::init_histo_array($Ni, $ki, \@his_id);
    FUNCS::init_histo_array($Ni, $ki, \@his_gc);
    my @msafile;
    my $prefix = "RF";
    my $suffix = $msatag;

    my $outfile = "$msadir/$msatag.out";
    print("outfile:$outfile\n");
    system("rm $outfile");
    
    FUNCS::sorted_files($msadir, \@msafile, $suffix, $prefix);
    my $nf = $#msafile+1;
    print OUTF "\nmsadir: $msadir\n";
    print OUTF "#msafiles $nf\n";
    
    for (my $f = 0; $f < $nf; $f ++) {
	my $cmd = "$esl_alistat $msafile[$f] >> $outfile";
	#print "$cmd\n";
	system("$cmd\n");

	printf OUTF "%d> %s\n", $f+1, $msafile[$f];
	$cmd = "grep -c STOCKHOLM $msafile[$f] > foo";
	system("$cmd\n");

	my $nmsa;
	open(FILE, "foo")  || die;
	while (<FILE>)
	{
	    if (/(\d+)/) { $nmsa = $1; print OUTF "$nmsa\n"; }
	}
	close(FILE);
	system("rm foo\n");

	# for GC
	$cmd = "$esl_seqstat -c $msafile[$f] >> $outfile";
	#print "$cmd\n";
	system("$cmd\n");
    }
    
    parse_alistats($outfile, \@his_alen, \@his_slen, \@his_id, \@his_gc, $alenfile, $slenfile, $idfile, $gcfile);	
}

sub parse_alistats {
    my ($statsfile, $his_alen, $his_slen, $his_id, $his_gc, $alenfile, $slenfile, $idfile, $gcfile) = @_;

    my $alen;
    my $slen;
    my $id;
    my $gc;
    
    my $nmsa = 0;
    my $nmsa_block = 0;
    open(FILE, "$statsfile") || die;
    while(<FILE>) {
	if (/Alignment length:\s+(\S+)/) {
	    $nmsa ++;
	    $alen = $1;
	    FUNCS::fill_histo_array(1, $alen, $Na, $ka, $shift, $his_alen);
	}
	elsif (/Average length:\s+(\S+)\s*$/) {
	    $slen = $1;
	    FUNCS::fill_histo_array(1, $slen, $Na, $ka, $shift, $his_slen);
	}
	elsif (/Average identity:\s+(\d+)\%/) {
	    $id = $1;
	    FUNCS::fill_histo_array(1, $id, $Ni, $ki, $shift, $his_id);
	}
	elsif (/residue: C\s+\d+\s+(\S+)\s*/) {
	    $nmsa_block ++;
	    $gc = $1 * 100;
	}
  	elsif (/residue: G\s+\d+\s+(\S+)\s*/) {
	    $gc += $1 * 100;
	    FUNCS::fill_histo_array(1, $gc, $Ni, $ki, $shift, $his_gc);
	}
    }
    close(FILE);
    print OUTF "nmsa = $nmsa\n";
	   
    FUNCS::write_histogram($Na, $ka, $shift, $his_alen, 1/$nmsa,       $alenfile, 0, 1);
    FUNCS::write_histogram($Na, $ka, $shift, $his_slen, 1/$nmsa,       $slenfile, 0, 1);
    FUNCS::write_histogram($Ni, $ki, $shift, $his_id,   1/$nmsa,       $idfile,   0, 1);
    FUNCS::write_histogram($Ni, $ki, $shift, $his_gc,   1/$nmsa_block, $gcfile,   0, 1);
}
```
