## Supplementary material for "RNA covariation at helix-level resolution for the identification of evolutionarily conserved RNA structure": profmark_analyze_power.pl

```
#!/usr/bin/perl -w
# 
# profmark_analyze_power
#
use strict;
use Class::Struct;
use POSIX;
    
#use constant GNUPLOT => '/usr/local/bin/gnuplot';
use constant GNUPLOT => '/n/helmod/apps/centos7/Core/gnuplot/5.2.6-fasrc01/bin/gnuplot';

#use lib '/Users/erivas/src/Mysrc/R-scape/scripts';
use lib '/n/eddy_lab/users/erivas/rscape_v2.0.0.p/scripts';
use FUNCS;

use vars qw ($opt_v);  # required if strict used
use Getopt::Std;
getopts ('v');

# Print a helpful message if the user provides no input file.
if (!@ARGV) {
    print "usage:  profmark_analyze_power.pl [options] <profmarkdir> <results> <aggtype> \n\n";
    print "options:\n";
    print "-v    :  be verbose\n";
    exit;
}

my $profmark_dir = shift;
my $results      = shift;
my $aggtype      = shift;
my $results_dir  = "$profmark_dir/$results";

my $viewplot = 0;
my $verbose  = 0;
if ($opt_v) { $verbose = 1; }

system("rm $results_dir/profmark.power.*\n");
my $outf = "$results_dir/profmark.power";

my $rfam_dir          = "$profmark_dir/select";
my $rfamnull_dir      = "$profmark_dir/select_R-scape_null";
my $synthetic_dir     = "$profmark_dir/select_synthetic";
my $syntheticnull_dir = "$profmark_dir/select_synthetic_null";

my $nf;
my @outfile;
parse_powerfiles($rfam_dir, $results_dir, \$nf, \@outfile);
#parse_powerfiles($rfamnull_dir, $results_dir, \$nf, \@outfile);
#parse_powerfiles($synthetic_dir, $results_dir, \$nf, \@outfile);
#parse_powerfiles($syntheticnull_dir, $results_dir, \$nf, \@outfile);

gnuplot_power($nf, \@outfile, $results_dir, $outf);

sub gnuplot_power {
    my ($nf, $outfile_ref, $results_dir, $outf) = @_;

    my $psfile = "$outf.ps";    
    my $pdfile = "$outf.pdf";

    open(GP,'|'.GNUPLOT) || die "Gnuplot: $!";
    print GP "set terminal postscript color 14\n";
    print GP "set output '$psfile'\n";
    FUNCS::gnuplot_define_styles (*GP);

    my $ylabel = "expected covariation";
    print GP "set ylabel '$ylabel'\n";
    
    my $m = 1;
    my $xlabel = "observed covariation";
    my $xmax = 3000;
    print GP "set xlabel '$xlabel'\n";
    print GP "set xrange [0:*]\n";
    my $cmd = "plot ";
    my $f;
    for ($f = 0; $f < $nf-1; $f ++) {
	$cmd   .= "'$outfile_ref->[$f]' using 1:2 title '' with lines ls $m, ";
    }
    $cmd   .= "'$outfile_ref->[$f]' using 1:2 title '' with lines ls $m";
    print GP "$cmd\n";

    $m = 1;
    $cmd = "plot ";
    for ($f = 0; $f < $nf-1; $f ++) {
	$cmd   .= "'$outfile_ref->[$f]' using 3:4 title '' with lines ls $m, ";
    }
    $cmd   .= "'$outfile_ref->[$f]' using 3:4 title '' with lines ls $m";
    print GP "$cmd\n";


   
    close(GP);
    
    system("ps2pdf $psfile $pdfile\n");
    system("rm $psfile\n");
    
    if ($viewplot) {
	system("open $pdfile&\n"); 
    }
    
}

sub parse_powerfiles {
    my ($dir, $results_dir, $nf_ret, $ref_outfile) = @_;

    my $powerdir = "$dir/R-scape_join";
    my $powertag = "power";

    my $hlxdir   = "$dir/R-scape";
    my $hlxtag   = "helixcov";

    my @powerfile;
    my $prefix = "RF";
    my $suffix = $powertag;    
    FUNCS::sorted_files($powerdir, \@powerfile, $suffix, $prefix);
    my $nfp = $#powerfile+1;
    
    my @hlxfile;
    $prefix = "RF";
    $suffix = $hlxtag;    
    FUNCS::sorted_files($hlxdir, \@hlxfile, $suffix, $prefix);
    my $nfh = $#hlxfile+1;
    if ($nfh != $nfp) { print "nf power $nfp != $nfh nf hlx\n"; }
    $$nf_ret = $nfp;
    
    for (my $f = 0; $f < $nfp; $f ++) {
	
	my $cov_obs      = 0;
	my $cov_obs_frac = 0;
	my $cov_exp      = 0;
	my $cov_exp_frac = 0;
	my $hlx_cov      = 0;
	my $hlx_cov_frac = 0;

	my $name = "$powerfile[$f]";
	if ($name =~ /^(\S+).power$/) { $name = $1; }
	if ($name =~ /\/([^\/]+)$/)   { $name = $1; }
	
	my $nbp = 0;
	open(POWER, "$powerfile[$f]") || die; 
	while (<POWER>)
	{
	    if    (/^#\s+BPAIRS\s+(\d+)/)                  { $nbp = $1; }
	    elsif (/^# BPAIRS expected to covary\s+(\S+)/) { $cov_exp = $1; $cov_exp_frac = ($nbp > 0)? $1/$nbp : 0; }
	    elsif (/^# BPAIRS observed to covary\s+(\S+)/) { $cov_obs = $1; $cov_obs_frac = ($nbp > 0)? $1/$nbp : 0; }
	    
	}
	close(POWER);

	my $nhlx_cov = 0;
	my $nhlx_tot = 0;
	open(HLX, "$hlxfile[$f]")  || die;
	while (<HLX>)
	{
	    if    (/^# aggregated $aggtype\s+E-value: (\S+) P-value: \S+\s+\*\s*$/) { $nhlx_cov ++; $nhlx_tot ++; }
	    elsif (/^# aggregated $aggtype\s+E-value: (\S+) P-value: \S+\s*$/)      {               $nhlx_tot ++; }
	}
	close(HLX);

	$hlx_cov      = $nhlx_cov;
	$hlx_cov_frac = ($nhlx_tot > 0)? $nhlx_cov/$nhlx_tot : 0;
    

	$ref_outfile->[$f] = "$results_dir/$name.power";
	open(OUTF, ">$ref_outfile->[$f]") || die;
	print OUTF " $cov_obs $cov_exp  $cov_obs_frac $cov_exp_frac\n";
	print OUTF "-$hlx_cov $cov_exp -$hlx_cov_frac $cov_exp_frac\n";
	close(OUTF);
    }
    
}

   
```
