## Supplementary material for "RNA covariation at helix-level resolution for the identification of evolutionarily conserved RNA structure": profmark_sender.pl

```
#!/usr/bin/perl -w
# 
# profmark_sender
#
use strict;
use Class::Struct;
use POSIX;

#use lib '/Users/erivas/src/Mysrc/R-scape/scripts';
use lib '/n/eddy_lab/users/erivas/rscape_v2.0.0.p/scripts';

use FUNCS;

use vars qw ($opt_v $opt_R $opt_F);  # required if strict used
use Getopt::Std;
getopts ('vRF');

# Print a helpful message if the user provides no input file.
if (!@ARGV) {
    print "usage:  profmark_sender.pl [options] <Rfam> <maindir> <rscapedir> <shdir>\n\n";
    print "options:\n";
    print "-v    :  be verbose\n";
    exit;
}

my $Rfam        = shift;
my $maindir     = shift;
my $rscapedir   = shift;
my $shscript    = "$maindir/shscripts/profmark.sh";

my $do_rscape = 0;
if ($opt_R) { $do_rscape = 1; }

my $do_flanking = 0;
if ($opt_F) { $do_flanking = 1; }

my $suffix = "sto";
my $prefix = "";
my @family;
my $profmarkdir = "$maindir/data/$Rfam/profmark";
my $stodir      = "$profmarkdir/select";
FUNCS::sorted_files($stodir, \@family, $suffix, $prefix);

# clean up before we start
my $resultsdir  = "$profmarkdir/results";
system("rm $resultsdir/* \n");

my $nf = $#family + 1;

printf("NFAM $nf\n");

for (my $f = 0; $f < $nf; $f ++) {
    my $thisfam = "$family[$f]";
    if ($thisfam =~ /(RF\d+)/) { $thisfam = $1; }

    my $cmd = "sbatch $shscript $Rfam $maindir $rscapedir $thisfam $do_rscape";
    print "$f>$cmd\n\n";
    system("$cmd\n");
}
```
