## Supplementary material for "RNA covariation at helix-level resolution for the identification of evolutionarily conserved RNA structure": Rfam_select.pl

```
#!/usr/bin/perl -w
# 
# Select Rfam families
#
use strict;
use Class::Struct;
use POSIX;
    
use constant GNUPLOT => '/usr/local/bin/gnuplot';
use lib '/Users/erivas/src/Mysrc/R-scape/scripts';
use FUNCS;

use vars qw ($opt_v);  # required if strict used
use Getopt::Std;
getopts ('v');

# Print a helpful message if the user provides no input file.
if (!@ARGV) {
    print "usage:  Rfam_select.pl [options] <R-scape.seed> \n\n";
    print "options:\n";
    print "-v    :  be verbose\n";
    exit;
}

my $file     = shift;
my $listfile = "$file.select.list";
my $filedir  = $file;
if ($filedir =~ /^(\S+)\/\S+/) { $filedir = "$1/profmark"; }
print "dir $filedir\n";

my $NatMethfile = "/Users/erivas/writing/papers/R-scape/rscape-supplement/alignments/R-scape_TestSet/R-scape_Testset.list";

my $rscapedir = "/Users/erivas/src/Mysrc/R-scape";
my $rscape    = "$rscapedir/bin/R-scape";
my $easel     = "$rscapedir/lib/hmmer/easel/miniapps";

my $statsfile = "$file.stats";
my $cmd = "$easel/esl-alistat $file > $statsfile";
system("$cmd \n");
#system("more $statsfile");

my $nali = 0;
my @name;
my @id;
my @ac;
my @nsq;
my @alen;
parse_alistat($statsfile, \$nali, \@name, \@id, \@ac, \@nsq, \@alen);
print ("Nali = $nali\n");

# select
my $min_nsq = 40;
my $min_len = 0;
my $max_id  = 95;
select_fam($listfile, $max_id, $min_nsq, $min_len, $nali, \@name, \@id, \@ac, \@nsq, \@alen);


sub parse_alistat {
    my ($statsfile, $ret_nali, $name_ref, $id_ref, $ac_ref, $nsq_ref, $alen_ref) = @_;
    open (FILE, "$statsfile") || die;

    my $nali = 0;
    while (<FILE>) {
	
	if (/\/\//) {
	    $nali ++;
	}
	elsif (/Alignment name:\s+(\S+)\s*$/) {
	    $name_ref->[$nali] = $1;
	}
 	elsif (/Alignment length:\s+(\d+)\s*$/) {
	    $alen_ref->[$nali] = $1;
	}
 	elsif (/Average identity:\s+(\d+)\%\s*$/) {
	    $id_ref->[$nali] = $1;
	}
	elsif (/Number of sequences:\s+(\d+)\s*/) {
	    $nsq_ref->[$nali] = $1;
	}
 
    }
    close(FILE);

    if (0) {
	for (my $n = 0; $n < $nali; $n ++) {

	    $ac_ref->[$n] = get_AC($name_ref->[$n], $file);
	    printf "%d %s %s\n", $n, $name_ref->[$n], $ac_ref->[$n];
	}
    }
    
    $$ret_nali = $nali;
}

sub select_fam {
    my ($listfile, $maxid, $minnsq, $minlen, $nali, $name_ref, $id_ref, $ac_ref, $nsq_ref, $alen_ref) = @_;

    open (my $OUT, ">$listfile") || die;
    
    my @useme;
    for (my $n = 0; $n < $nali; $n ++) { $useme[$n] = 1; }

    # remove micro-RNAs
    #
    for (my $n = 0; $n < $nali; $n ++) {
	if ($name_ref->[$n] =~ /mir\-/) { $useme[$n] = 0; }  
	if ($name_ref->[$n] =~ /MIR/)   { $useme[$n] = 0; }  
    }
    print "remove mir- ";
    howmany ($nali, \@useme);
    
    # remove SNOR
    #
    for (my $n = 0; $n < $nali; $n ++) {
	if ($name_ref->[$n] =~ /SNOR/) { $useme[$n] = 0; }  
	if ($name_ref->[$n] =~ /sno/) { $useme[$n] = 0; }  
    }
    print "remove SNOR ";
    howmany ($nali, \@useme);
    
    # remove nsq < minnsq
    #
    for (my $n = 0; $n < $nali; $n ++) {
	if ($nsq_ref->[$n] < $minnsq) { $useme[$n] = 0; }  
    }
    print "remove nsq < $minnsq ";
    howmany ($nali, \@useme);
    
    # remove len < minlen
    #
    for (my $n = 0; $n < $nali; $n ++) {
	if ($alen_ref->[$n] < $minlen) { $useme[$n] = 0; }  
    }
    print "remove len < $minlen ";
    howmany ($nali, \@useme);
    
    # remove id > maxid
    #
    for (my $n = 0; $n < $nali; $n ++) {
	if ($id_ref->[$n] > $maxid) { $useme[$n] = 0; }  
    }
    print "remove id > $maxid ";
    howmany ($nali, \@useme);

    # write to file
    write_fam ($OUT, \@useme, $nali, $name_ref, $id_ref, $ac_ref, $nsq_ref, $alen_ref);

    close($OUT);
}


sub howmany {
    my ($n, $useme_ref) = @_;

    my $howmany = 0;
    for (my $x = 0; $x < $n; $x ++) { 
	if ($useme_ref->[$x]) { $howmany ++; } 
    }
    printf "howmany = %d / %d [remove %d]\n", $howmany, $n, $n-$howmany;
				      
}

sub write_fam {
    my ($OUT, $useme_ref, $n, $name_ref, $id_ref, $ac_ref, $nsq_ref, $alen_ref) = @_;

    my $u   = 0;
    my $old = 0;

    for (my $x = 0; $x < $n; $x ++) {
	if ($useme_ref->[$x]) {
	    $u ++;
	    $ac_ref->[$x] = get_AC($name_ref->[$x], $file);
	    
	    my $isnew = compareto($ac_ref->[$x], $NatMethfile);
	    if ($isnew == 0) { $old ++; }

	    printf      "%d> %s %d %d %d %d %s | new? %d\n", $u, $ac_ref->[$x], $id_ref->[$x], $nsq_ref->[$x], $alen_ref->[$x], $id_ref->[$x], $name_ref->[$x], $isnew;
	    printf $OUT "%d> %s %d %d %d %d %s\n", $u, $ac_ref->[$x], $id_ref->[$x], $nsq_ref->[$x], $alen_ref->[$x], $id_ref->[$x], $name_ref->[$x];

	    # extract alignment
	    get_sto($ac_ref->[$x], $file);

	    # run R-scape to generate null alignments
	    run_rscape($ac_ref->[$x]);
	}
    }

    print "old $old\n";
}

sub get_AC {
    my ($name, $stofile) = @_;

    if (!$name) { return ; }
    
    my $AC = "";

    my $thisname;
    my $thisac;
	
    open(F, "$stofile") || die;
    while(<F>) {
	if    (/^#=GF AC\s+(\S+)\s*$/) { $thisac = $1; }
	elsif (/^#=GF ID\s+(\S+)\s*$/) { 
	    $thisname = $1;
	    if ($name =~ /^$thisname$/) { return $thisac; }
	}
	
    }
    close(F);

    return $AC;
}

sub compareto {
    my ($ac, $file) = @_;

    my $isnew = 1;

    open(F, "$file") || die;
    while(<F>) {
	if (/(RF\d+)/) {
	    my $rf = $1;
	    if ($rf =~ /^$ac$/) {		
		$isnew = 0; 
		return $isnew; 
	    }
	}
    }
    close(F);

    return $isnew;
}

sub get_sto {
    my ($ac, $file) = @_;

    my $stofile = "$filedir/select/$ac.seed.sto";
    my $cmd = "$easel/esl-afetch $file $ac > $stofile";
    print "$cmd\n";
    system("$cmd \n");

}


sub run_rscape {
    my ($ac) = @_;
    
    my $stoname = "$ac.seed.sto";
    my $stofile = "$filedir/select/$stoname";
    my $outdir  = "$filedir/select_R-scape_null";
    my $cmd = "$rscape --outnull --cacofold --nofigures --outdir $outdir $stofile > $outdir/$stoname.rscape";
    print "$cmd\n";
    system("$cmd \n");
}
```
