## Supplementary figures and images for "RNA covariation at helix-level resolution for the identification of evolutionarily conserved RNA structure"

### AntisenseAlu_alignment_1.R2R.sto.pdf

AntisenseAlu_alignment_1

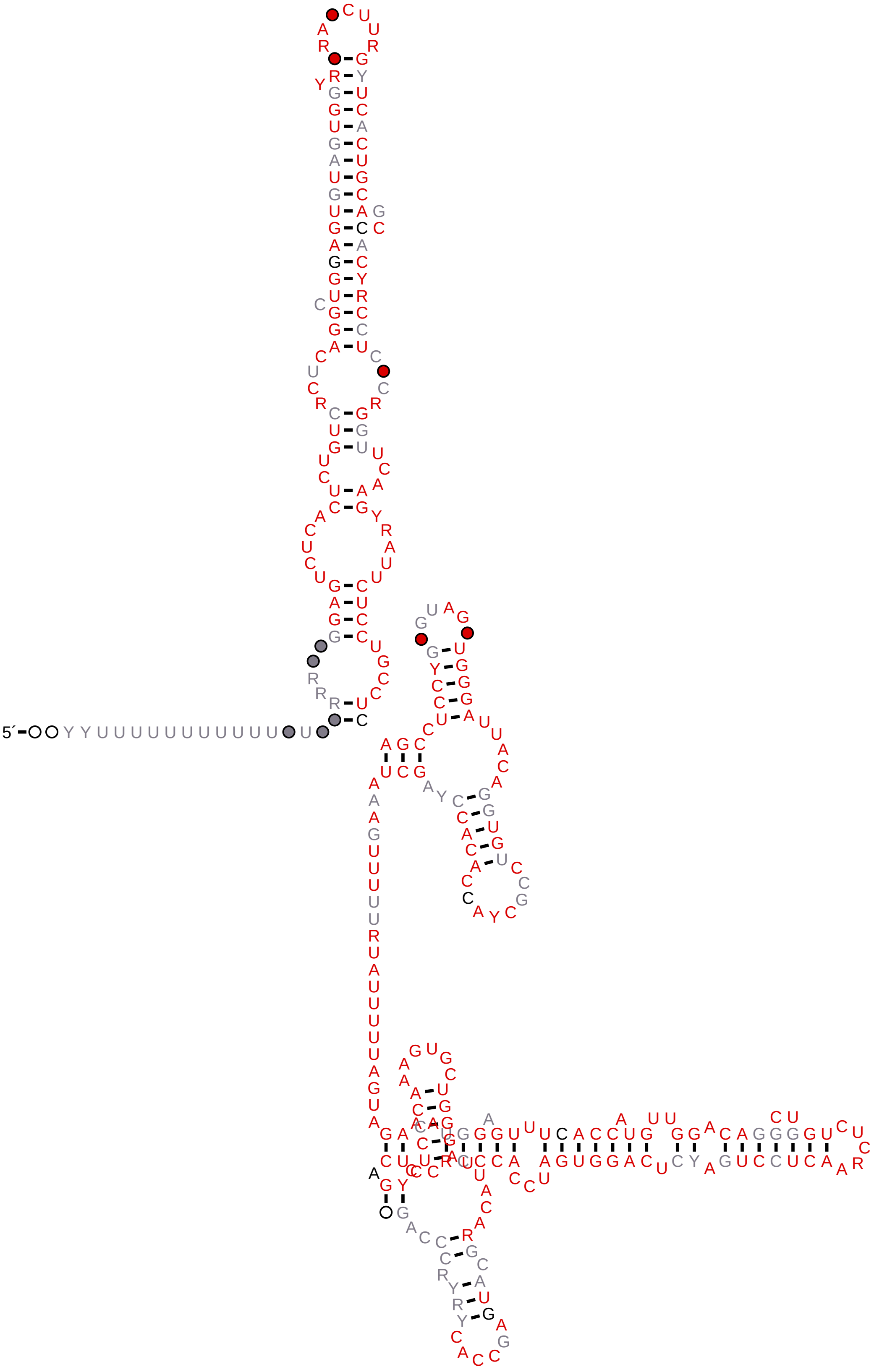

### Figure_1.pdf

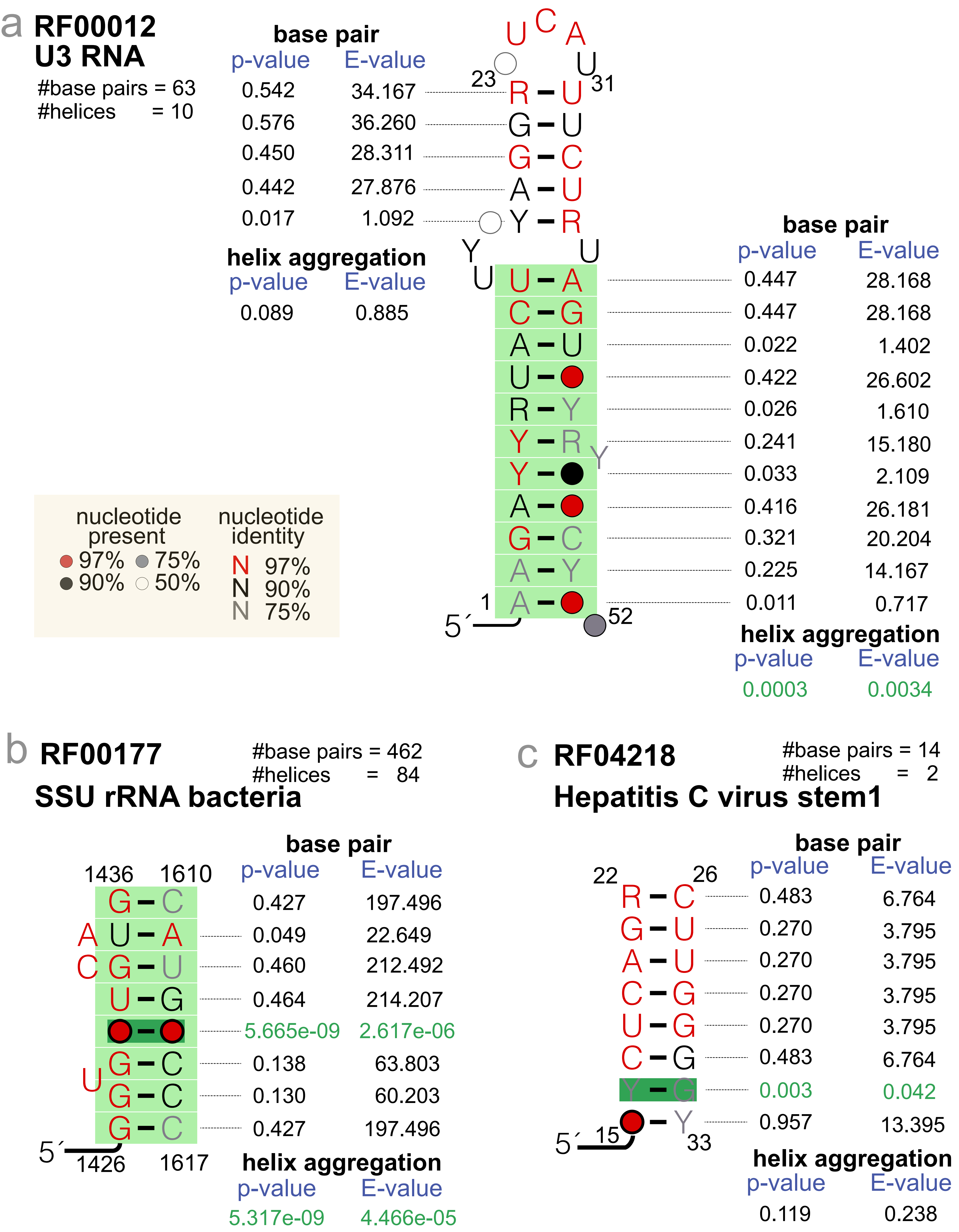

### Figure_2.pdf

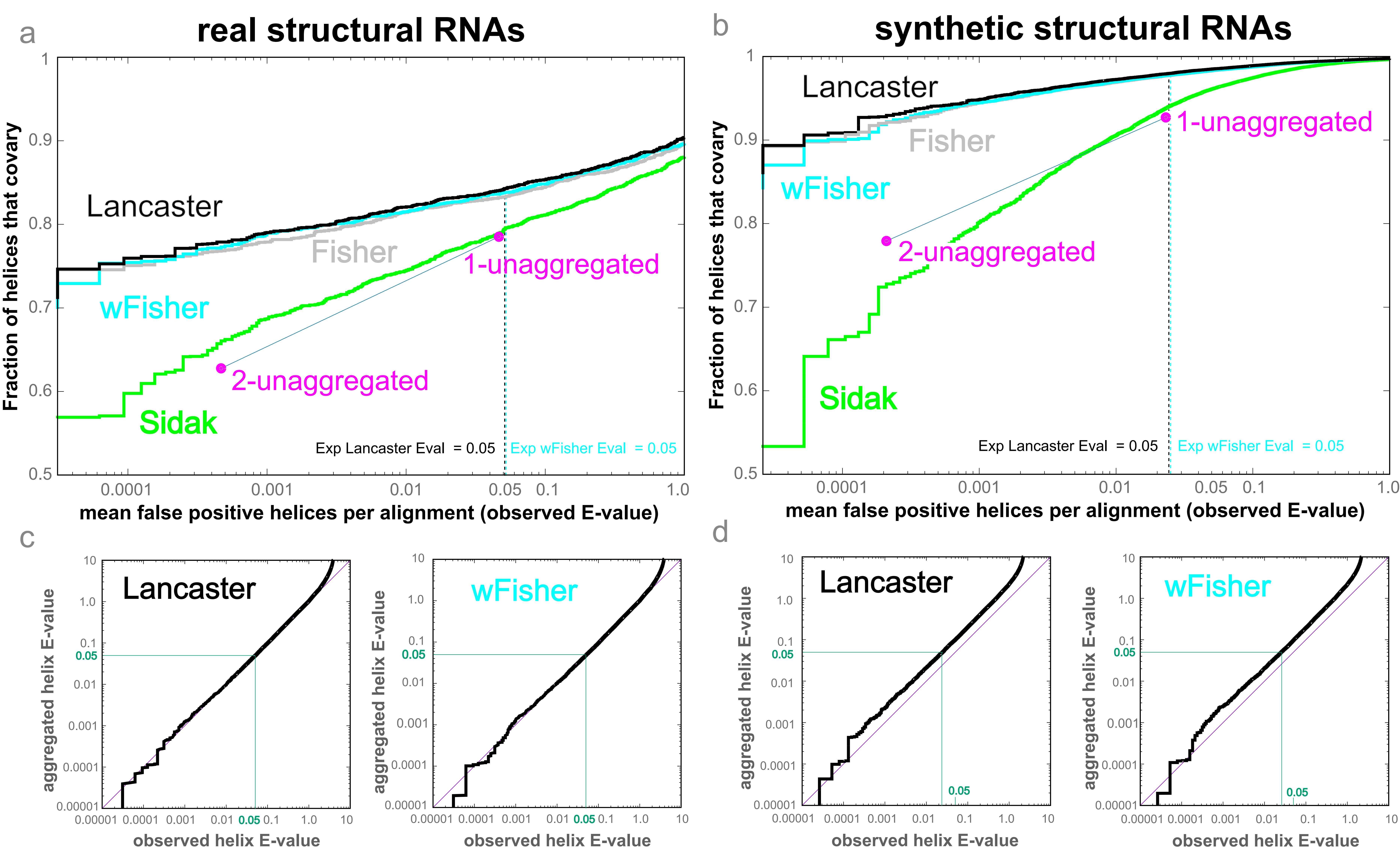

### Figure_S1.bck.pdf

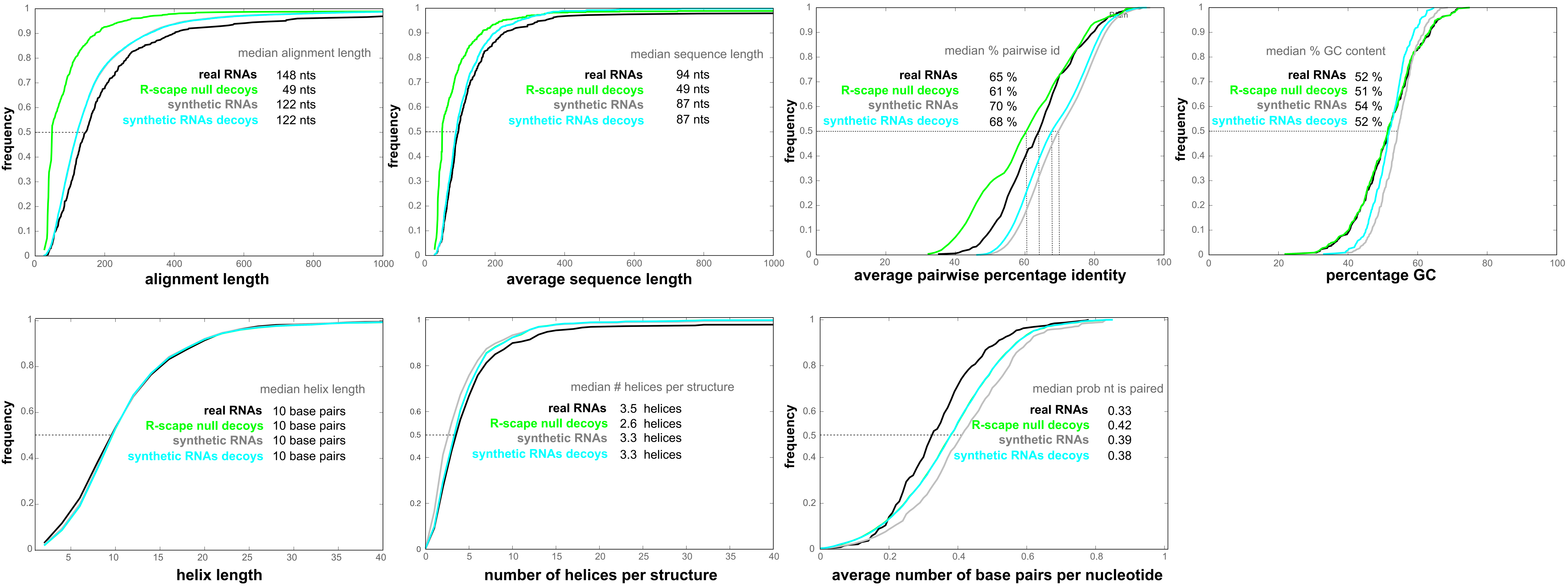

### Figure_S1.pdf

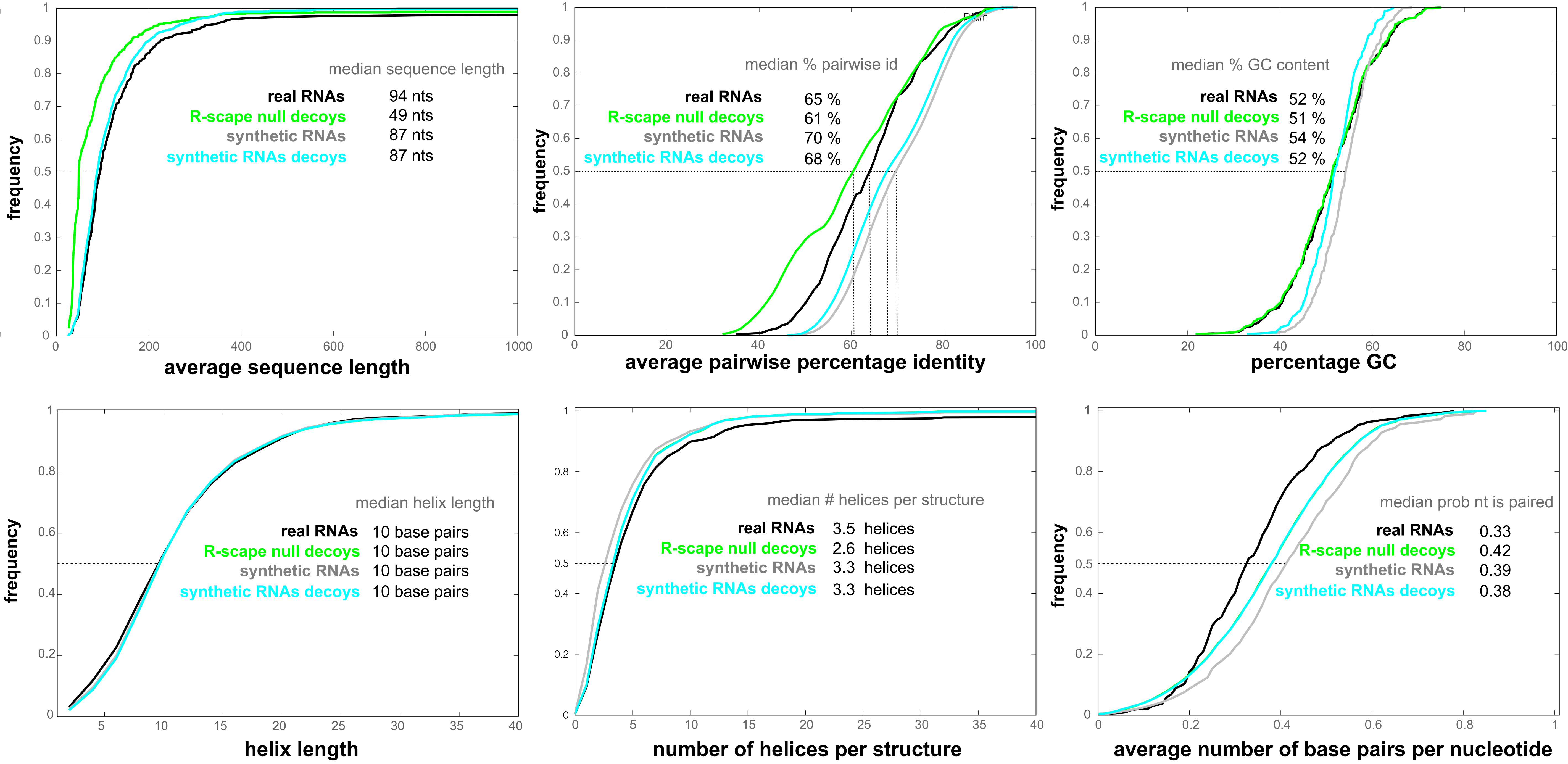

### Figure_S2.pdf

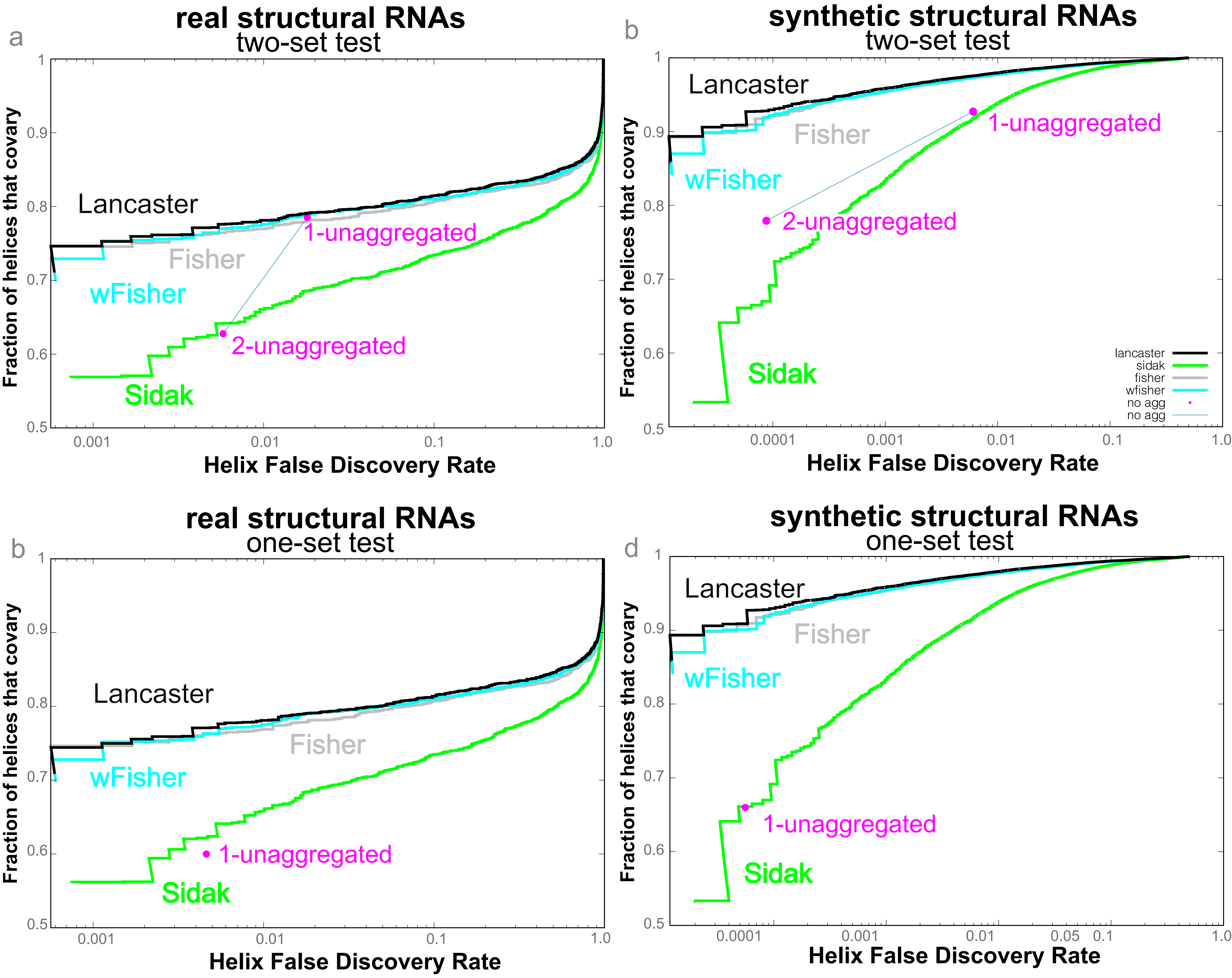

### Figure_S3.pdf

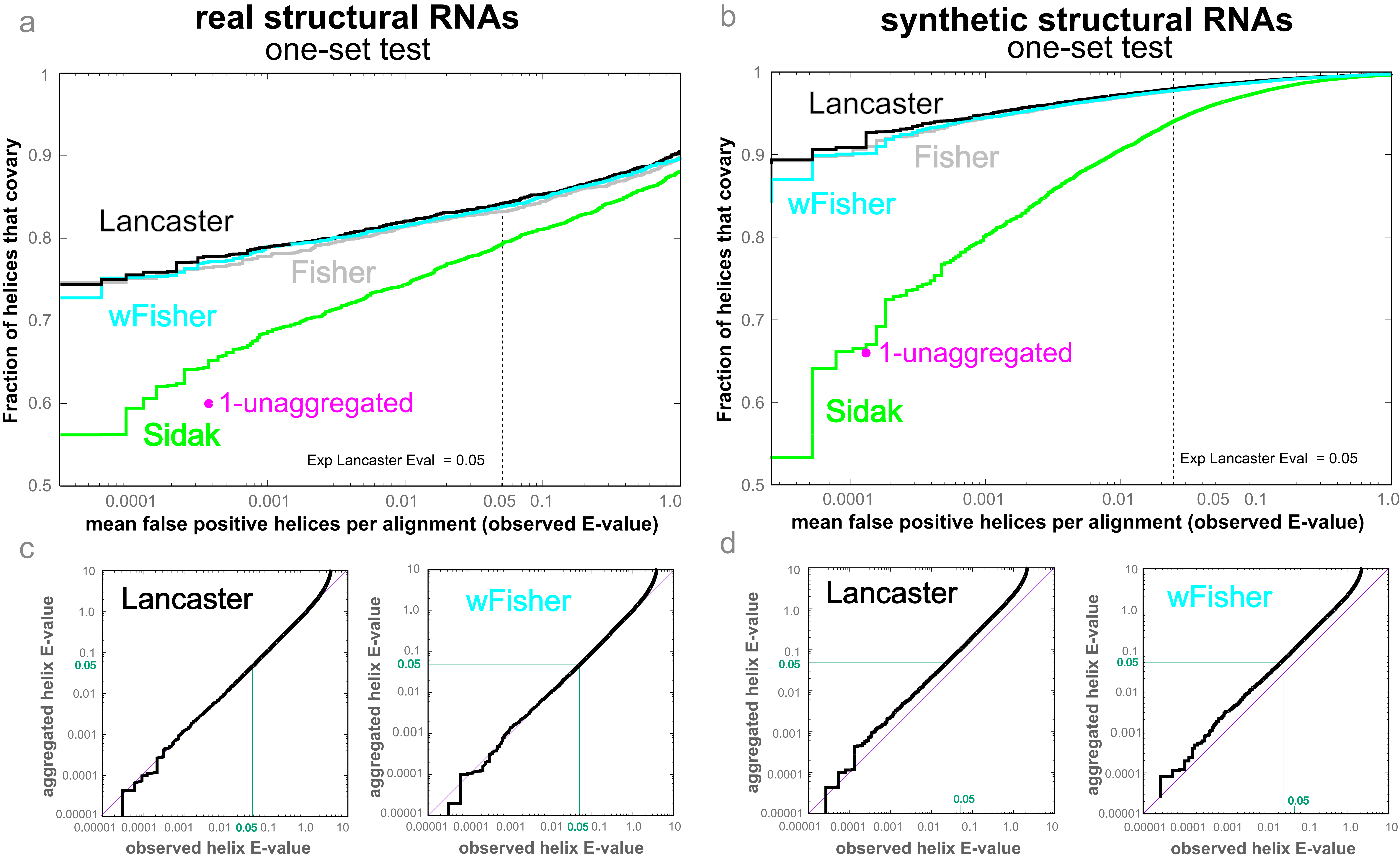

### Figure_S4.pdf

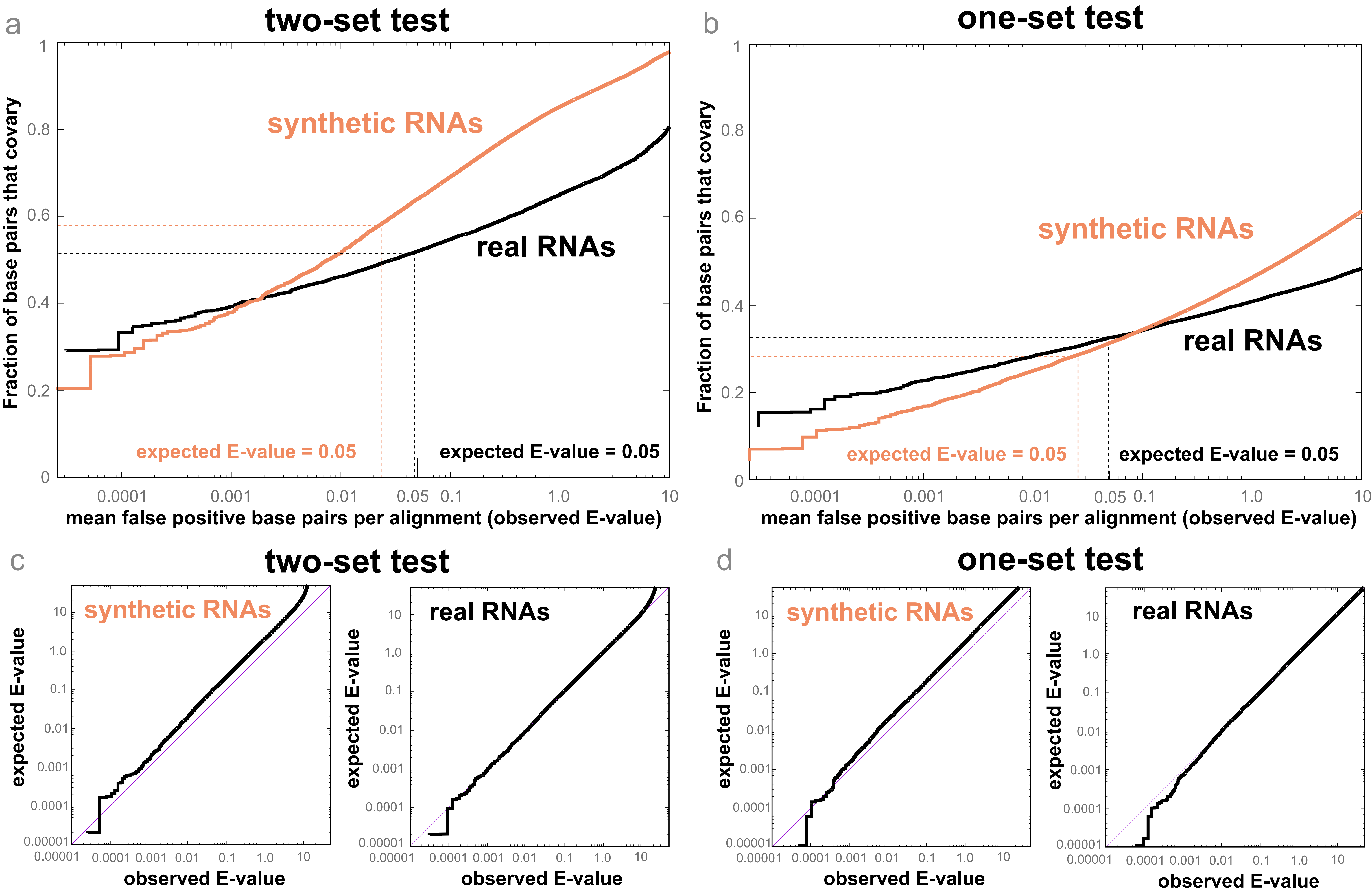

### Figure_S5.pdf

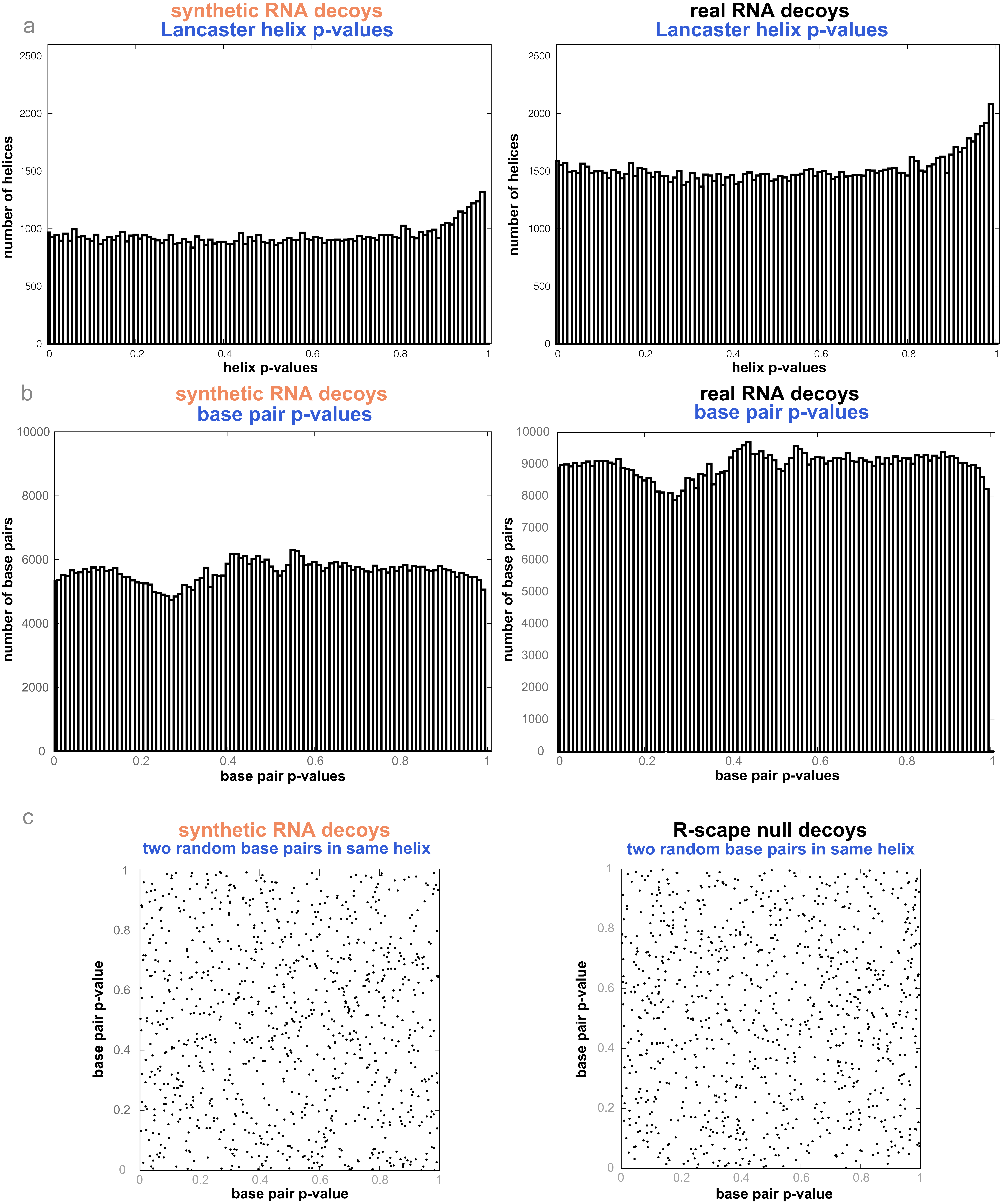

### HOTAIR_D1.human.infernal.cmalign_1.R2R.sto.pdf

HOTAIR_D1.human.infernal.cmalign_1

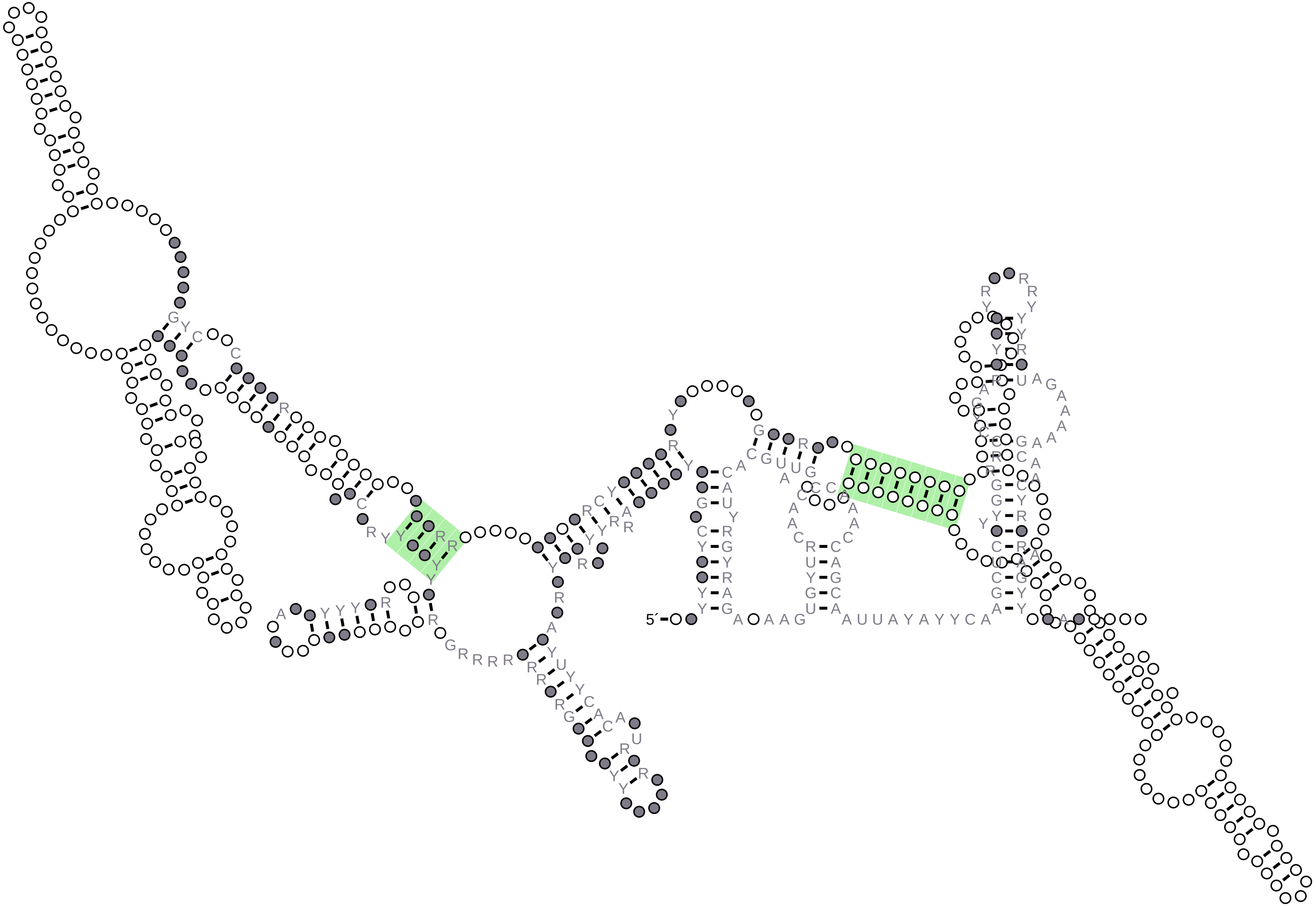

### HOTAIR_D1_1.R2R.sto.pdf

#### HOTAIR_D1_1

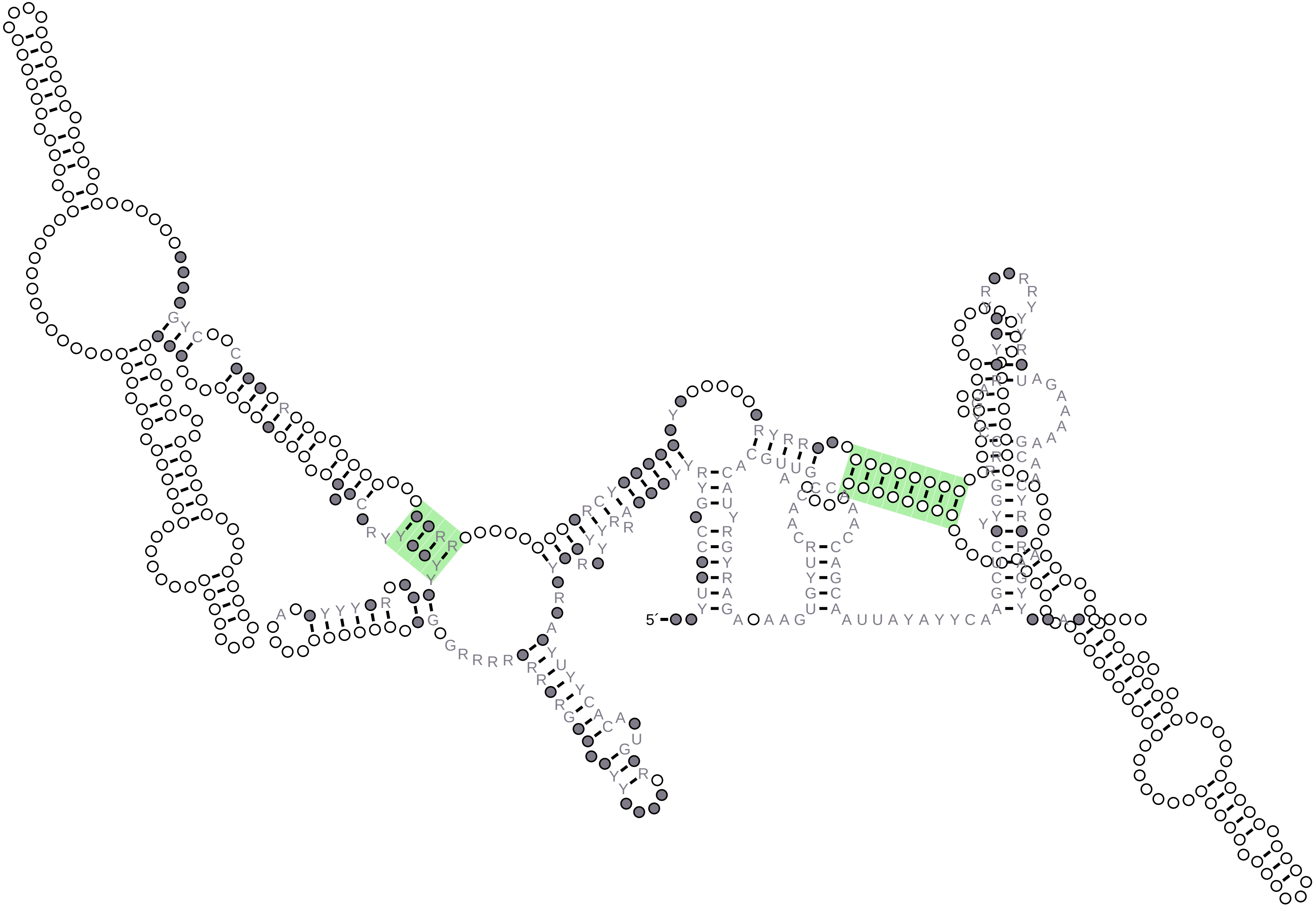

### HOTAIR_D2.human.infernal.cmalign_1.R2R.sto.pdf

HOTAIR_D2.human.infernal.cmalign_1

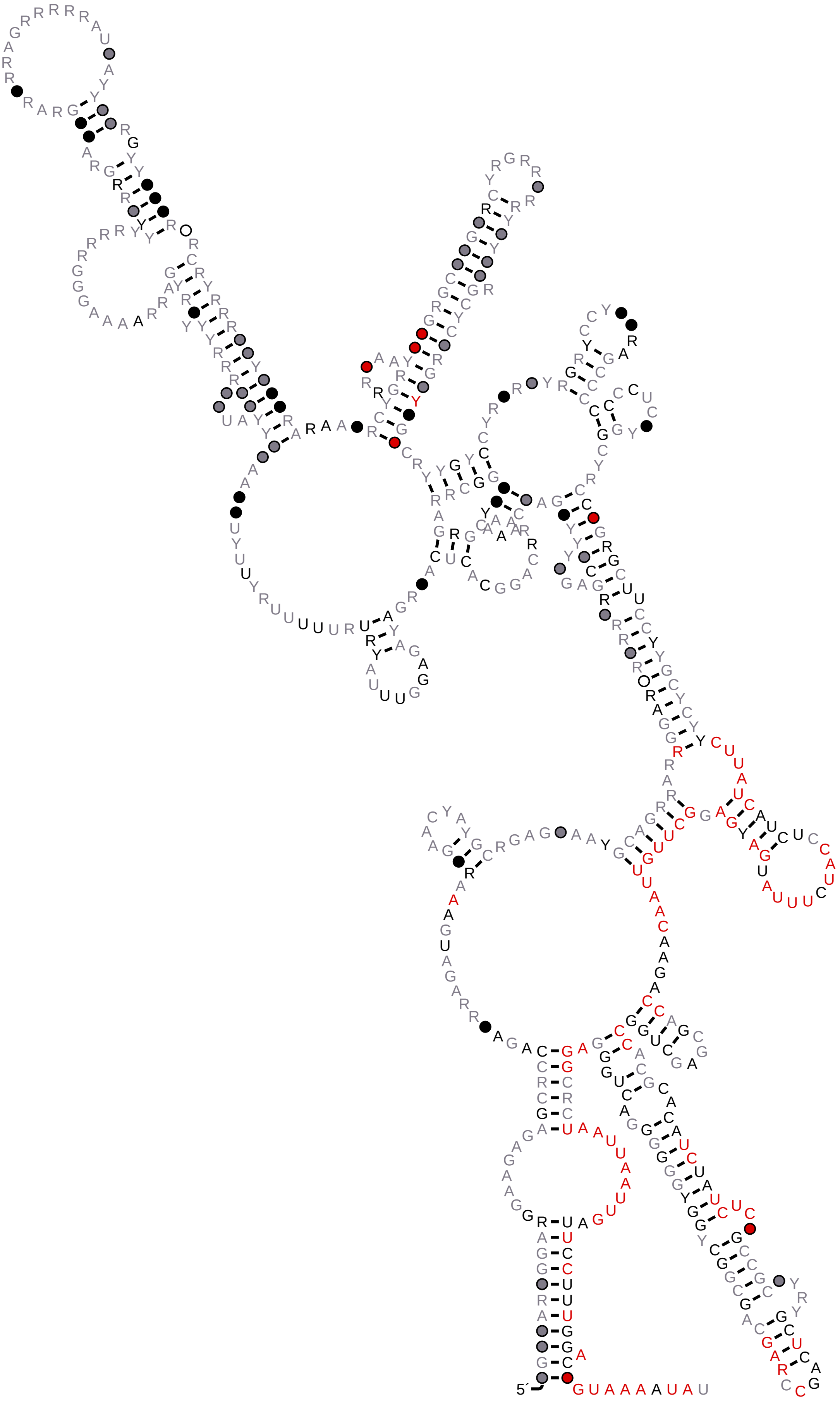

### HOTAIR_D2_1.R2R.sto.pdf

HOTAIR_D2_1

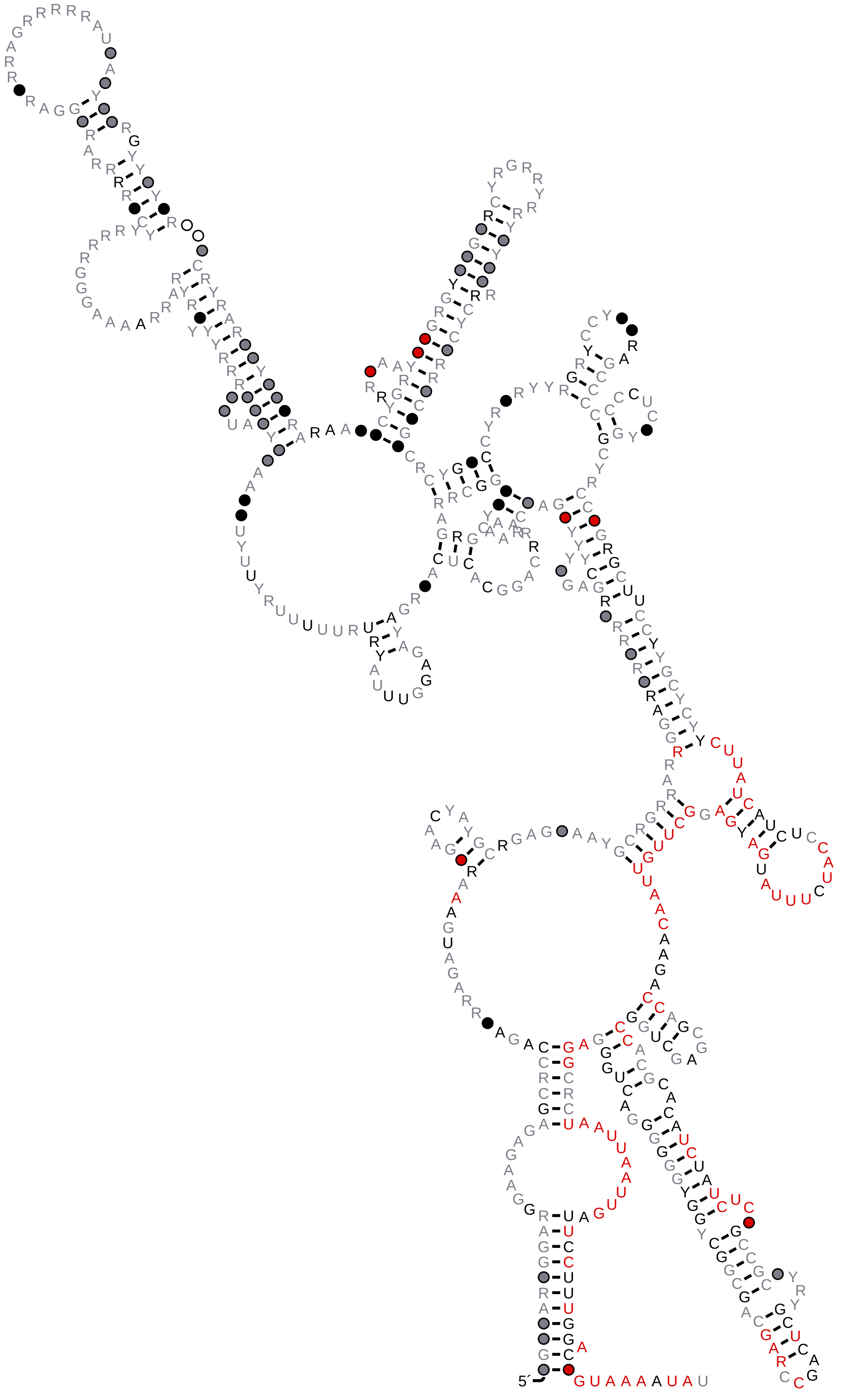

### HOTAIR_D3.human.infernal.cmalign_1.R2R.sto.pdf

HOTAIR_D3.human.infernal.cmalign_1

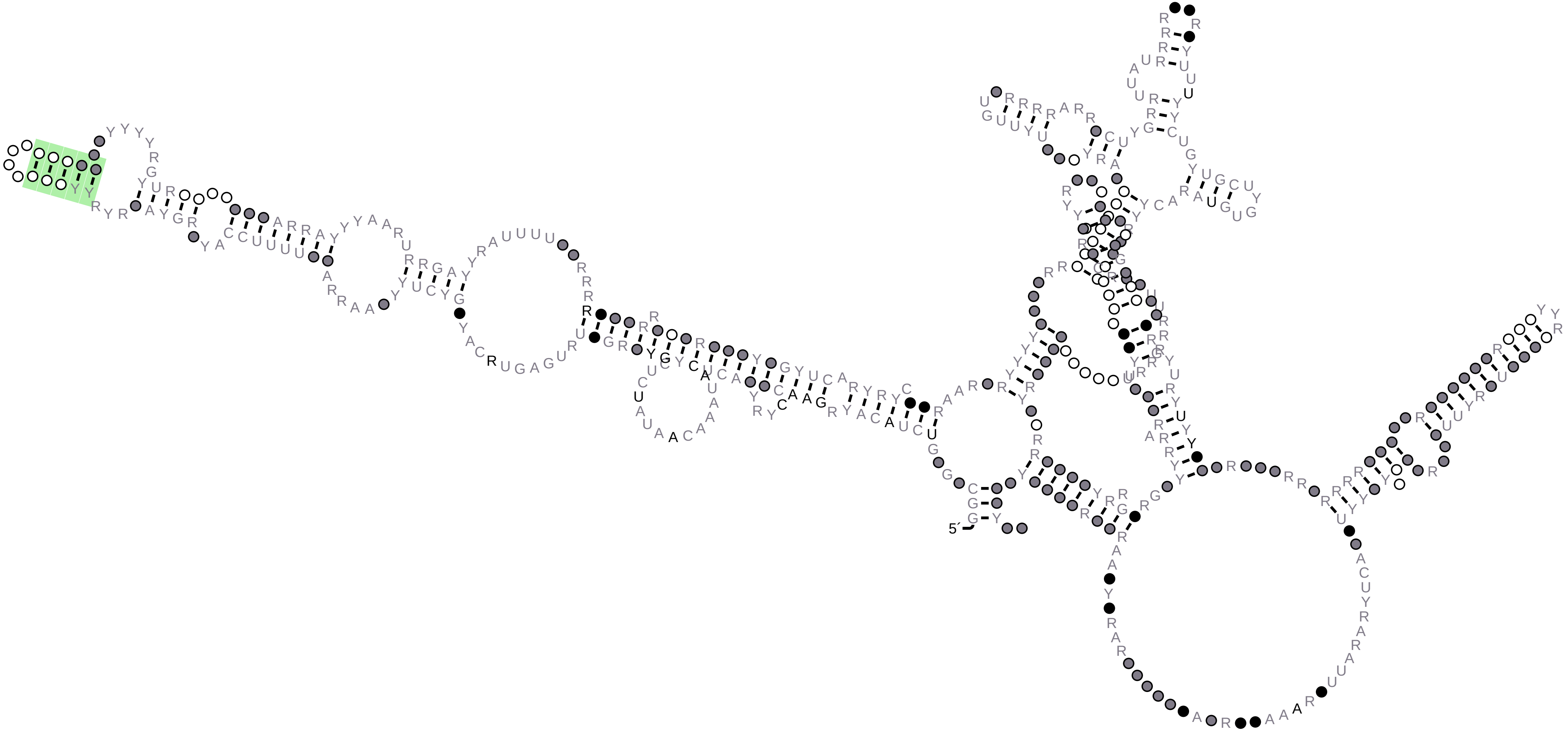

### HOTAIR_D3_1.R2R.sto.pdf

HOTAIR_D3_1

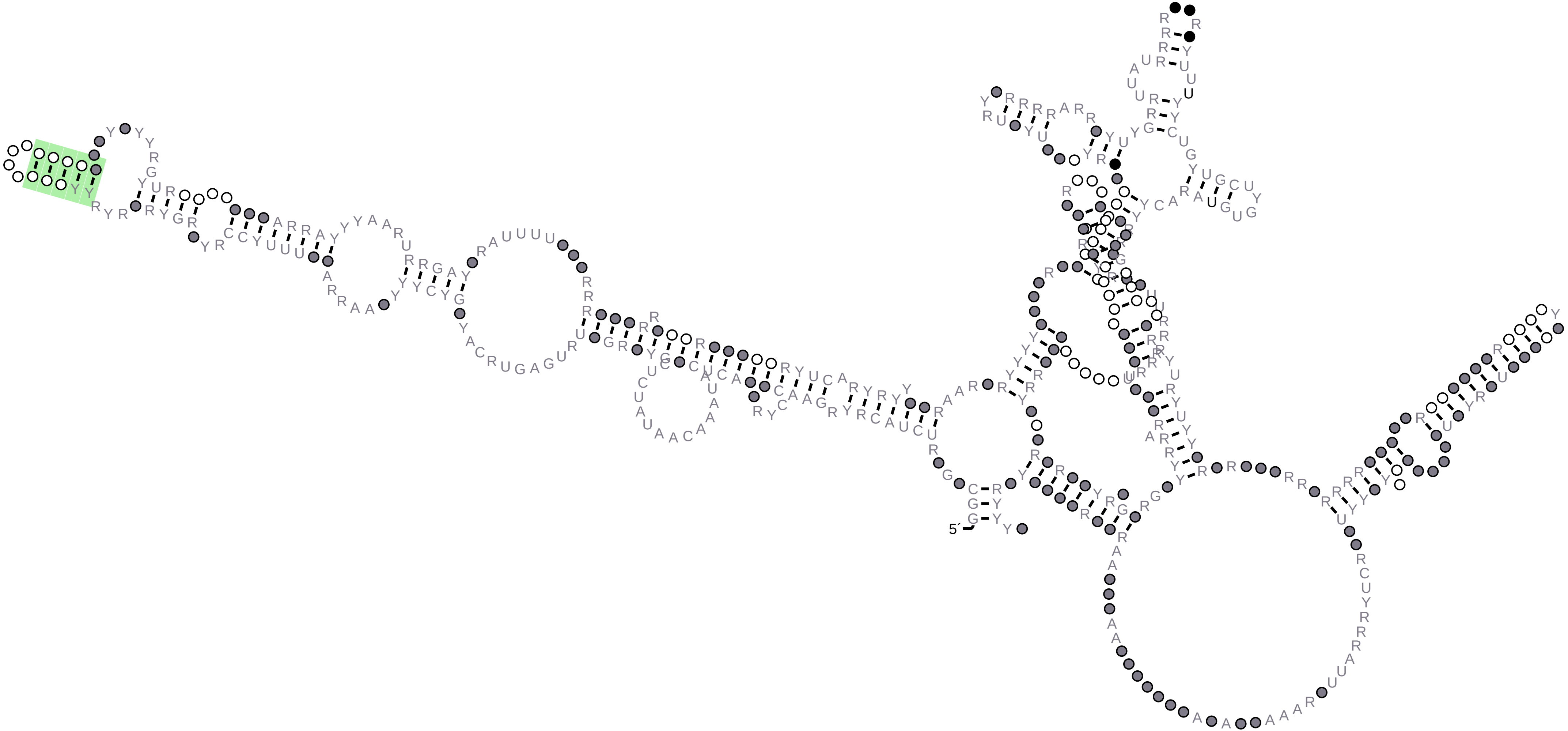

### HOTAIR_D4.human.infernal.cmalign_1.R2R.sto.pdf

#### HOTAIR_D4.human.infernal.cmalign_1

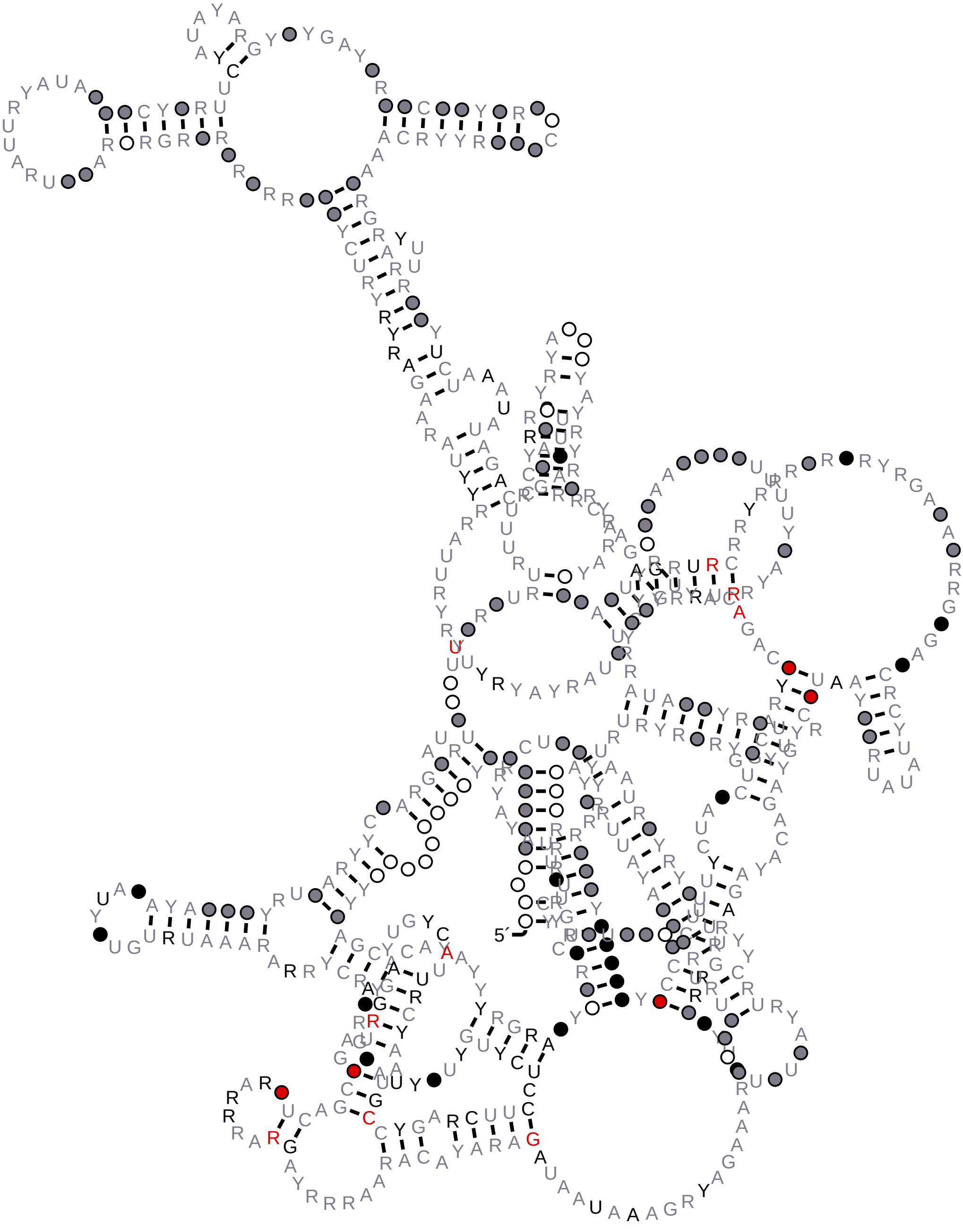

### HOTAIR_D4_1.R2R.sto.pdf

HOTAIR_D4_1

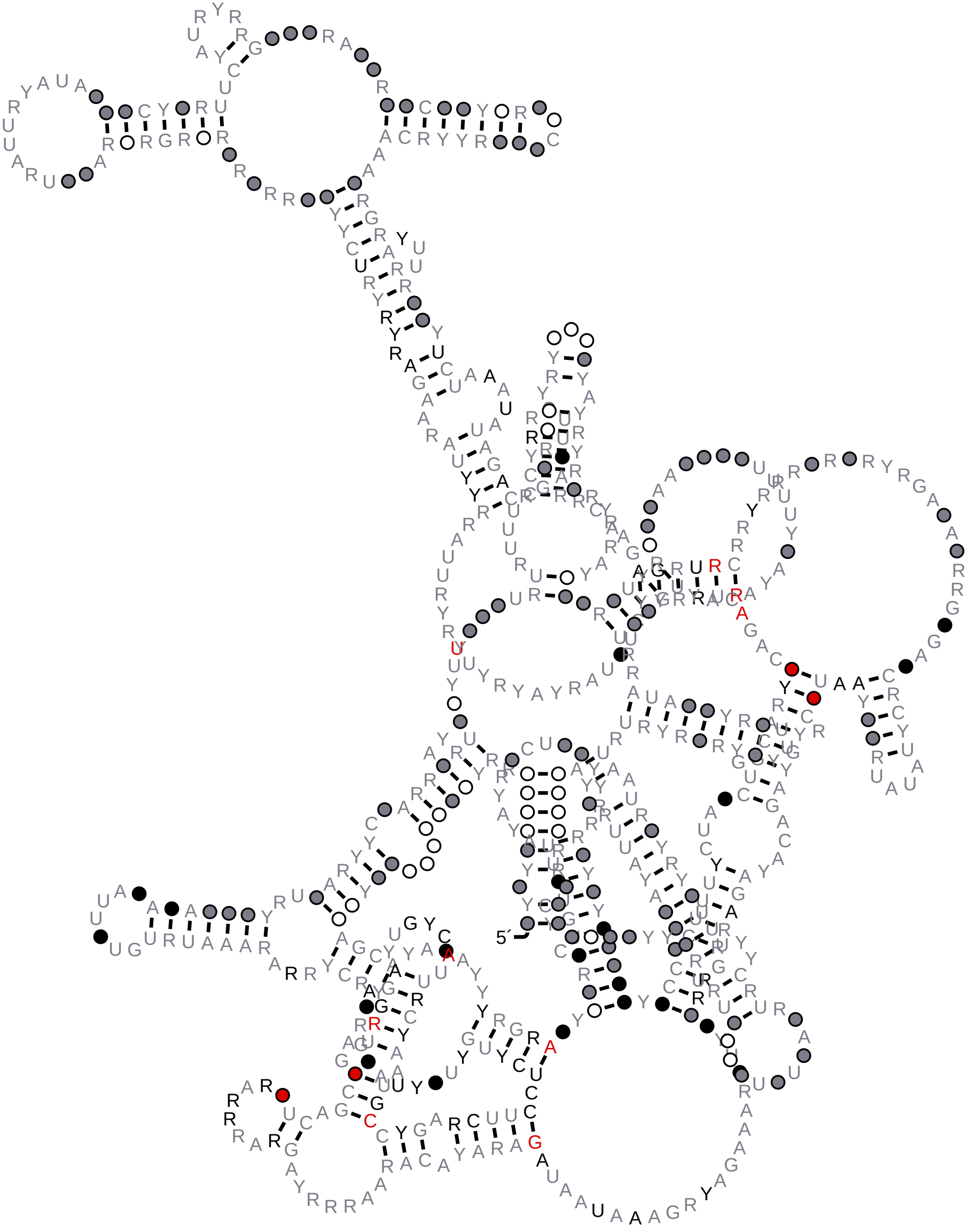

### NEAT1_21-322_cmalign_reformat.muscle.sto.withss.R2R.sto.pdf

#### NEAT1_21-322_cmalign_reformat.muscle.sto.withss

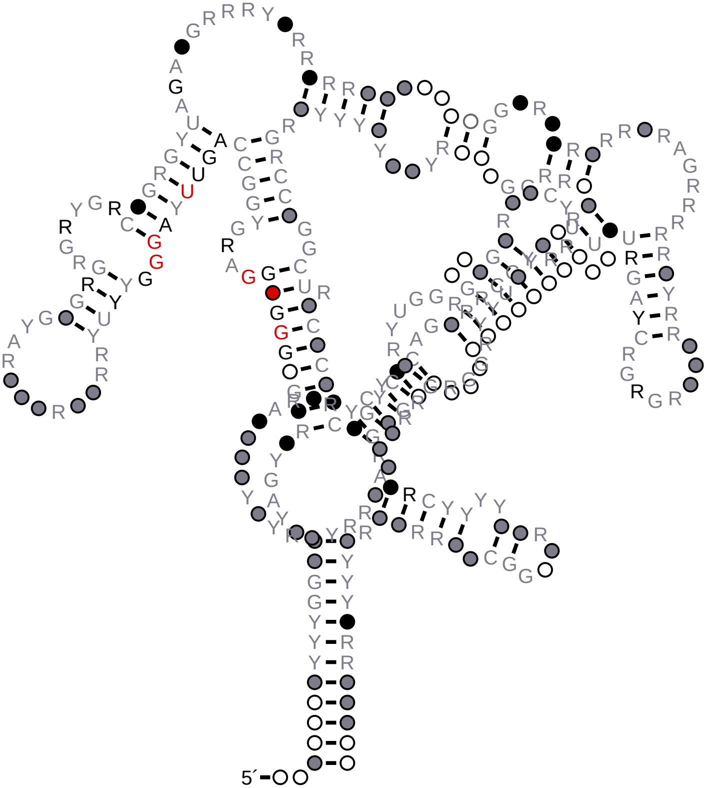

### NEAT1_323-501_cmalign_reformat.human_1.R2R.sto.pdf

#### NEAT1_323-501_cmalign_reformat.human_1

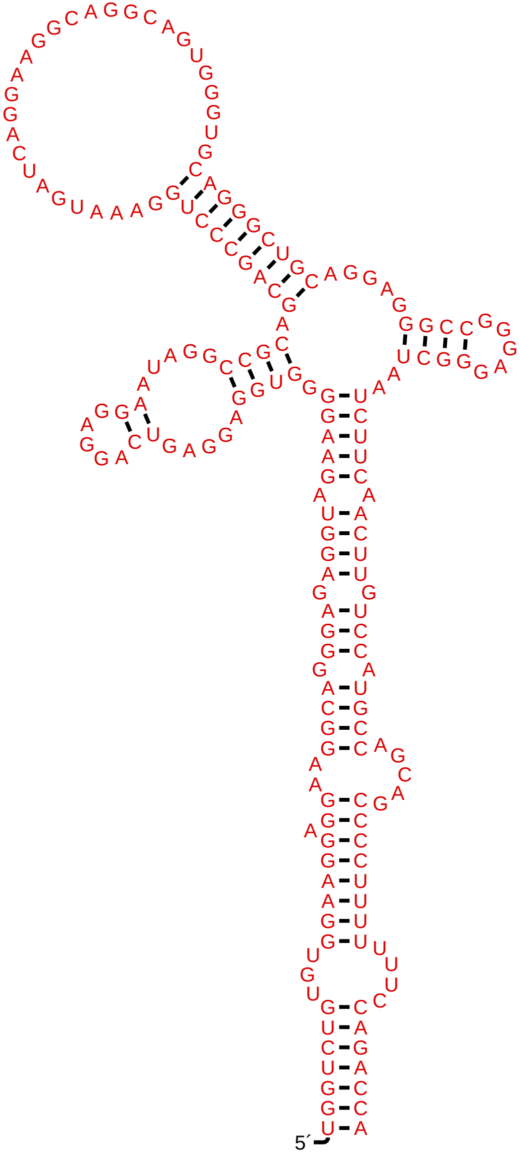

### NEAT1_323-501_cmalign_reformat_1.R2R.sto.pdf

#### NEAT1_323-501_cmalign_reformat_1

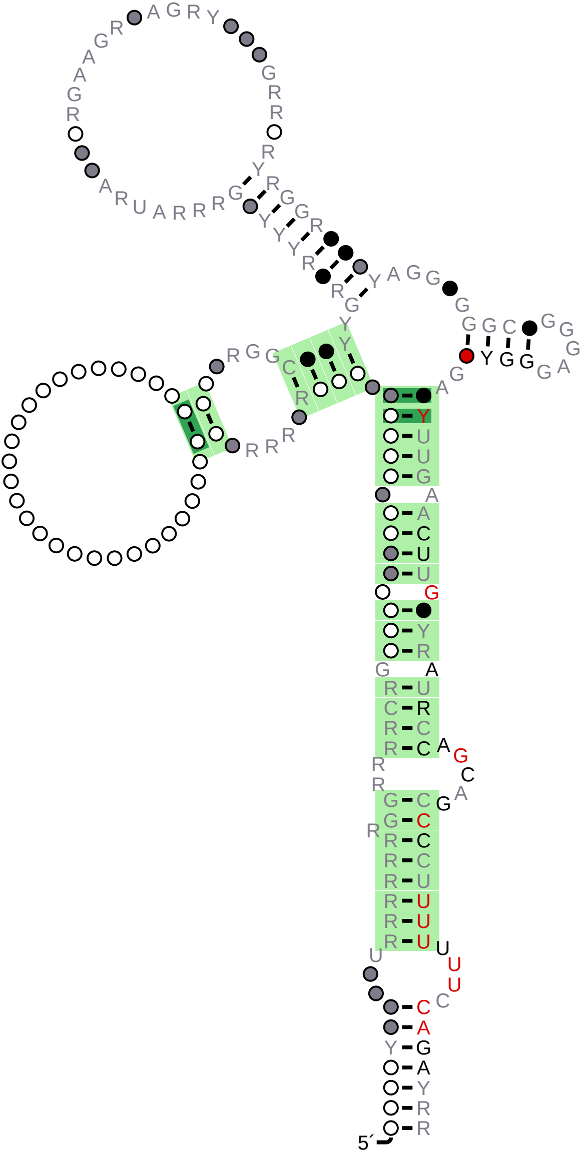

### NEAT1_1037-1268_cmalign_reformat.muscle.sto.withss.R2R.sto.pdf

NEAT1_1037-1268_cmalign_reformat.muscle.sto.withss

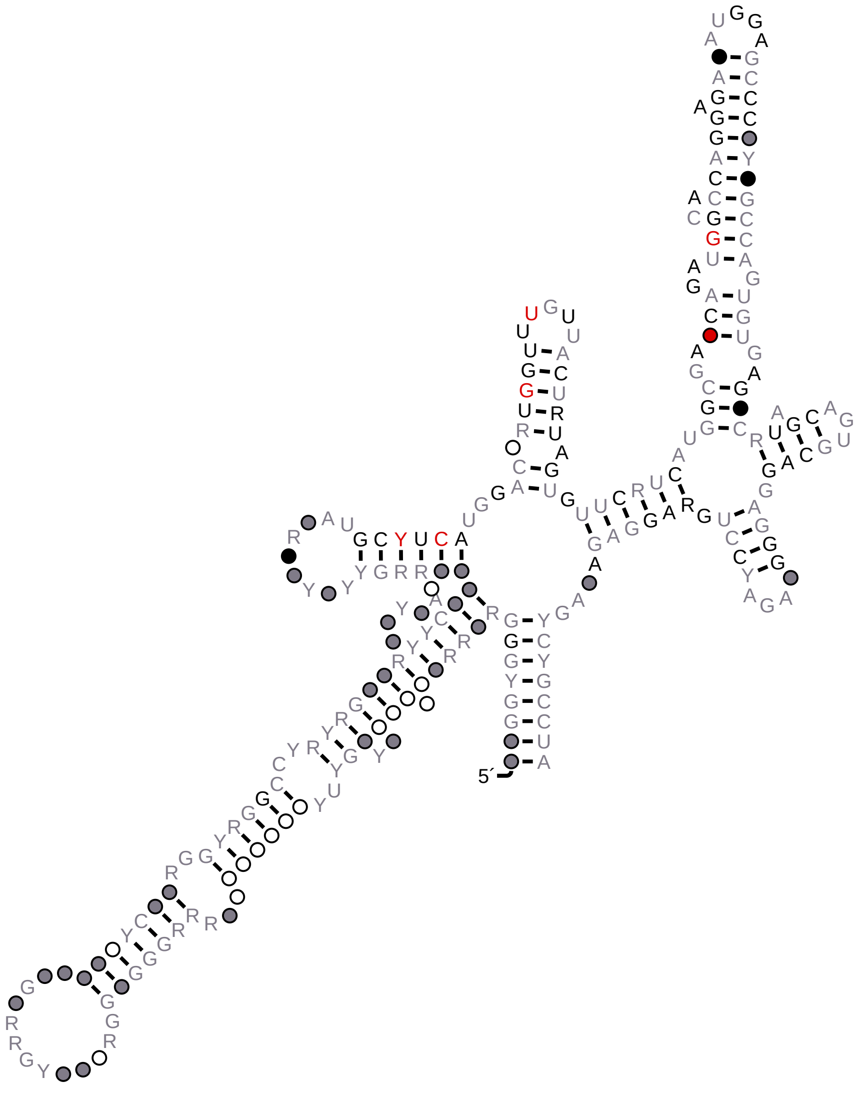

### NEAT1_1269-1467_cmalign_reformat.muscle.sto.withss.R2R.sto.pdf

#### NEAT1_1269-1467_cmalign_reformat.muscle.sto.withss

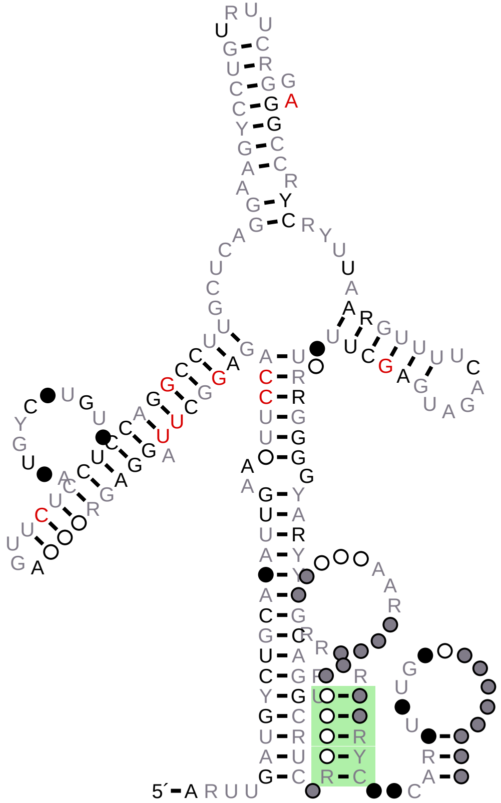

### SenseAlu_alignment_1.R2R.sto.pdf

SenseAlu_alignment_1

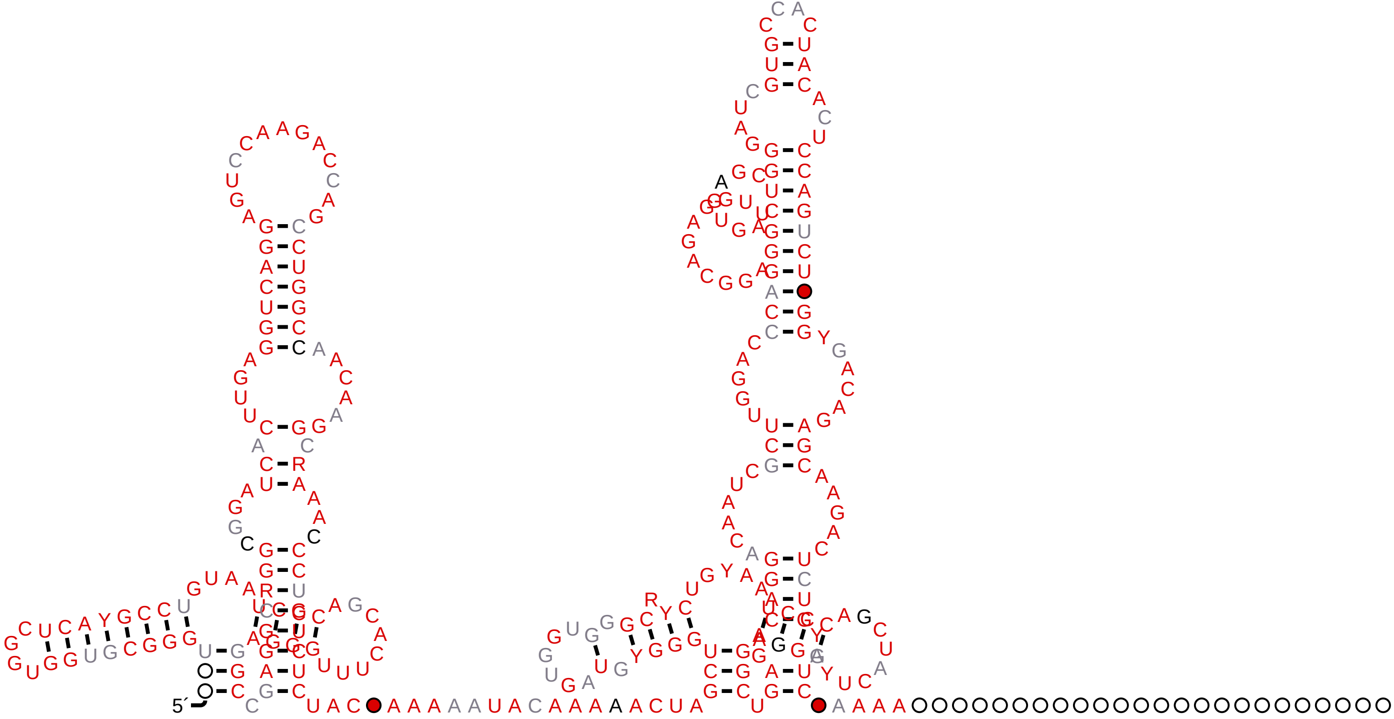
